# Harmonized Protocol for Subfield Segmentation in the Hippocampal Body on High-Resolution *in vivo* MRI from the Hippocampal Subfields Group (HSG)

**DOI:** 10.1101/2025.04.29.651039

**Authors:** Ana M. Daugherty, Valerie Carr, Kelsey L. Canada, Gustaf Rådman, Thackery Brown, Jean Augustinack, Katrin Amunts, Arnold Bakker, David Berron, Alison Burggren, Gael Chetelat, Robin de Flores, Song-Lin Ding, Yushan Huang, Ricardo Insausti, Elliott Johnson, Prabesh Kanel, Attila Keresztes, Olga Kedo, Kristen M. Kennedy, Joshua Lee, Nikolai Malykhin, Anjelica Martinez, Susanne Mueller, Elizabeth Mulligan, Noa Ofen, Daniela Palombo, Lorenzo Pasquini, John Pluta, Naftali Raz, Tracy Riggins, Karen M. Rodrigue, Samaah Saifullah, Margaret L. Schlichting, Craig Stark, Lei Wang, Paul Yushkevich, Renaud La Joie, Laura Wisse, Rosanna Olsen, the Alzheimer’s Disease Neuroimaging Initiative the Hippocampal Subfields Group

**Author notes:** A.M. Daugherty and V. Carr should be considered joint first author. R. La Joie, L. Wisse and R. Olsen should be considered joint senior author. R. Insausti passed away before the publication of this manuscript; lead and senior authors verify his valuable contribution to the work reported herein. Data used in preparation of this article were obtained from the Alzheimer’s Disease Neuroimaging Initiative (ADNI) database (adni.loni.usc.edu). As such, the investigators within the ADNI contributed to the design and implementation of ADNI and/or provided data but did not participate in analysis or writing of this report. A complete listing of ADNI investigators can be found at: http://adni.loni.usc.edu/wp-content/uploads/how_to_apply/ADNI_Acknowledgement_List.pdf.

## Abstract

Hippocampal subfields differentially develop and age, and they vary in vulnerability to neurodegenerative diseases. Innovation in high-resolution imaging has accelerated clinical research on human hippocampal subfields, but substantial differences in segmentation protocols impede comparisons of results across laboratories. The Hippocampal Subfields Group (HSG) is an international organization seeking to address this issue by developing a histologically-valid, reliable, and freely available segmentation protocol for high-resolution T_2_-weighted 3 tesla MRI (http://www.hippocampalsubfields.com). Here, we report the first portion of the protocol focused on subfields in the hippocampal body; protocols for the head and tail are in development. The body protocol includes definitions of the internal boundaries between subiculum, Cornu Ammonis (CA) 1-3 subfields, and dentate gyrus, in addition to the external boundaries of the hippocampus apart from surrounding white matter and cerebrospinal fluid. The segmentation protocol is based on a novel histological reference data set labeled by multiple expert neuroanatomists. With broad participation of the research community, we voted on the segmentation protocol via online survey, which included detailed protocol information, feasibility testing, demonstration videos, example segmentations, and labeled histology. All boundary definitions were rated as having high clarity and reached consensus agreement by Delphi procedure. The harmonized body protocol yielded high inter- and intra-rater reliability. In the present paper we report the procedures to develop and test the protocol, as well as the detailed procedures for manual segmentation using the harmonized protocol. The harmonized protocol will significantly facilitate cross-study comparisons and provide increased insight into the structure and function of hippocampal subfields across the lifespan and in neurodegenerative diseases.

## 1.0 Introduction

The hippocampus is one of the most prolifically studied brain regions indexed in PubMed (Simpson et al., 2021). Hippocampal structure and function measured from MRI are acknowledged correlates of learning and memory (Squire, 2004); its morphometric shape and volume dynamically change across childhood development and aging (Botdorf et al., 2022; Bussy et al., 2021; Langnes et al., 2020); and it is vulnerable to multiple pathophysiological, genetic and environmental factors (Walhovd et al., 2023). Hippocampal volume is of particular importance in the assessment, diagnosis and progression of Alzheimer’s disease and related dementia (Dubois et al., 2007; Frisoni & Jack, 2011; Sperling et al., 2011). However, the hippocampus is not a unitary structure: it is composed of subfields that have distinct cytoarchitecture, vascularization, gene expression, functional connectivity, and vulnerabilities to pathology (Braak & Braak, 1991; Duvernoy et al., 2013; Insausti & Amaral, 2004; Small et al., 2011). The human hippocampal subfields include subiculum complex (Sub), dentate gyrus (DG), and Cornu Ammonis (CA) sectors 1-3; and some neuroanatomists discern a CA4 region (Ding, 2013; Duvernoy et al., 2013; Palomero-Gallagher et al., 2020) whereas others consider it the hilus region of the DG (Insausti & Amaral, 2004). Identification and measurement of hippocampal subfields as distinct structures can improve specificity of functional correlates and early detection of diseases across the lifespan (e.g., La Joie et al., 2013; Mueller et al., 2010; Riphagen et al., 2020; Wisse et al., 2014).

Since the introduction of high-resolution T_2_-weighted *in vivo* imaging methods nearly two decades ago (e.g., Mueller & Weiner, 2009; Zeineh et al., 2000), the literature has grown exponentially, accompanied by a multitude of protocols to label human hippocampal subfields on MRI (Wisse et al., 2017). These protocols differ in anatomical nomenclature and segmentation definitions, resulting in barriers to synthesis and interpretation of the combined literature (Yushkevich et al., 2015) that slow scientific progress and clinical translation.

The Hippocampal Subfields Group (HSG) formed to address these barriers through the development of a harmonized protocol for segmenting hippocampal subfields that can be applied to samples across the lifespan and of different disease pathology. To date, the HSG has approximately 200 active members from 33 countries that support a distributed working group structure to develop a histologically-valid and reliable segmentation protocol for human hippocampal subfields (see Figure 1). Our previous publications described a comparison of 21 protocols to determine the scope of the disagreement in labels (Yushkevich et al., 2015), an overview of the HSG purpose and structure (Wisse et al., 2017), and an intermediate progress update on portions of the protocol development for the hippocampal body (Olsen et al., 2019). The current paper describes the procedures to develop the harmonized protocol to label subfields in the hippocampal body, and the supporting evidence for validation. At the end of the report, we include a summary of the protocol in the hippocampal body that is ready for application in the field.

**Figure 1.**
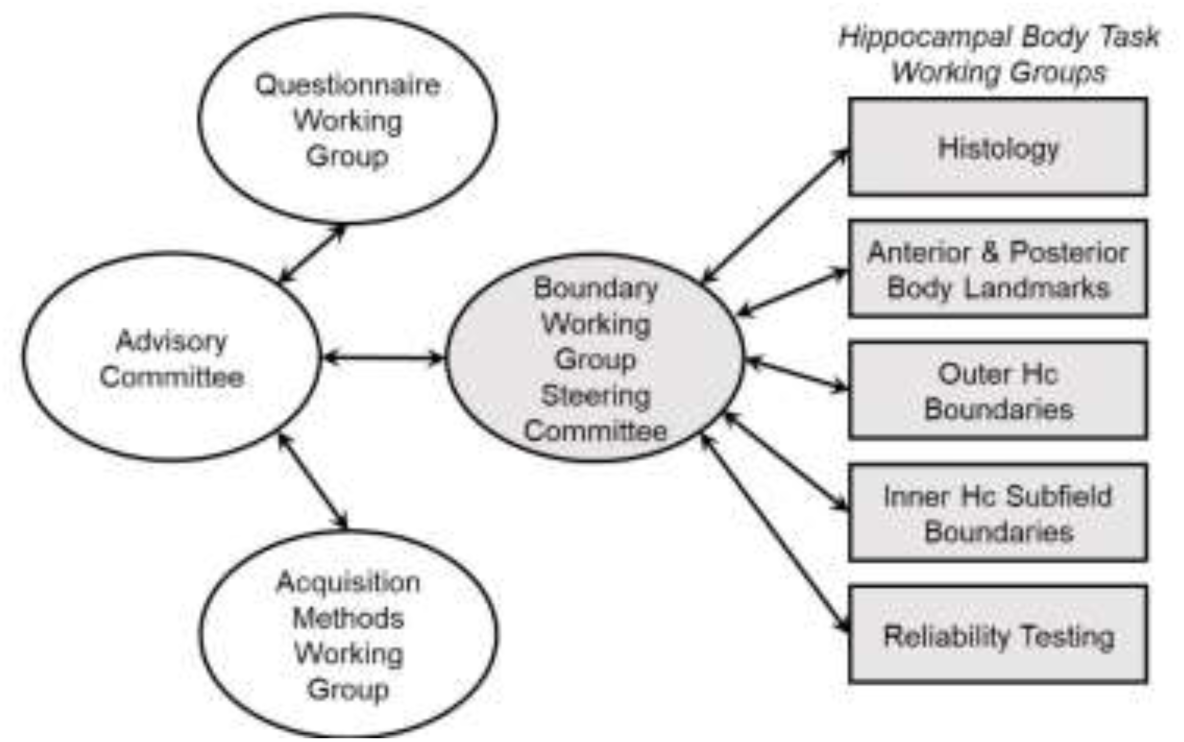
A flow chart of the Hippocampal Subfield Group (HSG) organization activity for supporting development and testing of the harmonized protocol for subfield segmentation in the hippocampal body. *Hc*—hippocampus.

Based on a survey of the literature and anatomical reference materials, the HSG Boundary Working Group began protocol development in the hippocampal body. This portion of the hippocampus begins immediately posterior to the uncus and terminates posteriorly with the last visualization of the lamina quadrigemina (Olsen et al., 2019). The hippocampal body is the largest portion of the structure along the anterior-posterior dimension (Daugherty et al., 2015; Malykhin et al., 2017; Poppenk et al., 2013), its anatomy is less complex than the anterior regions, and the subfield anatomy is relatively uniform over its span (Ding, 2013; Duvernoy et al., 2013; Insausti & Amaral, 2004). In this regard, the hippocampal body is well-suited to start developing a harmonized protocol. Because most of the existing protocols are restricted to the hippocampal body (Bender et al., 2018; Mueller & Weiner, 2009; Yushkevich et al., 2015), this first part of the harmonized protocol can be immediately adopted for many research questions while the head and tail protocols are still in development.

We developed and validated the hippocampal body protocol for a T_2_-weighted MRI sequence with a high in-plane coronal resolution (0.4 × 0.4 mm^2^), typically acquired with 2-mm slice thickness (e.g., Mueller et al., 2010; Yushkevich et al., 2010). In our previous review of the literature and technical requirements, we found T_2_-weighting optimal for visualization of key landmarks. For example, the internal structure of the hippocampus is partially defined by the stratum radiatum, lacunosum, and moleculare (SRLM), which is best visualized *in vivo* on T_2_-weighted images (Wisse et al., 2021). Because many of the structures to be delineated are smaller than a millimeter, high-resolution is needed for accurate segmentation (Canada et al., 2024; Wisse et al., 2017). A T_2_-weighted, 0.4 × 0.4 mm^2^ in-plane resolution sequence (typically anisotropic, with relatively thick slices) is one of the most commonly used sequences in applied research and clinical study of the hippocampal subfields as of this writing (see Homayouni et al., 2023; Iglesias et al., 2015; Wisse et al., 2017; Yushkevich et al., 2015).

1.1 *Overview of Protocol Development and Validation Process*

To develop a harmonized protocol for high-resolution T_2_-weighted images we used an “evidence-based Delphi panel” inspired by the EADC-ADNI working group that created a similar harmonized protocol for total hippocampal segmentation on common T_1_-weighted MRI (Boccardi et al., 2015). The Delphi procedure has several advantages for consensus building with a diverse representation of expertise in the field; the adaptation to introduce evidence for the evaluation and collect data from the evaluation process for iterative refinement accelerates protocol development and encourages wide adoption. In the initial process of surveying the existing methods (Wisse et al., 2017; Yushkevich et al., 2015), we noted a key difference between the scope of work for hippocampal subfield harmonization and the HarP development: namely, when we started our working group, there were no agreed upon, canonical definitions of hippocampal subfield nomenclature or boundaries on *in vivo* MRI.

Therefore, we designed a 5-step workflow to develop a harmonized, anatomically valid and reliable protocol for hippocampal subfields in the body (Figure 2; Olsen et al., 2019; Wisse et al., 2017). Due to the significant discrepancies among protocols in our initial survey, step 1 began by partnering with neuroanatomists to develop new histological reference materials to identify relevant landmarks and protocol definitions to then submit for Delphi procedure, as opposed to sequential voting on a set of existing protocols. In Steps 2 and 3, working groups were created with specific scopes of work for histology reference materials, identification of landmarks, and developing portions of the protocol. In Step 4, we held consensus voting and collected qualitative feedback from the broader HSG international network, and we would continue iteratively until consensus was met by statistical majority agreement. Following consensus, in Step 5 the protocol was tested and found to have strong inter- and intra-rater reliability, which has led to the finalization and now dissemination of the harmonized protocol for the hippocampal body.

**Figure 2.**
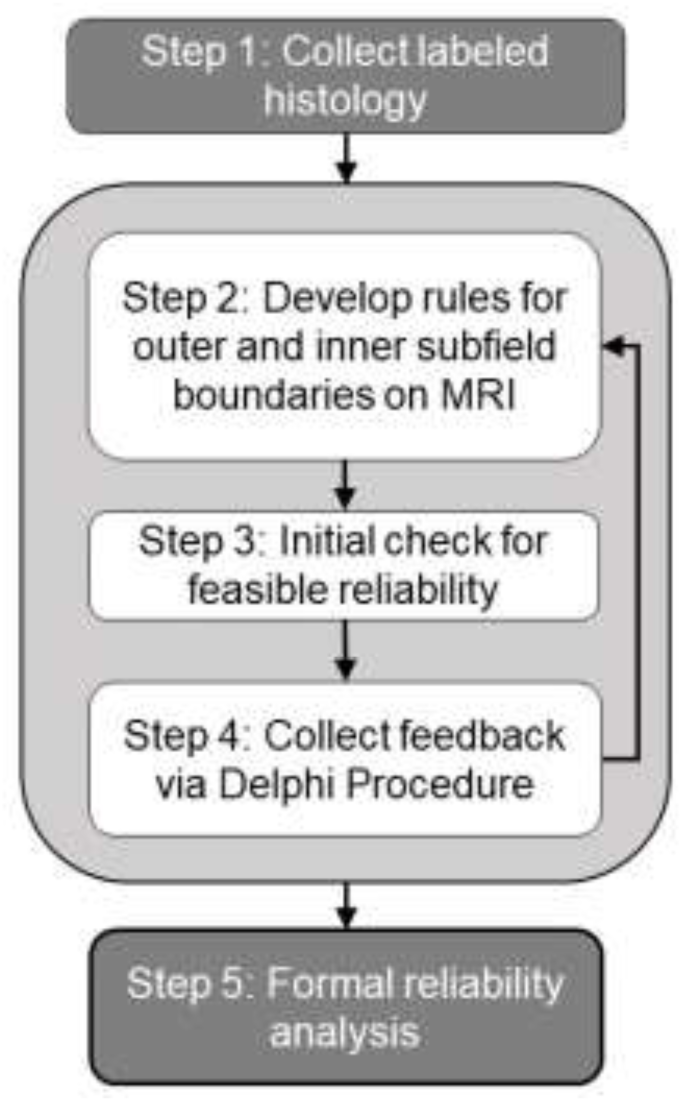
Depiction of the workflow for landmark identification, developing boundary definitions, and validation of the harmonized protocol.

## 2.0 Methods

### 2.1 Step 1: Collect labeled histology reference materials

Step 1 to collect labeled histology was implemented in a working group with international leaders in neuroanatomy and histology of the hippocampus: Drs. Ricardo Insausti (University of Castilla-La Mancha), Jean Augustinack (Massachusetts General Hospital), Katrin Amunts (Research Centre Jülich), and Olga Kedo (Research Centre Jülich). All new reference materials were designed to address limitations identified in existing neuroanatomy atlases and peer-reviewed publications of hippocampal subfield anatomy. Namely, inclusion of slices along the length of the hippocampal body to capture potential anterior-posterior variation; samples from multiple brains to represent individual differences; multiple tissue staining methods; and multiple neuroanatomists labeling the same slice images for direct comparison.

The histological dataset has been described in our previous progress report (Olsen et al., 2019). Briefly, high-resolution images of the stained sections were provided to the neuroanatomists for their annotation of hippocampal body subfields and the boundaries between adjacent subfields. Figure 3 shows representative images from the reference set with hippocampal subfield labels by different neuroanatomists. (See online supplement^i^ for additional reference images). Variability in boundary locations is noted across the images that is assumed to reflect true individual differences and anatomical variation throughout the structure, in addition to reliability of the neuroanatomists. Because these annotations represent expert judgments, all sources of variability were considered in the working group process when identifying landmarks and developing boundary definitions.

**Figure 3.**
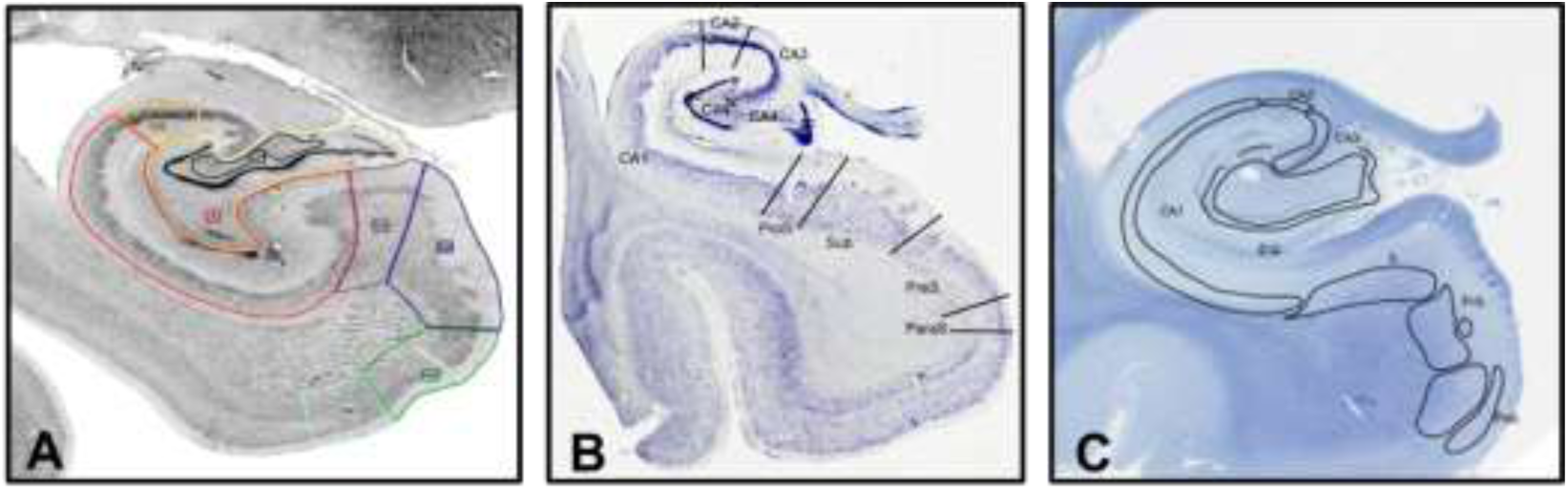
Example histology reference materials for the hippocampal body developed by the HSG to have representation of variability in hippocampal subfield definitions across brains and common histological procedures. Example images include (A) silver stain for cell bodies, (B) Nissl stain; and (C) the Klüver-Barrera method method. Regional labels include Cornu Ammonis (CA) sectors 1, 2, 3; some neuroanatomists include CA4 and the granular cell layer (gc); the dentate gyrus (DG); the subiculum (Sub); prosubiculum (ProS); presubiculum (PreS or PrS), and parasubiculum (ParS or PaS).

### 2.2 Step 2: Develop rules for outer and inner subfield boundaries on MRI

Working groups were structured for specific tasks in landmark identification and developing boundary definitions for the hippocampal body (described before in Wisse et al., 2017). The leads of each working group collaborated with the broader HSG community through multiple open meetings that were scheduled with major international scientific conferences 2014-2018 (hosted in Irvine, CA, USA; Chicago, IL, USA; San Diego, CA, USA; Montreal, Canada; London, England; Washington DC, USA; and Magdeburg, Germany).

We devised a working order to identify landmarks on *in vivo* MRI to denote the anterior and posterior limits of the hippocampal body, develop the outer boundary definitions to distinguish hippocampal tissue from surrounding white matter and cerebrospinal fluid (CSF), and demarcate he inner boundaries between adjacent hippocampal subfields in the coronal plane. The initial results for the anterior-posterior landmarks and the outer boundaries of the hippocampal body have been published (Olsen et al., 2019).

Here, we report the inner boundaries that are used to create labels for subiculum, CA subfields 1-3, and dentate gyrus. The subiculum label included portions of the pro-, pre- and para-subiculum regions that were labeled variably by the neuroanatomists. Each CA1, 2 and 3 subfields were defined separately. Although combining CA subfield labels (e.g., CA1-2) is common in the literature (Wisse et al., 2017; Yushkevich et al., 2015), we developed a protocol to label these regions separately to improve sensitivity and specificity to functional correlates, but the investigator may choose to combine labels based on their specific research question. An additional label for DG included the CA4 or hilus, separate from the CA3. We identified two possible rule definitions for the CA3-DG boundary—one based on a geometric heuristic that had strong evidence for feasibility, and another that referenced the endfolial pathway and may have relatively higher face validity in comparison to the histological reference materials. Both rules were tested for feasibility and presented for voting by Delphi procedure to determine which rule would be retained (see online supplement^i^ for complete alternative rule descriptions). The SRLM is the internal boundary that distinguishes the DG, which the working group determined to be included in the CA field region labels (i.e., excluded from DG) and the portion of the molecular layer that extends medially is included with the subiculum.

#### 2.2.1 Anatomical reference materials

The process of landmark identification and boundary definitions was completed in reference to the histology materials collected in Step 1; published neuroanatomy references (Amaral & Insausti, 1990; Ding, 2013; Ding & Van Hoesen, 2010; Duvernoy et al., 2013; Insausti & Amaral, 2004, 2012; Mai et al., 2015; Zeineh et al., 2001, 2015); and example *in vivo* neuroimaging data. In line with the goal of a harmonized protocol that can be applied in the field broadly, we made an open call to the HSG membership for MRI data sharing to collate example high-resolution (0.4 × 0.4 mm^2^ in plane), T_2_-weighted images collected in brains of children, younger adults and older adults; and with representation of common health co-morbidity (e.g., hypertension). We supplemented the shared data from HSG membership with additional scans representing cognitive impairment and Alzheimer’s disease-related pathology (Yushkevich et al., 2024) from the Alzheimer’s Disease Neuroimaging Initiative (ADNI) database (adni.loni.usc.edu). The ADNI was launched in 2003 as a public-private partnership, led by Principal Investigator Michael W. Weiner, MD. For up-to-date information on ADNI, see www.adni-info.org. From the collated scans, a common MR image reference set was developed for protocol development and feasibility studies to be used for the body protocol, in addition to future work for the head, tail and medial temporal lobe cortices.

### 2.3 Step 3: Initial check for feasible reliability

Early in the working group process, we identified that information about feasibility to meet standards for reliability was important information for experts to reference during the Delphi procedure for consensus voting. The initial feasibility check was on a small, representative image set to provide reliability estimates and rater qualitative feedback for the subsequent Delphi procedure. Two expert raters (more than 4 years of experience manually segmenting the hippocampus and subfields on MRI), who were naïve to the protocol prior to training, participated in the feasibility assessment. We have previously reported the anterior-posterior ranging protocol as highly reliable (Olsen et al., 2019). In this stage of the protocol development, the anterior-posterior ranges were provided to the raters. Training included detailed documentation with example image tracings, a 2-hour introductory training session (via video conference), followed by prescribed practice and then an additional 1-3 hours of individualized feedback (via video conference).

The feasibility study dataset included brains of healthy, typically developed children (n = 2, both male, age 9 and 15 years), healthy adults (n = 2, female age 31, and male age 66), and dementia of the Alzheimer’s type collected by ADNI (n = 1, female age 70). Between-rater intraclass correlation coefficients (*ICC* (2,k); (Shrout & Fleiss, 1979)) and average Dice Similarity Coefficient (*DSC*; (Dice, 1945; Zou et al., 2004)) were calculated for bilateral labels. Raters indicated how well they understood the protocol and their confidence when applying the protocol on a 7-level Likert scale: e.g., asked if they understood the protocol, responses were recorded 1 = Not at all to 7 = Extremely well. Open responses were recorded to provide additional qualitative information. For ease of use and standardization during the protocol development, all segmentations were made with the freely available ITKSnap software (Yushkevich et al., 2006; http://www.itksnap.org/pmwiki/pmwiki.php; last accessed 04/27/2025). However, it should be noted that the protocol can be implemented in any modern available software that allows manual segmentation.

### 2.4 Step 4: Collect feedback via Delphi voting procedure

We applied an “evidence-based Delphi panel” procedure, similar to that developed by Boccardi and colleagues (Boccardi et al., 2015) when creating the harmonized protocol for (total) hippocampal segmentation on T1-weighted MRI. Briefly, this procedure presented quantitative and qualitative evidence to a panel of experts to apply when voting on agreement of landmark and boundary definitions to be used for segmentation. The Delphi procedure was anonymous and recursive until experts reached consensus for all components of the segmentation protocol. Delphi panel participants rated each rule on a 9-level Likert scale for agreement (1 = do not agree at all; 9 = fully agree) and clarity (1 = extremely unclear and requires major revisions; 9 = extremely clear and requires no changes), with the option to indicate no opinion that would not count in consensus evaluation. Open fields collected additional qualitative feedback. Consensus was declared when the number of agreement responses (Likert rating 6-9) was statistically greater than the number of disagreements. This definition of consensus is more conservative than the definition in a traditional Delphi method, which uses median greater than 5 in the 9-level Likert rating for declaring consensus (Boccardi et al., 2015). If consensus was not achieved, the quantitative ratings and qualitative comments collected during the Delphi procedure were used to revise the boundary definitions; and the procedure repeated iteratively until consensus was reached for all boundary rules. If statistically significant consensus on a given rule was not reached after four rounds, the details of the rule agreed upon by the majority of respondents would be taken as the final rule. Responses were collected via Qualtrics, version December 2017 (https://www.qualtrics.com; Qualtrics, Provo, UT). We recruited participants with an open call via HSG member email listserv (approximately 200 subscribers), website and social media accounts; and participants were encouraged to share the call with other investigators who had relevant expertise. Included responses for analyses were confirmed to have self-reported expertise relevant to human hippocampal anatomy and its segmentation on *in vivo* MRI. To limit non-independence of responses, we instructed lab groups to complete one questionnaire together, which was confirmed based on the reported principal investigator before anonymizing responses for analysis. The Delphi procedure for the anterior-posterior landmarks and outer boundaries of the hippocampal body was completed December 2017 – March 2018 and reported previously (Olsen et al., 2019); here we report the procedure for the inner subfield boundaries (survey dates December 2021 – April 2022).

#### 2.4.1 Information provided to the panel for the Delphi procedure

The Qualtrics questionnaire presented the complete protocol description and example image with segmentations, and then each landmark or boundary definition was summarized for evaluation with contextual information, relevant evidence with example images, and acknowledged limitations. A summary of this information was included in the Qualtrics questionnaire, with additional reference materials and detailed explanation in a 79-page supplement (see online supplement^i^). Protocol training documents, a video recording of the protocol as an applied segmentation, example MRI with segmentations by the two expert raters from the feasibility assessment, and the resulting data were also available for download. Delphi panelists were encouraged to download the supplemental materials and try a first-hand experience with the protocol before reporting on their level of agreement. The survey also assessed the community’s preference between two alternative definitions for the boundary between CA3 and DG: one based on a geometric heuristic and the other by approximating the endfolial pathway (see online supplement^i^ for additional details).

### 2.5 Step 5: Formal reliability analysis on consensus protocol

Following the consensus by Delphi procedure on the inner boundaries, the rule definitions were combined with the consensus definitions for outer hippocampal boundaries (Olsen et al., 2019) to create one protocol for formal reliability analysis before accepting as final. Three raters who were naïve to the protocol were assembled: 2 expert raters (with more than 4 years of experience manually tracing hippocampal subfields on T_2_-weighted MRI) and a rater with no manual segmentation experience. Training materials, video demonstration, practice MRI scans and example tracings were provided to the raters. All raters met with the trainer by video conference to review the documentation, general procedures in ITK-Snap, and a brief demonstration with time for questions. All raters completed practice segmentations on 2-5 scans and received specific feedback from the trainer in subsequent online meetings. All raters required the trainer’s approval before starting reliability testing. This procedure was similar to training in a lab setting and limited to 2-3 contact hours to mimic the anticipated scalability of the procedures for dissemination of the protocol to the broader research community.

An additional blinded set of MRI scans was reserved for reliability testing with blinded review. The tracers were provided the anterior-posterior body ranges and had no other information on scan demographics or study of origin. The reliability dataset included N = 24 brains representing healthy, typically developing children (n = 7, ages 4-15), healthy adults and those with hypertension without dementia (n = 9, ages 31-94), adults with mild cognitive impairment (n = 5, ages 70-75) and with dementia (n = 3, ages 76-79). MRI scans were sourced from data sharing by multiple members of the HSG; and all scans with cognitive impairment and five of the scans in cognitive-typical older adults were sourced from ADNI. The dataset was representative of the practices in the broad research field: all images were collected with 0.4 × 0.4 mm^2^ resolution in the coronal plane and 2-mm slice thickness, aligned perpendicular to the hippocampal long axis; and were at 3 tesla field strength (manufacturers differed by site, including Siemens and Phillips). Images were selected following common quality control (Canada et al., 2024) for visualization of the SRLM as a key landmark for the protocol, and included mild-to-moderate forms of common imaging artifacts related to motion or reconstruction error. Inter-rater reliability was assessed among the 3 raters, in addition to intra-rater reliability for an expert rater and the novice rater following >2-week delay.

### 2.6 Statistical Analyses

During the Delphi procedure, ratings were reported on 9-level Likert scales, and responses were summarized with descriptive statistics. Consensus was determined by statistical majority of agreement, recoding agreement response (Likert rating 6-9) vs. no agreement (Likert rating 1-5), and clear rule description (Likert rating 6-9) vs. unclear (Likert rating 1-5), with differences in frequency assessed by binomial tests (α = 0.05). In establishing reliability of the harmonized protocol, the reliability metrics included inter-rater reliability (*N* = 24) of volumes by *ICC*, assuming random raters (*ICC*(2,k);(Shrout & Fleiss, 1979)) and average *DSC* among all pairs of raters (Dice, 1945; Zou et al., 2004). Consistency of inter-rater *DSC* reliability was tested with non-parametric Mann-Whitney U tests of the distributions between child and adult age groups, and by cognitive diagnosis with the data sourced from ADNI. Intra-rater reliability was tested for absolute agreement (*ICC*2,1) and average *DSC* of the rater with themselves on a subset of scans (n = 11) for one expert and the novice rater.

## 3.0 Results

### 3.1 Step 1: Collected Histology Reference Materials

The expert neuroanatomists provided independent labels of the histological dataset representing the anterior-posterior length of the hippocampal body in the coronal plane, similar to common neuroimaging procedures. The neuroanatomists referenced similar cytoarchitectural details when making labels, and yet differences emerged in the expert judgment on the boundary location between adjacent subfields. (See online supplement^i^ for complete set of labeled images.) As a prime example of the value of this additional histological reference dataset, we will briefly describe the information gained on the CA1-Sub boundary. Similar to the pattern of differences we originally observed between different neuroimaging protocols (Yushkevich et al., 2015), the disagreement between neuroanatomists on the location of the CA1-Sub boundary was the most prominent (Figure 4). Some neuroanatomists labeled pro-subiculum, as a transition region from Sub to CA1 (see Ding & Van Hoesen, 2015; Rosenblum et al., 2024). Taking this into account, for all neuroanatomists the location of the Sub-CA1 boundary fell within a medial-lateral range of possible positions on a slice (see Fig 4) that was consistent with anatomical variation noted by other histological studies (Ding, 2013; Zeineh et al., 2015), and the range of boundaries in existing MRI segmentation protocols (Yushkevich et al., 2015). When compared between slices in the anterior and posterior hippocampal body, there was relative consistency for an individual brain, although between-individual differences were noted (see online supplement^i^ for details). The different sources of variability (i.e., different expert judgment, individual differences and anterior-posterior differences) were all considered when developing the protocol definitions that applied a geometric heuristic to align the internal subfield boundaries in reference to hippocampal macrostructural landmarks.

**Figure 4.**
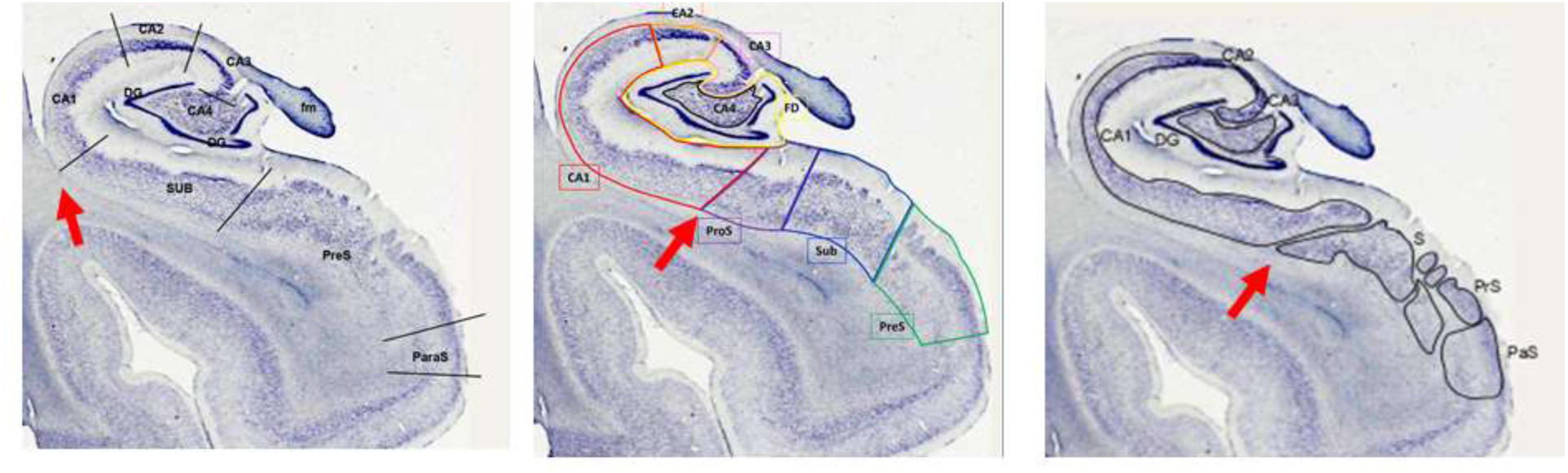
Example anatomical labels between hippocampal subfields on the same slice of a Nissl-stained brain by 3 expert neuroanatomists. The variability in annotation style reflects the differences among neuroanatomists. The red arrow indicates the location of the CA1-Sub boundary. Regional labels include Cornu Ammonis (CA) sectors 1, 2, 3; some neuroanatomists include CA4 and the granular cell layer (gc); the dentate gyrus (DG); the subiculum (Sub); prosubiculum (ProS); presubiculum (PreS or PrS), and parasubiculum (ParS or PaS).

### 3.2 Step 3: Initial check for feasible reliability of proposed protocol to combine inner and outer boundaries

The initial feasibility assessment suggested that the protocol could be implemented reliably, pending formal testing: all *ICC*(2) ≥ 0.83, except Sub (0.45), and all *DSC* ≥ 0.61. The lower reliability of the Sub volume was due to ambiguity in the procedure for the medial boundary with cortex, which was subsequently revised with feedback on rule clarity collected during the Delphi procedure. Additional assessment of the rater experience and understanding of the protocol are available in the online supplemental materials^i^. The feasibility test results and qualitative reporting were provided as background information for the “evidence-based Delphi panel”.

### 3.3 Step 4: Delphi procedure evaluating the inner boundaries between subfields in the hippocampal body

The Delphi procedure included responses from 26 participating laboratories, all indicating at least 4 years previous experience with manual segmentation of the hippocampal subfields, and 85% of labs had more than 5 years’ experience. All labs had experience with hippocampal measures on 3 tesla data and 92% of labs had experience with relevant T_2_-weighted images. With one iteration, all inner boundary definitions were found to be clear and achieved consensus (all binomial test *p-*values ≤ 0.004; Table 1). No revisions to rule definitions were required for consensus, although the working group used the feedback on clarity to implement minor changes to wording and sample images.

**Table 1.** Summary of Delphi evaluation responses for inner boundary definitions based on geometric heuristic.

| Rule Definition | Question | *N* | Range | *M* | *SD* | % Agree | binomial test |
| --- | --- | --- | --- | --- | --- | --- | --- |
| Sub-CA1 Boundary | How clear is the boundary description? | 26 | 7 - 9 | 8.42 | 0.64 | 100 | < 0.001 |
|  | Do you agree with the boundary rule? | 25 | 3 - 9 | 8.08 | 1.41 | 96.0 | < 0.001 |
| CA1-CA2 Boundary | How clear is the boundary description? | 26 | 5 - 9 | 8.46 | 0.90 | 96.2 | < 0.001 |
|  | Do you agree with the boundary rule? | 23 | 1 - 9 | 8.09 | 2.13 | 87.0 | < 0.001 |
| CA2-CA3 Boundary | How clear is the boundary description? | 26 | 7 - 9 | 8.58 | 0.64 | 100 | < 0.001 |
|  | Do you agree with the boundary rule? | 23 | 1 - 9 | 8.09 | 2.02 | 91.3 | < 0.001 |
| CA3-DG Boundary | How clear is the boundary description? | 26 | 7 - 9 | 8.65 | 0.56 | 100 | < 0.001 |
|  | Do you agree with the boundary rule? | 22 | 1 - 9 | 7.14 | 2.27 | 81.8 | 0.004 |
| Treatment of SRLM | How clear is the boundary description? | 26 | 5 - 9 | 8.50 | 0.95 | 96.2 | < 0.001 |
|  | Do you agree with the boundary rule? | 24 | 3 - 9 | 8.21 | 1.47 | 91.7 | < 0.001 |

*Note:* Boundary definitions based on the geometric heuristic were presented with relevant evidence and contextual information alongside the questions for evaluation. The SRLM was defined as an internal structure to be included within the label for subiculum and CA subfields (i.e., excluded from DG). Ratings were made on a 9-level Likert scale; responses > 5 were considered endorsement of clarity or agreement. Consensus was determined by statistical majority agreement with significant binomial test ( $\alpha = 0.05$ ). *N* = 26 laboratories participated; items with fewer responses are due to indicating no opinion and were omitted from summary calculations for consensus evaluation. *Sub*—subiculum; *CA*—*Cornu Ammonis*; *DG*—*dentate gyrus*; *SRLM*—stratum radiatum, lacunosum, and moleculare.

When presented with the alternative rule definition for the CA3-DG boundary that referenced the endfolial pathway (see online supplement^i^ for details), the majority preferred the geometric heuristic (57.69%), fewer preferred the endfolial pathway rule (23.08%), and a similar percentage had no preference (19.23%). A representative comment relating to the endfolial pathway rule underscores the need to balance reliability with face validity: “Overly complex [and] is prone to errors and more importantly cannot be properly assessed for validity with MRI compared to neuroanatomical sections. Therefore, the additional potential validity is mitigated by complex instructions (prone to errors).” Therefore, the geometric heuristic rule set to define all internal boundaries between the hippocampal subfields, including the CA3-DG boundary, was retained for the final protocol development.

Although all boundary definitions in the retained final protocol had statistical majority agreement (81-96% responses), there was a range of responses that included individuals who disagreed (see Table 1). The qualitative comments provided some insights into the ratings and informed the discussed limitations of the protocol (see Table 2 for representative comments; the online supplement^i^ reports all comments).

**Table 2.** Representative qualitative comments collected during the Delphi procedure on rules for defining subfield boundaries in the hippocampal body.

| Theme of Constructive Feedback | Example Comment | Agreement Rating |
| --- | --- | --- |
| Delineation of CA2 may be misaligned | "I am concerned that the current segmentation of CA2 doesn't fully capture the CA2 subregion...Given the fact that the CA2 subregion is so small, I would be concerned about potential interpretations of CA2 segmentations, and especially if individuals are using CA2 segmentations to look at functional activation. I'm not convinced at the moment that you can delineate CA2." | 1 |
|  | "I would suggest including CA2 as part of CA3, given how hard it is to distinguish these two. I'm also concerned that as we move anterior to posterior, the boundaries between CA2 and CA3 may shift, and I don't know if the current rule would capture that shift well enough." | 1 |
|  | "The segmentation will likely not be perfect (proposed rule more lateral than expert ratings), but we agree that the proposed boundary definition is okay. As a general remark, one may need to be careful when investigating CA2 alone and be aware that one might not only be measuring CA2." | 9 |
| Specificity of defining CA3 apart from DG | "The inferior part of the CA3 may extend too far into DG, but this may also be due to the rather flat HC in the example... So, we are generally in agreement with this rule given the trade-off between accuracy and complexity." | 7 |
|  | "From the current geometric rule, I'm concerned since DG/CA3 are very wrapped around each other and with the extent of DG/CA3 boundaries shifts from anterior to posterior." | 1 |
|  | "We have concerns with this boundary since it is sampling DG within CA3, although it seems easy to follow the instructions and relatively consistent between raters." | 3 |
| Treatment of SRLM | "I agree with the boundary, however, similar to one of the raters, I would also like to note the potential for biases towards greater volume in CA regions and subiculum when the SLRM is more difficult to distinguish." | 5 |
|  | "We believe this structure should be given a distinct label. Some consider this structure to be white matter given the lack of neuronal cell bodies, and it may also include blood vessels and CSF pockets along the hippocampal sulcus which should ideally be excluded (or at least separated) from subfield volume measures." | 3 |

*Note:* Representative comments providing constructive critique of rules in the harmonized protocol are reported here arranged by
theme and the relevant individual rating on agreement for the relevant rule (1 = Strongly disagree; 9 = Strongly agree).

One common theme in the qualitative comments from Delphi respondents was on the delineation of CA2, which shares boundaries with CA1 and CA3. The CA2 region is the smallest subfield to segment and thus particularly vulnerable to segmentation error and misalignment. The region has distinct cytoarchitecture that may be critical for memory (Ding et al., 2010), and so it is desirable to create a separate label for the region despite the challenge of its small size. The feedback from the Delphi panel is included as a limitation of the protocol applied to images with 0.4 x 0.4 mm^2^ in-plane resolution and informs a discussion of alternative approaches to combine CA2 with adjacent subfield labels (e.g., CA1-2).

A second theme in the comments was in the specificity of the definition of CA3 apart from DG. These regions are closely connected and present with complex morphometry that has the CA3 folding into the DG following hippocampal development as an allocortical structure (Duvernoy et al., 2013; Insausti & Amaral, 2004; Zeineh et al., 2001). The comments from the Delphi panelists align with the challenges the working group experienced to have a definition that had strong face validity with the available reference materials and would have good reliability. As seen in the example comments from the Delphi procedure, the balance between reliability and face validity was weighed when indicating for agreement of the boundary definition. In addition, there was reference to not having a distinct label for the CA4/hilus region. Although neuroanatomists generally agree on the presence of this region, there is disagreement among neuroanatomists on naming and allocation as CA4 region (Ding, 2013; Duvernoy et al., 2013; Williams et al., 2023) or as the hilus of the dentate gyrus (Insausti & Amaral, 2004). Based on the resolution and contrast of the typical high-resolution T_2_-weighted MRI on 3 tesla, the working group did not separately label this region and it is included in the DG label. These details are included to qualify the protocol use and external validation based on the histological reference materials available to date.

A third point of discussion in the Delphi panel was on the treatment of the SRLM. The SRLM lines the internal edge of the hippocampal fissure and separates the DG from the CA subfields. The protocol includes the SRLM within the CA1-3 subfield labels (excluded from the DG), and the molecular layer extension medially is included in the subiculum label. This protocol decision was informed by review of published manual segmentation of ultra high-resolution (∼200 μm^3^ isotropic) *ex vivo* MRI with histology that identified the dark band on T_2_-weighted MRI falls within the CA-SRLM region (Adler et al., 2018; de Flores et al., 2020). Due to limitations of typical high-resolution *in vivo* imaging (0.4 × 0.4 mm^2^ in plane), the working group determined it was not feasible to have a separate, reliable label for SRLM, and therefore it was included in the CA field labels, and the extension of the molecular layer within the Sub label. This is a limitation of the protocol that was also noted by the Delphi panelists.

### 3.4 Step 5: Formal reliability analysis of the consensus protocol

Following consensus, inter-rater reliability was tested on the sample of *N* = 24 brains and found to be excellent for all regions, with lower but acceptable reliability of the CA2 label (Table 3). Reliability between the two expert raters (all *ICC*(2,2) = 0.63-0.93; all *DSC* = 0.59-0.85) was similar to the novice with either expert (all *ICC*(2,2) = 0.57-0.97, except CA2-R for one comparison was 0.45; all *DSC* = 0.60-0.87). Intra-rater reliability by one expert and one novice rater following > 2-week delay (*n* = 11) indicated excellent agreement (Table 3), and was consistent between the expert rater (all *ICC*(2,1) = 0.89-0.97; all *DSC* = 0.68-0.89) and the novice rater (all *ICC*(2,1) = 0.75-0.97; all *DSC* = 0.69-0.91). Similar reliability ratings between expert and novice raters suggest the protocol can be consistently applied regardless of prior experience with manual segmentation or knowledge of hippocampal anatomy, although experience with the specific protocol is expected to confer some stability of the skill.

**Table 3.** Summary inter- and intra-rater reliability of the harmonized protocol.

| Region | Inter-Rater |  | Intra-Rater |  |
| --- | --- | --- | --- | --- |
|  | (3 raters, n = 24) |  | (2 raters, n = 11) |  |
| | $ICC(2,k)$ | Avg. $DSC$ | Avg. $ICC(2)$ | Avg. $DSC$ |
| Sub-L | 0.87 | $0.86 \pm 0.01$ | 0.87 | $0.90 \pm 0.02$ |
| Sub-R | 0.87 | $0.84 \pm 0.01$ | 0.93 | $0.89 \pm 0.02$ |
| CA1-L | 0.88 | $0.84 \pm 0.01$ | 0.84 | $0.88 \pm 0.03$ |
| CA1-R | 0.94 | $0.83 \pm 0.02$ | 0.88 | $0.88 \pm 0.03$ |
| CA2-L | 0.66 | $0.61 \pm 0.02$ | 0.89 | $0.73 \pm 0.08$ |
| CA2-R | 0.76 | $0.62 \pm 0.03$ | 0.90 | $0.69 \pm 0.11$ |
| CA3-L | 0.91 | $0.70 \pm 0.02$ | 0.96 | $0.80 \pm 0.05$ |
| CA3-R | 0.90 | $0.71 \pm 0.03$ | 0.96 | $0.76 \pm 0.08$ |
| DG-L | 0.86 | $0.82 \pm 0.03$ | 0.86 | $0.87 \pm 0.04$ |
| DG-R | 0.91 | $0.80 \pm 0.03$ | 0.93 | $0.86 \pm 0.05$ |

*Note:* Reliability was tested on a blinded dataset by two expert raters and 1 novice rater ( $k = 3$ raters), all naïve to the protocol. Average *ICC*, and average *DSC* ( $\pm SD$ ) are reported. *ICC*—*intra-class correlation coefficient*; *DSC*—*Dice Similarity Coefficient*; *Sub*—*subiculum*; *CA*—*Cornu Ammonis*; *DG*—*dentate gyrus*.

Average reliability was also similar among raters regardless of scanned individual’s demographic characteristics. With the available sample, we provide initial evidence that there is no differential reliability systematic with age (children vs. adults), or by cognitive impairment among older adults following ADNI procedures (Table 4).

**Table 4.** Summary of average inter-rater dice similarity coefficient by scan demographic.

|  | Age-Group Comparisons |  |  | Comparisons by Cognitive Diagnosis |  |  |  |
| --- | --- | --- | --- | --- | --- | --- | --- |
|  | Children | Adults | *p*- | Cognitive | MCI | Dementia | *p*- |
| Region | (*n* = 7) | (*n* = 17) | value | Typical (*n* = 5) | (*n* = 5) | (*n* = 3) | value |
| Sub-L | $0.86 \pm 0.03$ | $0.86 \pm 0.02$ | 0.80 | $0.87 \pm 0.01$ | $0.87 \pm 0.01$ | $0.85 \pm 0.03$ | 0.59 |
| Sub-R | $0.85 \pm 0.02$ | $0.84 \pm 0.02$ | 0.32 | $0.85 \pm 0.02$ | $0.85 \pm 0.03$ | $0.82 \pm 0.02$ | 0.29 |
| CA1-L | $0.86 \pm 0.02$ | $0.83 \pm 0.03$ | 0.06 | $0.84 \pm 0.02$ | $0.81 \pm 0.03$ | $0.82 \pm 0.04$ | 0.21 |
| CA1-R | $0.85 \pm 0.01$ | $0.82 \pm 0.03$ | 0.17 | $0.83 \pm 0.02$ | $0.82 \pm 0.02$ | $0.81 \pm 0.06$ | 0.66 |
| CA2-L | $0.62 \pm 0.08$ | $0.60 \pm 0.10$ | 0.71 | $0.63 \pm 0.09$ | $0.55 \pm 0.14$ | $0.61 \pm 0.02$ | 0.84 |
| CA2-R | $0.63 \pm 0.10$ | $0.62 \pm 0.09$ | 0.76 | $0.64 \pm 0.07$ | $0.60 \pm 0.06$ | $0.55 \pm 0.10$ | 0.24 |
| CA3-L | $0.73 \pm 0.06$ | $0.70 \pm 0.06$ | 0.26 | $0.73 \pm 0.03$ | $0.65 \pm 0.09$ | $0.69 \pm 0.02$ | 0.11 |
| CA3-R | $0.70 \pm 0.09$ | $0.72 \pm 0.04$ | 0.46 | $0.73 \pm 0.05$ | $0.72 \pm 0.05$ | $0.70 \pm 0.03$ | 0.60 |
| DG-L | $0.83 \pm 0.05$ | $0.81 \pm 0.04$ | 0.35 | $0.84 \pm 0.03$ | $0.81 \pm 0.05$ | $0.79 \pm 0.04$ | 0.21 |
| DG-R | $0.82 \pm 0.04$ | $0.80 \pm 0.04$ | 0.32 | $0.81 \pm 0.03$ | $0.81 \pm 0.04$ | $0.77 \pm 0.04$ | 0.24 |

*Note:* Average dice similarity coefficient ( $DSC$ ) $\pm SD$ is reported stratified by age group and cognitive diagnosis that was associated with the scan. Distributions were compared for statistically significant difference by Mann-Whitney U test ( $\alpha = 0.05$ ). Comparisons by cognitive diagnosis were made only among scans collected by ADNI and included standardized protocol for diagnosis. *MCI—Mild cognitive impairment; Sub—subiculum; CA—Cornu Ammonis; DG—dentate gyrus.*

### 3.5 HSG Harmonized Protocol for segmenting subfields in the hippocampal body

Following confirmation of reliability, we have formalized the procedures as the HSG Harmonized Protocol for subfield segmentation in the hippocampal body. The hippocampal subfields are drawn to be contiguous (sharing internal boundaries) and label the entire hippocampal body volume (Figure 5). The procedure begins by selecting the anterior-posterior range of the hippocampal body (see Olsen et al., 2019), and then applying a geometric heuristic in reference to macrostructural landmarks of the hippocampus and surrounding neuroanatomy to approximate the inner boundaries between contiguous subfield regions (summarized in Figure 6). The hippocampal subfield regions are then segmented by applying the outer boundaries of the hippocampus (previously described in Olsen et al., 2019) with the inner boundaries, as detailed here. A brief summary of the complete, harmonized segmentation protocol for the subfields in the hippocampal body is provided here; detailed procedures, examples and training materials are available for download (https://hippocampalsubfields.com/harmonized-protocol/).

**Figure 5.**
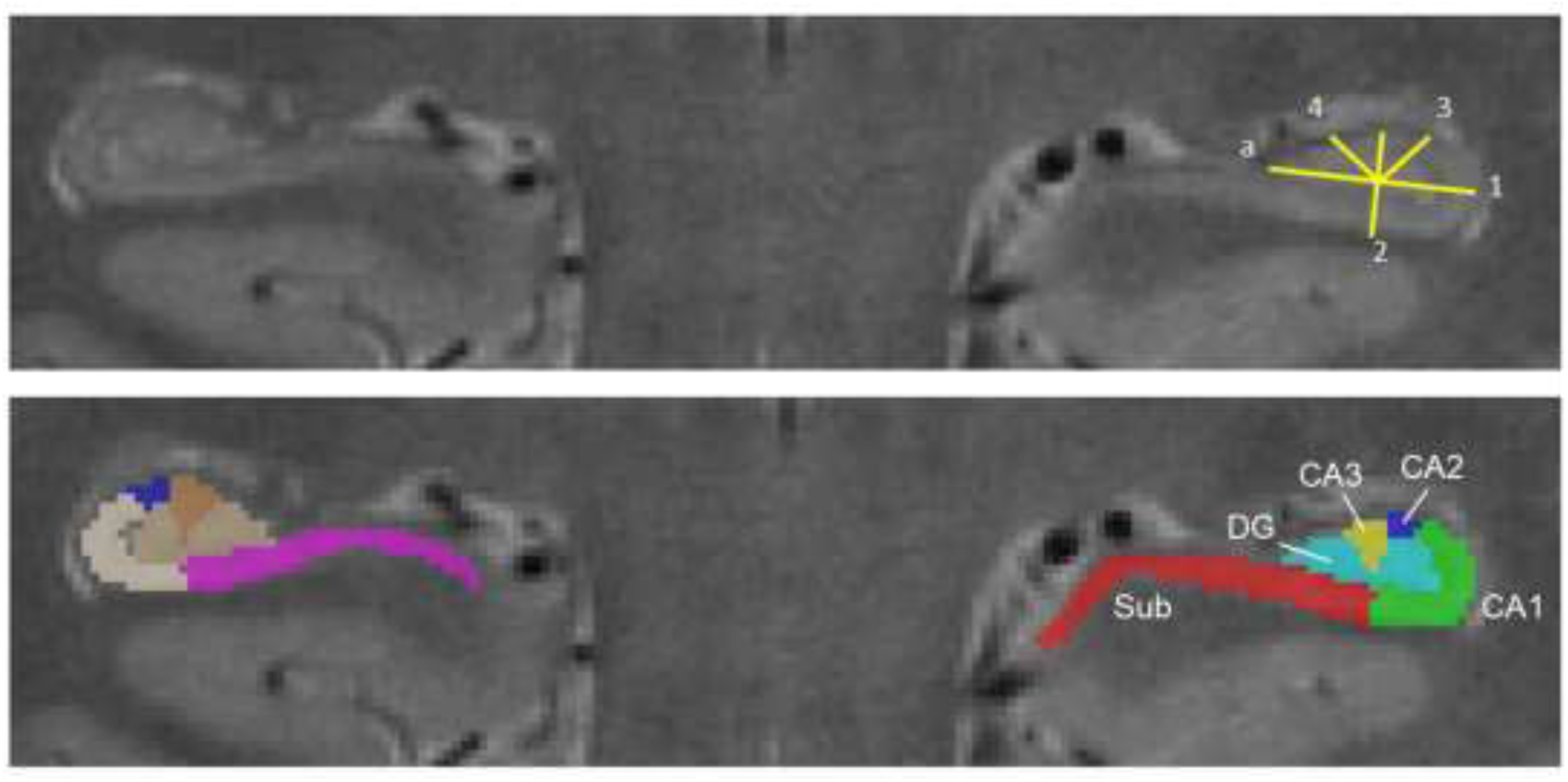
Hippocampal Subfield Group (HSG) Harmonized Protocol for segmenting subfields in the hippocampal body. The protocol is illustrated on a high-resolution (0.42 x 0.42 mm^2^ in-plane), T_2_-weighted MRI showing the same image with the geometric heuristic illustrated (top) and the subfield segmentation labels (bottom; Sub—subiculum; CA—Cornu ammonis; DG— dentate gyrus).

**Figure 6.**
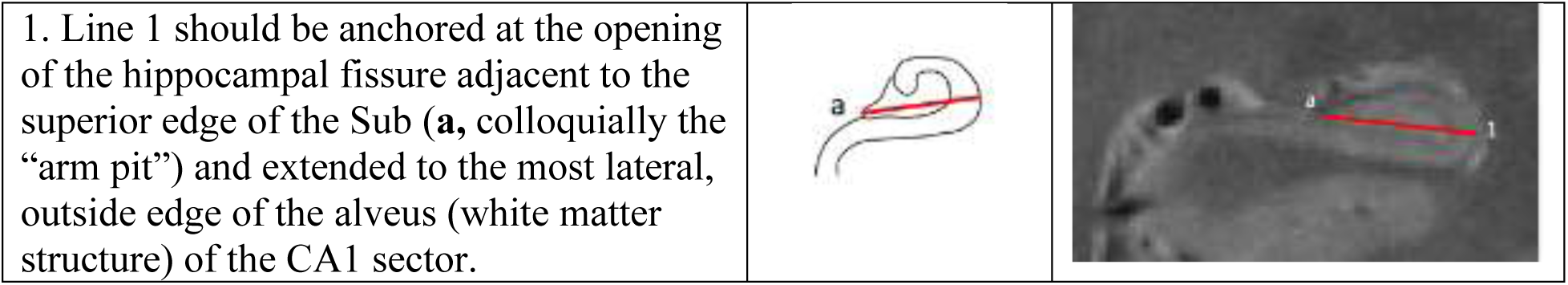
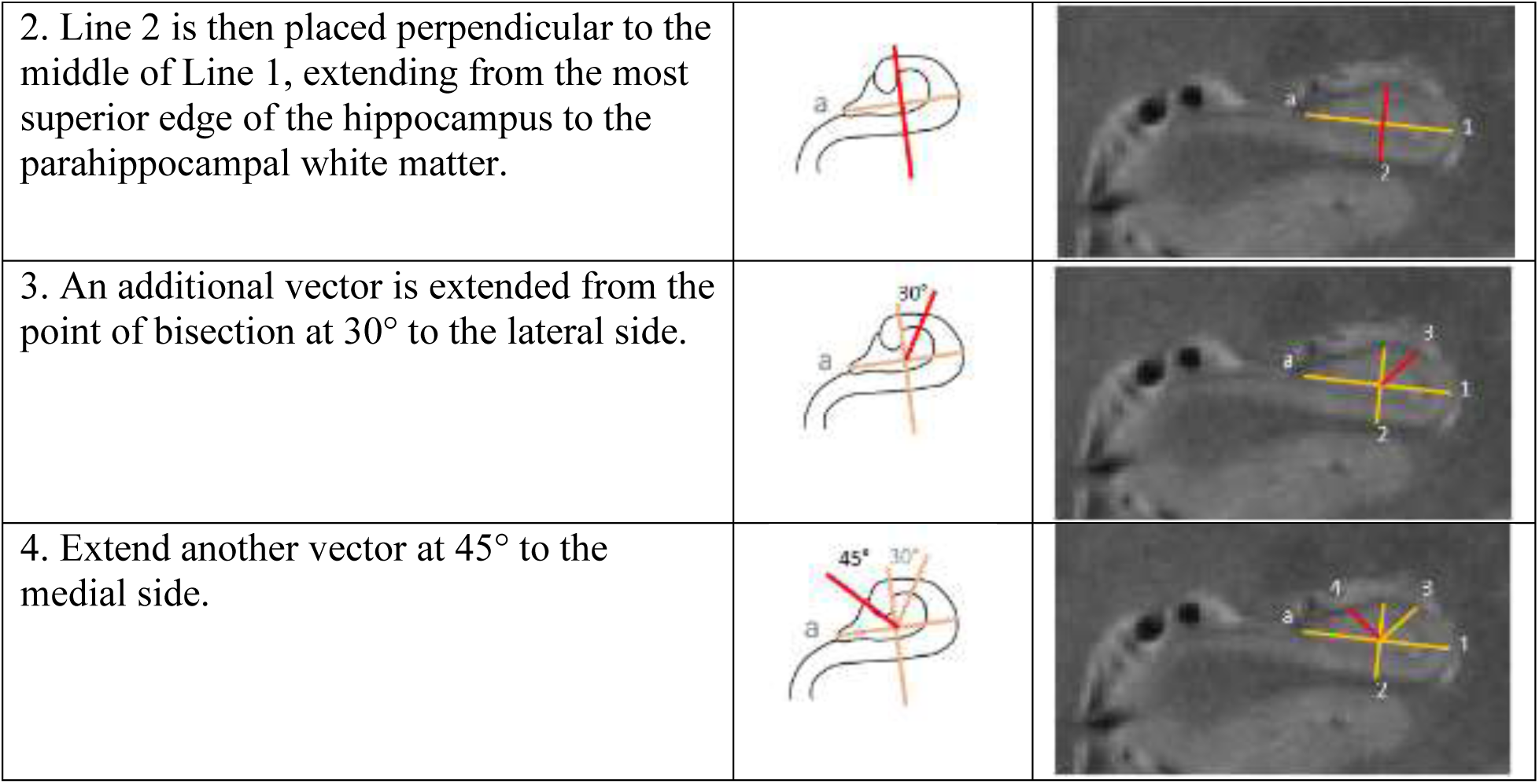
Summary of the geometric heuristic applied to the hippocampus on high-resolution (0.4 x 0.4 mm^2^ in-plane) T_2_-weighted images to approximate the location of boundaries between contiguous hippocampal subfields. This refers to the inner boundaries of the hippocampal subfields, and when combined with the outer boundary definitions, allows segmentations of subfield areas throughout the length of the hippocampal body.

#### 3.5.1 Placement of the Geometric Heuristic Definitions to Identify Inner Boundaries between Hippocampal Subfields

See Figure 6 for a summary illustration of the steps in the geometric heuristic line placement.

#### 3.5.2 Complete harmonized segmentation protocol to apply outer and inner boundaries to label subfields in the hippocampal body

##### 3.5.2.1 Subiculum label

The Sub label includes portions of the pro-, pre- and para-subiculum regions that the neuroanatomists labeled with subiculum. In the anterior body, the medial Sub-cortex boundary is defined as a horizontal line extending from the most medial superior aspect of the parahippocampal white matter to the CSF. In the posterior body, the Sub-cortex boundary is defined as the superior medial point—it extends to the medial edge of the hippocampus where it meets the parahippocampal gyrus; visualized as the medial portion of the Sub tapering and terminating at the beginning of the calcarine sulcus, appearing as a “notch”. Across the anterior-posterior length, the lateral boundary is the CA1-Sub boundary defined at the inferior portion of Line 2, spanning from the SRLM/molecular layer to the white matter. The superior Sub boundary is drawn to include the SRLM/molecular layer. The inferior boundary is drawn on the border of the adjacent white matter.

##### 3.5.2.2 CA1 label

The medial/internal boundary is at the CA1-Sub border (the inferior portion of Line 2, spanning from the SRLM to the white matter). The CA1 boundary is drawn to include the SRLM, and to exclude it from the dentate gyrus. Any visualization of a hippocampal sulcal cyst should be excluded. A hippocampal sulcal cyst (or cavity) is a CSF-filled space in the hippocampal fissure (most commonly between the SRLM and the inferior, lateral edge of the DG). The external white matter is excluded from the lateral boundary.

##### 3.5.2.3 CA2 label

The medial boundary of CA2 is the superior portion of Line 2, marking the location of the CA2-3 boundary. The lateral boundary is the 30° vector to the superior lateral edge of the hippocampus (marks the location of the CA1-CA2 boundary). The inferior/internal CA2 boundary is drawn to include the SRLM. The superior boundary is drawn on the border of the external white matter to exclude it.

##### 3.5.2.4 CA3 label

The lateral boundary is the superior portion of Line 2 (marks the location of the CA2-3 boundary). The superior and medial boundary is drawn to exclude external white matter and CSF. The inferior/internal boundary is the 45° vector to the superior medial edge of the hippocampus (marks the location of the CA3-DG boundary), including any remaining visualization of the SRLM within that zone.

##### 3.5.2.5 DG label

The remainder of the internal volume that is visualized as a wedge from Line 2 and the medial 45° bisector is the DG. The superior/internal boundary is the 45° vector to the superior medial edge of the hippocampus (marks the location of the CA3-DG boundary) and excludes external white matter (fimbria) and CSF. The lateral and inferior boundaries are internal at the hippocampal fissure/SRLM.

## 4.0 Discussion

Through the efforts of multiple working groups over several years, the HSG has developed a histologically-valid, reliable, and freely available segmentation protocol for high-resolution T_2_-weighted imaging (https://hippocampalsubfields.com/harmonized-protocol/). Dozens of protocols exist to label human hippocampal subfields on MRI that proliferated over the past decade with the wide implementation of high-resolution imaging. Significant discrepancies between protocols make reconciling the current literature difficult, if not impossible. The harmonized hippocampal subfield segmentation protocol provides a solution to this barrier and can facilitate deeper insights into development and aging of the structures, their unique cognitive and functional correlates, and their vulnerability in clinical conditions (e.g., Berron et al., 2016; Daugherty et al., 2016; La Joie et al., 2010; Mueller et al., 2010; Shah et al., 2019). The HSG has been inspired to address this challenge by creating a harmonized protocol through consensus and that would be valid for scans of all ages and clinical applications.

This is a lofty task. We were not the first ones to be thus inspired and we followed the roadmap of the EADC-ADNI working group that tackled a similar challenge in definitions of total hippocampal segmentation on T_1_-weighted MRI (aka, HarP). The HarP group pioneered an approach for an “evidence-based Delphi panel” that presented data and relevant publications during the process of experts voting on landmark and boundary definitions (Boccardi et al., 2015). However, in our process we identified unique circumstance for a harmonized protocol in hippocampal subfields (Wisse et al., 2017). The foremost issue was that no existing canonical definitions of hippocampal subfields on MRI existed—the regions are traditionally defined by cytoarchitecture that can only be seen with post-mortem histological staining (Ding, 2013; Duvernoy et al., 2013; Insausti & Amaral, 2004; Williams et al., 2023). Second, there was little consistency of the anatomical nomenclature used in available MRI segmentation protocols, let alone the definitions for applying the same label. Our first investigation as a group found striking differences in boundary definitions and nomenclature across common protocols in the field that could not be directly reconciled (Yushkevich et al., 2015), and undermined the validity of our shared literature. We determined the HSG would begin by developing new definitions as opposed to voting among an existing set as the EADC-ADNI HarP procedure had (Boccardi et al., 2015). We started from ground truth by creating a novel histological reference set that had multiple neuroanatomists rating multiple images, so as to represent variability in anatomy and expert judgment in the process of developing an MRI protocol.

The Delphi procedure was efficient and reached consensus with one iteration of voting (similar to the outer boundary consensus; Olsen et al., 2019). The presentation of evidence that included the novel histological reference set in this process was key (Boccardi et al., 2015). By virtue of our international working group structure, we had an opportunity to combine many different neuroanatomical source materials, methods, and collaborating neuroanatomists to weigh in on the developing protocol. That same information was then presented to a wide representation of experts in the field to vote on the protocol. Second, we collected qualitative comments from the raters in the feasibility check and from the Delphi panelists to iteratively refine the written protocol descriptions, supporting materials, and to contextualize the consensus vote.

The initial validation of the protocol was demonstrated by the Delphi procedure and the reliability analysis. All boundary definitions had strong agreement and clarity ratings, suggesting support from the broad research community. To elicit participation in the protocol development process and the Delphi panels, we had recurring calls via our open listserv, website, social media accounts, and through numerous abstract presentations at international conferences. We have prioritized collecting expert opinions from investigators in different disciplines related to hippocampal subfield study and incorporating the feedback on accessibility of the materials from novice scholars.

All hippocampal subfield labels had high reliability. As reasonably expected, the intra-rater reliability was slightly higher than inter-rater reliability, but nonetheless all indicated for good quality measurement. Critically, the reliability metrics were similar between expert raters and the novice rater – suggesting that prior experience with manual segmentation is not a requisite to reliably apply the protocol. These results from the reliability analysis support our aim to have a protocol accessible for wide adoption, regardless of prior experience with manual segmentation or knowledge of human hippocampal anatomy. Moreover, the distributions of average *DSC* of the hippocampal subfield labels did not differ on scans collected in children as compared to adults, and among ADNI scans it did not differ as a function of cognitive impairment. An additional strength of the reliability dataset is that scans were collected at different sites on MRI machines by different manufacturers. Taken together, this evidence suggests the harmonized protocol can be reliably applied to scans age 4-94 years, with common age-related health comorbidity and dementia, and across common imaging environments to support valid comparisons across studies.

The relatively lower reliability of CA2 measures as compared to the other regions is not surprising given the small size of the region, in which even small differences between raters can have great weight in the reliability. For example, the average intra-rater reliability of the CA2 measures were excellent as compared to the modest inter-rater reliability. Although the metrics fall within an acceptable range based on previous publications (Homayouni et al., 2021; Winterburn et al., 2013), the implications for applied hypothesis testing should be carefully considered as a limitation of the protocol. The first consideration is on the quality of inference about the correlates of CA2, as the measurement error may weaken the accuracy and specificity of the label. This was a noted concern by the Delphi panelists that was a source of disagreement for some individuals (although majority agreement was achieved). Second, measurement reliability goes to the power of hypothesis tests with the data (Zimmerman & Zumbo, 2015). Because most studies will be interested in hippocampal subfield measurement to make comparisons of differential effects or functional correlates, the amount of measurement error per subfield should be considered when interpreting comparative results and determining statistical power. For research questions that are not specific to CA2, the investigator may choose to combine the region with another label to further improve the measurement reliability, or they may specify analysis *a priori* to only the subfields relevant for the hypothesis excluding this region.

Indeed, investigators could segment the hippocampal subfields by the harmonized protocol and later choose to aggregate any of the contiguous labels based on their research question. This approach may appeal to investigators applying structural masks from the T_2_-weighted images to functional MRI or multi-modal data that often use resolutions ≥ 1mm^3^, in which aggregated labels provide larger sampling areas to improve the quality of the derived measurement. Although an aggregated label loses regional specificity, it may be an acceptable compromise in the context of research that necessitates maximal measurement reliability. If the original boundary definitions and nomenclature are consistent with the protocol, the aggregated label could still be compared to other studies with the harmonized protocol—addressing the primary limitation in the current literature that originally inspired the HSG.

In some applications, though, aggregating labels may not be an acceptable compromise even for the sake of reliability. For example, this was the main critique of the Delphi panelists of the CA3-DG boundary, which on some slices of the example cases would have potential mix of tissue between the two region labels, and thus reduce potential functional specificity. The neuroanatomists generally had strong similarity in identifying the CA3 boundary, but there were discrepancies on the DG boundaries with consideration of inclusion for CA4/hilus regions. The CA3 morphometry is complicated and could be approximated more closely by the endfolial pathway rule, which in particular has been shown effective on high-field strength MRI (Berron et al., 2017). However, when adapting a similar rule in the current protocol development, the Delphi panel voted to not move the rule forward due to its complexity and weaker reliability. The cost-benefit assessment by the expert Delphi panelists emphasized that the loss to measurement reliability may not be worth the few pixels difference in accuracy.

Second only to face validity, measurement reliability is a forefront issue in application of the harmonized protocol. Because reliability is a property of both the sample and raters, and not the measurement instrument *per se*, investigators interested in applying the harmonized protocol are strongly encouraged to establish reliability in their own sample with their own raters (see online supplemental training materials^i^ for advice on implementation). As reliability is necessary for valid interpretation of the measures, the protocol reliability should be established before processing all sample data for analysis. The reliability assessment and implementation of the anterior-posterior body ranging landmarks (Olsen et al., 2019) is separate from the segmentation protocol reliability reported here; these two parts of the protocol do not require the same rater, and so the labor could be distributed in a team and implemented sequentially.

The information provided throughout the development process and Delphi panel was used to refine and enhance the clarity of the protocol description, and led to the development of a substantial set of training materials that are available to support protocol adoption. In addition to written rule descriptions with example images, including those with common artifacts or variations in morphometry, we have developed an instructional video with demonstration of manual segmentation. Example images and segmentation files in ITK-Snap are available for download from our website^i^. We are additionally offering periodic in-person sessions to provide a hands-on training experience.

The protocol can be implemented in any modern software that allows manual segmentation; for ease of use and freely available resource, we have implemented all example materials in ITK-Snap (Yushkevich et al., 2006). There are no specific requirements for the tracing environs or hardware; any idiosyncrasies specific to a laboratory are tolerable as long as reliability is confirmed to be similar to the metrics we report here. Standardization of equipment and software in the field is not required for harmonized hippocampal subfield measurements with our protocol, however it may contribute variation to measurement that we could not evaluate here. Continuing work with wider adoption of the protocol will provide opportunities to assess the effects of software and segmentation hardware (e.g., tracing tablet vs. mouse) in the future. We envision that the HSG Harmonized Protocol for the hippocampal body can be applied to existing and new high-resolution T_2_-weighted datasets and used as reference to translate current published findings to a common nomenclature to improve comparisons in qualitative and quantitative review.

### 4.1 Limitations of the Protocol and Continuing Work by the HSG

Several limitations of the protocol and continuing work should be noted. First, the boundaries drawn on MRI are approximations of the location of microstructural features that cannot be visualized on typical *in vivo* images (e.g., 0.4 × 0.4 mm^2^ in-plane resolution, collected at 3 tesla field strength). We validated the protocol with visual evaluation in comparison to labeled histological images; future *ex vivo* studies can continue to validate the protocol in reference to specific anatomical variation or disease pathology, in addition to dementia as was done here. Our continued work will apply the protocol on *in vivo* MRI scans in different clinical samples to further test convergent and divergent validity with other biomarkers of age-related neurodegeneration and dementia.

Second, the protocol was developed for an imaging sequence that assumes high in-plane coronal resolution with T_2_-weighting, and it is typically applied as an anisotropic voxel with 2-mm slice thickness. We selected this imaging protocol because it was the first available for clinical research (e.g., Iglesias et al., 2015; Mueller & Weiner, 2009; Winterburn et al., 2013; Zeineh et al., 2000), and it remains among the most popular methods in the field (e.g., see Homayouni et al., 2023 for a summary in a recent meta-analysis). Based on the HSG’s prior review, sub-millimeter in-plane coronal resolution with T_2_-weighting is the minimum required to visualize SRLM and other key landmarks to segment hippocampal subfields (Wisse et al., 2017), and we do not recommend applying the protocol to lower-resolution (e.g., 1 mm^3^) T_1_-weighted images (Wisse et al., 2021). Although our validated protocol is designed to be implemented with these imaging restrictions, it presented limitations to the set of structures that could be reliably demarcated and excluded separate labels for CA4/hilus and the SRLM. The white matter structures at the external surface of the hippocampus - alveus and fimbria - were also excluded. As the field continues to rapidly develop new high-resolution imaging methods, including high-field strength clinical imaging, the protocol can be expanded in the future to divide out these additional labels. Additional precision may be gained with high-resolution isotropic voxels that allow applying the segmentation rules in multi-plane views. In a similar regard, advances in post-mortem histological methods continue to improve our understanding of hippocampal micro- and macro-anatomy (Ding & Van Hoesen, 2015; Palomero-Gallagher et al., 2020; Williams et al., 2023). We designed the protocol anticipating these possible future developments so that labels and contiguous boundary definitions could remain, with new labels further subdividing the structure. This can provide some continuity in the field while keeping the protocols relevant to the state-of-the-science.

Third, this portion of the harmonized protocol is designed to be applied within the hippocampal body only. The protocol is not designed for labels in the hippocampal head or tail, which will have additional labels currently under development by HSG working groups. The body is the largest portion of the hippocampus (Daugherty et al., 2015; Malykhin et al., 2017; Poppenk et al., 2013), and there are several examples of current protocols that exclusively measure the body as a representative measure (e.g., Bender et al., 2018; Mueller & Weiner, 2009; see Yushkevich et al., 2015 for comparisons). Until all definitions have been developed and published, assessment of the subfields within the hippocampal body is feasible. However, such measurement should be noted as an estimate representative of only that portion of the hippocampus.

### 4.2 Summary

Through collaborative working groups and Delphi consensus procedures, the HSG has developed a harmonized protocol for subfield segmentation in the hippocampal body. We have validated the protocol with multi-site and multi-manufacturer imaging data from healthy 4-94 years olds, and people with cognitive impairment. In complement to strong face validity of the protocol compared to a novel histological reference set, the high reliability of the protocol is the HSG’s contribution to support well-powered MRI studies with feasible sample sizes (Homayouni et al., 2021). The protocol is available for immediate adoption and application to existing and new high-resolution, T_2_-weighted datasets. The harmonized protocol for the hippocampal body can be adopted immediately for manual segmentation, and the HSG is currently developing an automated segmentation atlas as well. Our ongoing work is following the same procedures reported here to provide future updates to the harmonized protocol with hippocampal subfield labels in the head and tail, and labels for medial temporal lobe cortices, all developed from detailed parcellations of histology by neuroanatomists. Future HSG studies will apply the harmonized protocol to clinical samples for further validation against established biomarkers of neurodegenerative disease and dementia risk.

## Resource and Data Sharing Statement

^i^All supporting data and resource materials are available for download from the Hippocampal Subfield Group (HSG) website: https://hippocampalsubfields.com/harmonized-protocol/.

## Acknowledgements

In memory of Dr. Ricardo Insausti, a dear friend and colleague who shared his expertise and love of the hippocampus with this group.

This work was supported NIH/NIA R01-AG070592 (multi-site award; lead PI L. Wang); NIH/NIA P30-AG072931 (A.M. Daugherty); NIH/NIA R01-AG011230 (A.M. Daugherty and N. Raz); NIH/NICHD F32-HD108960 (K.L. Canada); NIH/NIA R01-AG073250 (T. Brown); NIH/NIA R01-AG072056 (J. Augustinack); NIH/NIA K99/R00-AG065457 (L. Pasquini); NIH/NIMH R01-MH107512 (N. Ofen); NIH/NICHD R01-HD079518 (T. Riggins); NIH/NIA P30-AG066519 (C. Stark); NIH/NIA RF1-AG056014 and R01-AG069474 (P. Yushkevich); European Union’s Horizon Europe Programme under the Specific Grant Agreement No. 101147319 (EBRAINS 2.0 Project), Helmholz Association’s Initiative and Networking Fund through the Helmholz International BigBrain Analytics and Learning Laboratory (HIBALL) under the Helmholz International Lab grant agreement InterLabs-0015, and the Joint Lab Supercomputing and Modeling for the Human Brain (all to K. Amunts); European Union’s Horizon 2020 research and innovation programme (grant agreement number 667696), Fondation Alzheimer, Agence Nationale de la Recherche, Région Normandie, Fondation Vaincre Alzheimer, Fondation Recherche Alzheimer and Fondation pour la Recherche Médicale (all to G. Chetelat); Canadian Institutes of Health Research (CIHR) PJT-162292 (R.K. Olsen); MultiPark - A Strategic Research Area at Lund University (L. Wisse).

Portions of the included MRI scans included in this project was funded by the Alzheimer’s Disease Neuroimaging Initiative (ADNI) (National Institutes of Health Grant U01 AG024904) and DOD ADNI (Department of Defense award number W81XWH-12-2-0012). ADNI is funded by the National Institute on Aging, the National Institute of Biomedical Imaging and Bioengineering, and through generous contributions from the following: AbbVie, Alzheimer’s Association; Alzheimer’s Drug Discovery Foundation; Araclon Biotech; BioClinica, Inc.; Biogen; Bristol-Myers Squibb Company; CereSpir, Inc.; Cogstate; Eisai Inc.; Elan Pharmaceuticals, Inc.; Eli Lilly and Company; EuroImmun; F. Hoffmann-La Roche Ltd and its affiliated company Genentech, Inc.; Fujirebio; GE Healthcare; IXICO Ltd.; Janssen Alzheimer Immunotherapy Research & Development, LLC.; Johnson & Johnson Pharmaceutical Research & Development LLC.; Lumosity; Lundbeck; Merck & Co., Inc.; Meso Scale Diagnostics, LLC.; NeuroRx Research; Neurotrack Technologies; Novartis Pharmaceuticals Corporation; Pfizer Inc.; Piramal Imaging; Servier; Takeda Pharmaceutical Company; and Transition Therapeutics. The Canadian Institutes of Health Research is providing funds to support ADNI clinical sites in Canada. Private sector contributions are facilitated by the Foundation for the National Institutes of Health (www.fnih.org). The grantee organization is the Northern California Institute for Research and Education, and the study is coordinated by the Alzheimer’s Therapeutic Research Institute at the University of Southern California. ADNI data are disseminated by the Laboratory for Neuro Imaging at the University of Southern California.

## Conflict of Interest Statement

The authors have no conflicts of interest to disclose.

^i^ Supplemental content, materials to support protocol training and adoption, and complete results from the Delphi procedure are available on the Hippocampal Subfields Group website: https://hippocampalsubfields.com/harmonized-protocol/. Please refer to the website for updates and additional protocol developments by the Hippocampal Subfields Group.

## Notes

https://hippocampalsubfields.com/harmonized-protocol/

### Competing Interest Statement

The authors have declared no competing interest.

