## Supplemental Information for "Harmonized Protocol for Subfield Segmentation in the Hippocampal Body on High-Resolution *in vivo* MRI from the Hippocampal Subfields Group (HSG)"

Proposed Protocol for Subfield Boundaries within the Hippocampal Body

On High-Resolution T2-weighted Images Collected at 3 Tesla

On behalf of the Boundary Working Group,

Hippocampal Subfield Segmentation Group

hippocampalsubfields.com

Acknowledgments and working group members: Ana Daugherty, David Berron, Valerie Carr, Robin de Florès, Nikolai Malykhin, Rosanna Olsen, and Samaah Saifullah

This document provides supplemental information for the evaluation of the proposed draft protocol to segment subfields within the hippocampal body. Please do not implement this information in your own research or disseminate outside of the HSG until the protocol is evaluated and published.

Appendix A: Protocol Training Documentation .......... i
Appendix B: Description of Outer Boundaries .......... viii
Appendix C: Example Tracing Information and Anterior-Posterior Ranges .......... xxx
Appendix D: Brief Introduction to ITK Snap for Applying Manual Segmentation Protocol .......... xxxii

### Geometric Heuristic

<u>Geometric Heuristic Description:</u>
Boundaries within the hippocampal body are determined by a geometric heuristic that approximates the location of anatomical landmarks observable in histological samples. The steps that are required to execute this protocol are described in detail in Figure 1 below.

| | | |
|---|---|---|
| 1. Line 1 should be anchored at the opening of the hippocampal fissure adjacent to the superior edge of the subiculum (**a,** the "arm pit") and extended to the most lateral, outside edge of the alveus (white matter structure) of the CA1 sector. | 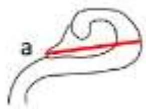  | 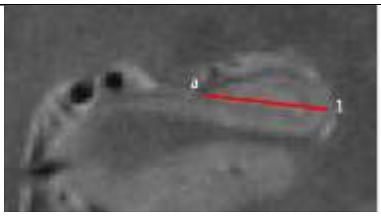 |
| 2. Line 2 is then placed perpendicular to the middle of Line 1, extending from the most superior edge of the hippocampus to the parahippocampal white matter. | 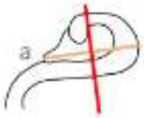  | 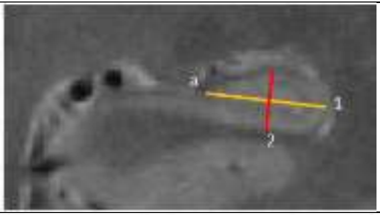 |
| 3. An additional vector is extend from the point of bisection at 30° to the lateral side. | 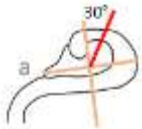  | 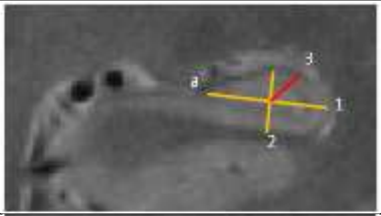 |
| 4. Extend another vector at 45° to the medial side. | 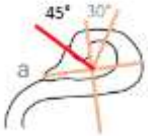  | 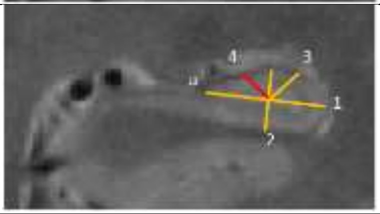 |

Fig 2. Example MR image (0.42 x 0.42 $mm^2$ in plane) with the geometric heuristic illustrated with subfield labels.

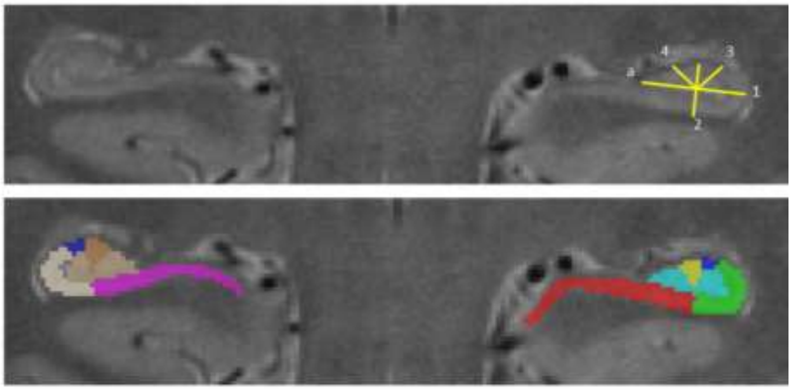

The subiculum-CA1 boundary is the inferior portion of Line 2, spanning from the SRLM to the white matter inferior to the hippocampus. The superior portion of Line 2 marks the location of the CA2-3 boundary. The 30° vector to the superior lateral edge of the hippocampus marks the location of the CA1-CA2 boundary, and the 45° vector to the superior medial edge of the hippocampus marks the location of the CA3-DG boundary. The remainder of the volume is the dentate gyrus region.

The SRLM defines the internal boundary of the CA1 and CA2 subfields. The hippocampal fissure/SRLM defines the internal boundary of the dentate gyrus. The subiculum, CA1 and CA2 subfields are drawn to include the SRLM, and it is excluded from the dentate gyrus. The medial boundary of the dentate gyrus is the CSF, and the superior boundaries are the CA3 and fimbria (visualized as hypointense on a T2-weighted image). The CA3-DG boundaries are defined by the wedge formed from Line 2 and the medial 45° bisector.

*Limitations.* Several limitations of the protocol should be noted. First, the boundaries drawn on MRI are approximations of the location of microstructural features that cannot be visualized on typical *in vivo* images (e.g., 0.4 x 0.4 $mm^2$ in-plane resolution, collected at 3 Tesla field strength). Second, the internal boundary of the dentate gyrus depends on visualization of the SRLM; the protocol has not been validated when applied to images with poor contrast. Third, this portion of the protocol is designed to be applied within the hippocampal body on images collected approximately perpendicular to the hippocampal long axis. The protocol has not been tested for application to images collected in a different orientation or in the hippocampal head or tail. Additional definitions are under development for the same subfield labels with the anterior and posterior hippocampal regions. Until all definitions have been developed and published, a limited assessment of the subfields within the hippocampal body is feasible. However, such measurement should be noted as an estimate representative of only that portion of the hippocampus.

*Feasibility Assessment.* The protocol was developed with the intent that it could be applied to brains imaged from different populations (e.g., children, healthy aging, Alzheimer's disease, epilepsy). A consideration was also made for typical imaging artifacts (e.g., motion, poor gray-white matter contrast) and variability in morphometry (e.g., round hippocampus shape vs. canonical shape) that occurs between persons, possibly in correlation with development or disease progression, and along the anterior-posterior axis. All of these sources of variability were represented in the feasibility data set.

Two expert raters who were naïve to the protocol prior to training participated in the feasibility assessment. Training included detailed documentation with example image tracings, a 2-hour introductory training session (via Skype), followed by prescribed practice and then an additional 1-3 hours of individualized feedback (via Skype).

Raters had good agreement and overlap in CA1, CA2, CA3 and dentate gyrus labels (N = 6). Based upon review of the tracings, the poor intra-class correlation agreement in total subiculum volume is due to variability in the outer boundaries.

| Subfield | Rater 1, Bilateral Average Volume Across Cases (mm$^3$' M ± SD) | Rater 2, Bilateral Average Volume Across Cases (mm$^3$; M ± SD) | ICC(2) | Average Dice (M ± SD) |
|---|---|---|---|---|
| Subiculum | 352.58 ± 66.76 | 464.78 ± 62.37 | 0.45 | 0.76 ± 0.07 |
| CA1 | 235.12 ± 56.90 | 257.22 ± 61.93 | 0.94 | 0.82 ± 0.06 |
| CA2 | 27.32 ± 6.95 | 24.64 ± 5.73 | 0.83 | 0.61 ± 0.10 |
| CA3 | 50.85 ± 13.41 | 42.56 ± 11.03 | 0.84 | 0.69 ± 0.07 |
| Dentate gyrus | 236.73 ± 50.87 | 242.03 ± 54.52 | 0.99 | 0.79 ± 0.06 |

All segmentations were completed throughout the length of the hippocampal body, as determined by the anterior-posterior landmarks (see outer boundary protocol description). Example segmentations on three slices of the hippocampal body on one MRI scan are shown. Please download the example segmentation files to review in ITK Snap for further detail.

Fig 3. Segmentations on example MRI by two raters from the feasibility assessment.

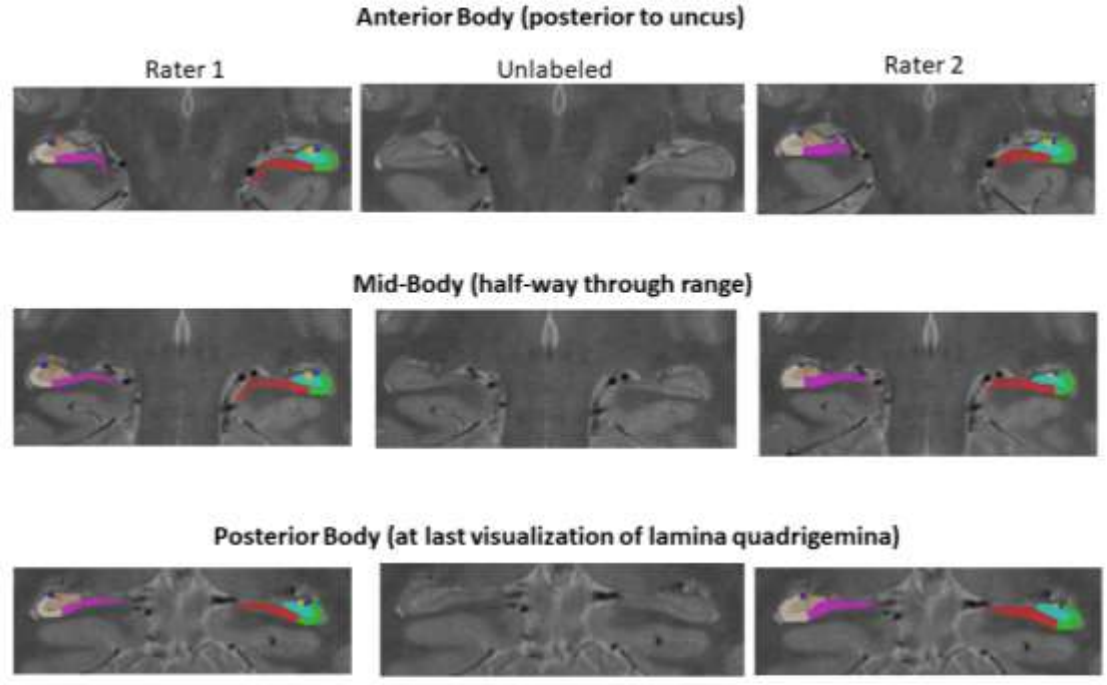

*Rater Experience*. When asked on a 7-point scale (1—Not at All, 7—Extremely Well), the raters indicated that they understood the protocol well (rater 1 = 6, rater 2 = 7) and that the rules were clearly defined in the training documents (rater 1 = 6, rater 2 = 7). When asked about the overall difficulty of performing the tracings on a 7-point scale (1—Very Easy, 7—Very Difficult), rater 2 indicated it was difficult (response = 6) and rater 1 indicated it was somewhat easy (response = 3).

When asked, both raters indicated that they had a similar experience applying the protocol on anterior and posterior slices. Both raters reported that it was more difficult to apply the protocol on a hippocampus with unusual shape, but they did not believe it biased decisions made during tracing. Both raters indicated that images with poor contrast that compromised visualization of the SRLM made applying the rules more difficult, and one rater believed this introduced bias when tracing whereas the other thought no bias occurred.

> One rater elaborated: “It was more difficult to apply the rules on image with poor contrast and I believe it introduced bias when the SLRM is blurred and visualized thicker, which influences boundaries between CA regions and DG.”
>
> The other explained, “In general, N/A, although there were a couple of instances where I felt some boundaries looked a little "off"...”

Subiculum-CA1 Boundary Histology Information and Reference Material

The geometric heuristic definitions were developed taking into account the variability between HSG neuroanatomists' labeling, variability within-person along the anterior-posterior axis, and variability between persons. The proposed definitions aim to fall approximately in the middle of the range of variability (i.e. to "split the difference"). Here we describe the variability we observed.

Comparing MRI protocols for labeling subfields, we noted the most disagreement between protocols in the placement of the subiculum-CA1 boundary (Yushkevich et al., 2015); and this boundary is also the most discrepant between neuroanatomists. Further, we observed notable individual variability in the subiculum-CA1 boundary within an individual along the anterior-posterior axis and between individuals. Example illustrations are shown below, please refer to the p. 27-39 of the online supplement for a complete depiction of the histological reference set. We note several differences in labeling protocols between neuroanatomists, including the choice to label pro-subiculum, which for the purpose of developing a MRI protocol, pro-subiculum was considered a part of the subiculum label.

Fig 4. Comparison of three HSG neuroanatomists labeling the same slice of the same brain (post-mortem image with Nissl stain). Subiculum-CA1 boundary is indicated by red arrow.

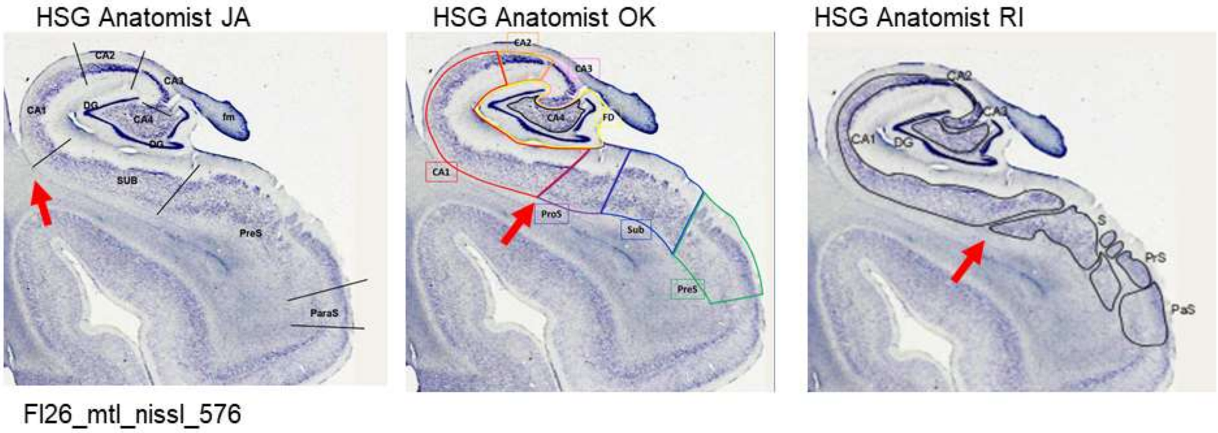

The variability between neuroanatomists reflect differences in protocols as well as differences in informed opinion, similar to what we observe when comparing MRI protocols. The variability observed between HSG neuroanatomists falls within the range of individual differences in anatomy reported in a recent publication (Zeineh et al., 2015).

Fig 5. A portion of Figure 3 from Zeineh et al. (2015, Neurobiol Aging) of two different example brain slices with Nissl stain is shown in comparison to the HSG neuroanatomists' notations (marked with red arrow). Note that Zeineh et al. (2015) are illustrating individual differences in the (pro)subiculum-CA1 boundary and the yellow lines indicate the range in which the boundary may fall on that specific slice in cross-reference to other anatomical information and histological methods (see supplement and original publication for examples of other information referenced). The variability observed between HSG neuroanatomists falls within the range of variability in anatomy observed between individuals reported by Zeineh et al. (2015).

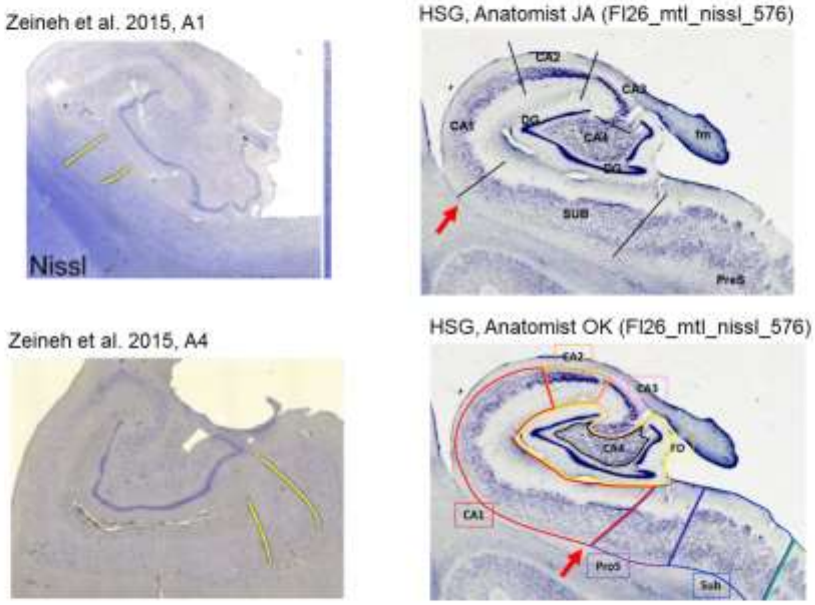

Completing a similar exercise, one of the HSG neuroanatomists (OK) illustrated the range in which the subiculum-CA1 boundary may fall on an anterior and posterior example image, including additional notation of the anatomical landmarks she referenced in making the decision.

Fig 6. The range in which the (pro)subiculum-CA1 boundary may fall on a single slice as labeled by HSG neuroanatomist (OK). The straight yellow lines indicate the range in which the boundary may fall and additional landmarks are labeled that were used in making the notations. Images are high-resolution MRI (0.2 × 0.2 × 0.2 mm) scans of a post-mortem sample obtained from a 61-year old with semantic variant dementia with primary progressive aphasia, acquired approximately perpendicular to the AC-PC axis. Left image: anterior hippocampal body (0.8mm beyond the end of the uncal apex). Right image: posterior hippocampal body (at the level of the lateral geniculate nucleus).

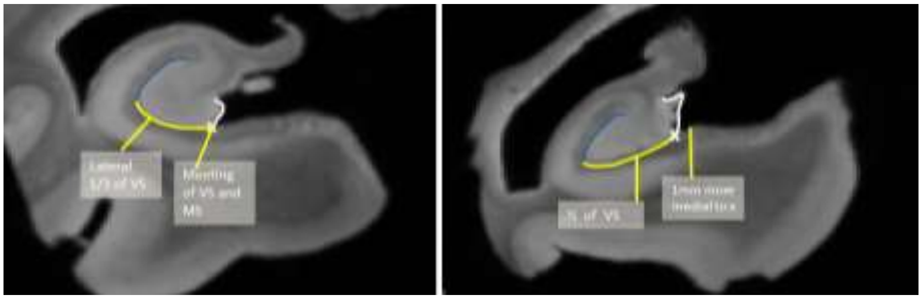

Consistent with the available literature (e.g., Ding, 2013; Zeineh et al., 2015), we note variability in the subiculum-CA1 boundary between individuals when compared at approximately the same anatomical level (Fig #) and within an individual along the anterior-posterior axis (Fig #). For additional illustrations, please refer to the complete histological reference set (p. 22-39).

Fig 7. Comparison of two individuals at approximately the same anatomical level (posterior to the uncus), labeled by the same HSG neuroanatomist. The subiculum-CA1 boundary is indicated with the red arrow.

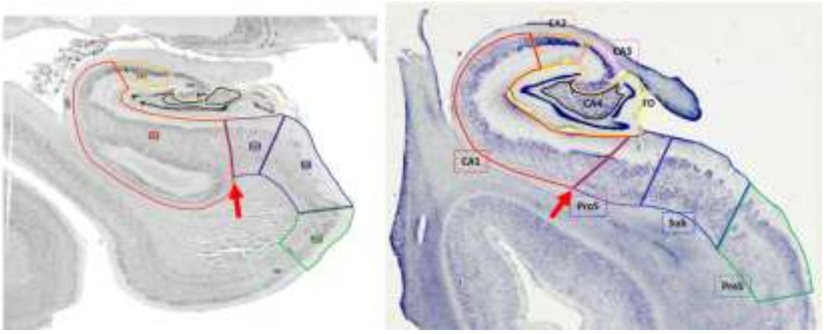

Fig 8. Comparison of the subiculum-CA1 boundary along the length of the hippocampal body, and between individuals at approximately the same anatomical level along the anterior-posterior axis. The first (left) image of each series is in the anterior body (posterior to the uncus) and images are shown with approximately 4mm gaps. The subiculum-CA1 boundary is indicated with a red arrow and all images are labeled by the same HSG neuroanatomist.

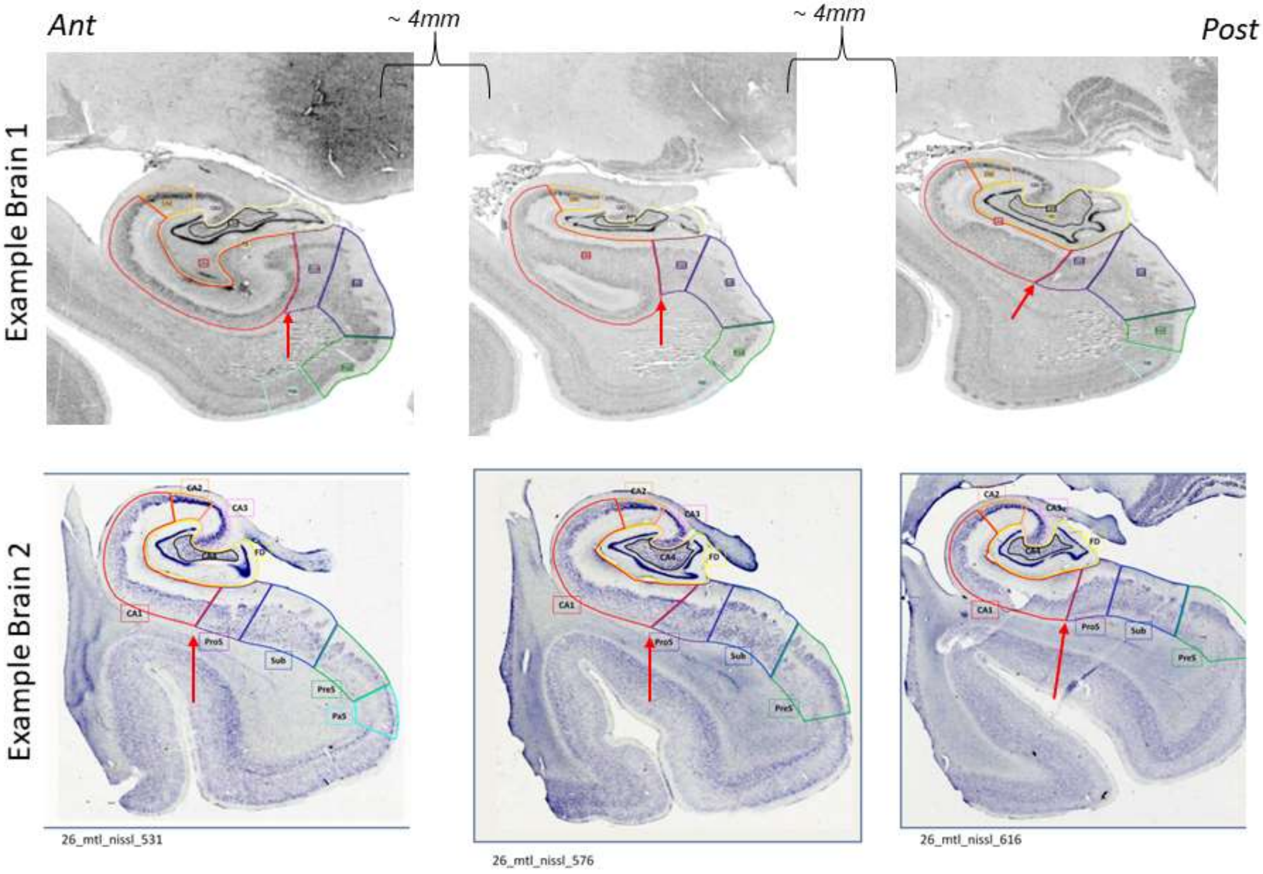

#### Subiculum-CA1 Boundary Definition Evaluation

*Rater Experience.* Example segmentations on multiple slices of one MRI dataset by the two raters in the feasibility attempt are provided. Please refer to the online supplemental materials for multiple examples of segmented MRI datasets.

When asked about the difficulty of each boundary when segmenting according to the geometric heuristic, rated on a 7-point scale, both raters indicated the subiculum-CA1 boundary was easy (rating = 2). The outer boundary of the medial subiculum-cortex was reported to be relatively more difficult by both raters (rating = 4).

*Validation.* As a step towards assessing construct validity, the geometric heuristic was compared to the notations by the HSG neuroanatomists on post-mortem histology images. Examples of the visual comparisons are shown on a slice from two different brains (see Section # for complete annotated data). The proposed boundary appears to fall within the range of variability noted from multiple sources.

Fig 9. Visual comparison of the subiculum-CA1 boundary between three neuroanatomists (blue green and yellow lines) and the geometric heuristic (red line). Two different brains (Boston and Philadelphia samples) are depicted in the panel.

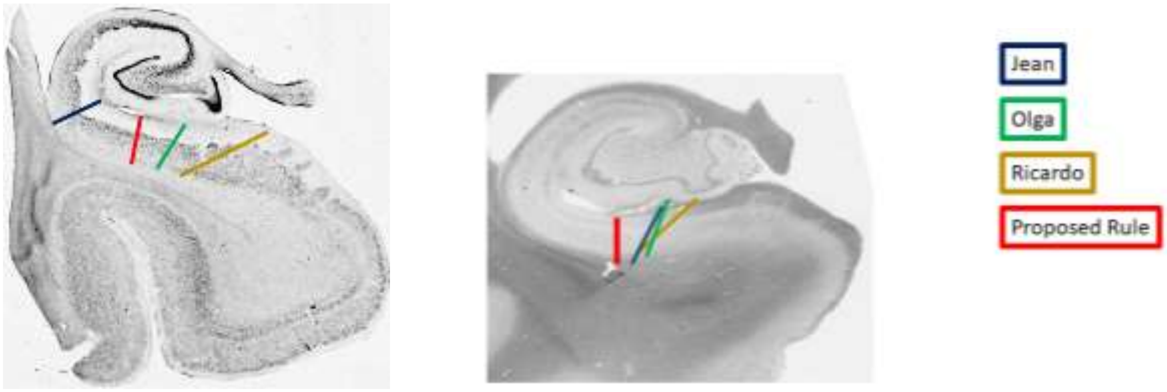

### CA1-CA2 Histology Information and Reference Material

As compared to the subiculum-CA1 boundary, we observed less variability in the CA1-CA2 boundary. Nonetheless, there were some differences between HSG neuroanatomists, between individuals, and within-individual along the anterior-posterior axis.

Fig 10. Comparison of 3 HSG neuroanatomists labeling the same slice of the same brain (post-mortem image with Nissl stain). CA1-CA2 boundary is indicated by red arrow.

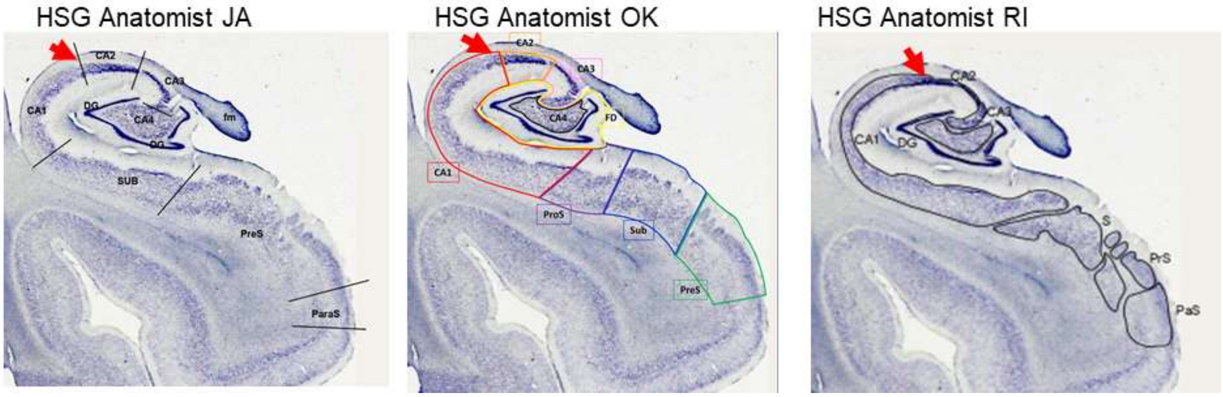

Fig 11. Comparison of the CA1-CA2 boundary along the length of the hippocampal body, and between individuals at approximately the same anatomical level along the anterior-posterior axis. The first (left) image of each series is in the anterior body (posterior to the uncus) and images are shown with approximately 4mm gaps. The CA1-CA2 boundary is indicated with a red arrow and all images are labeled by the same HSG neuroanatomist.

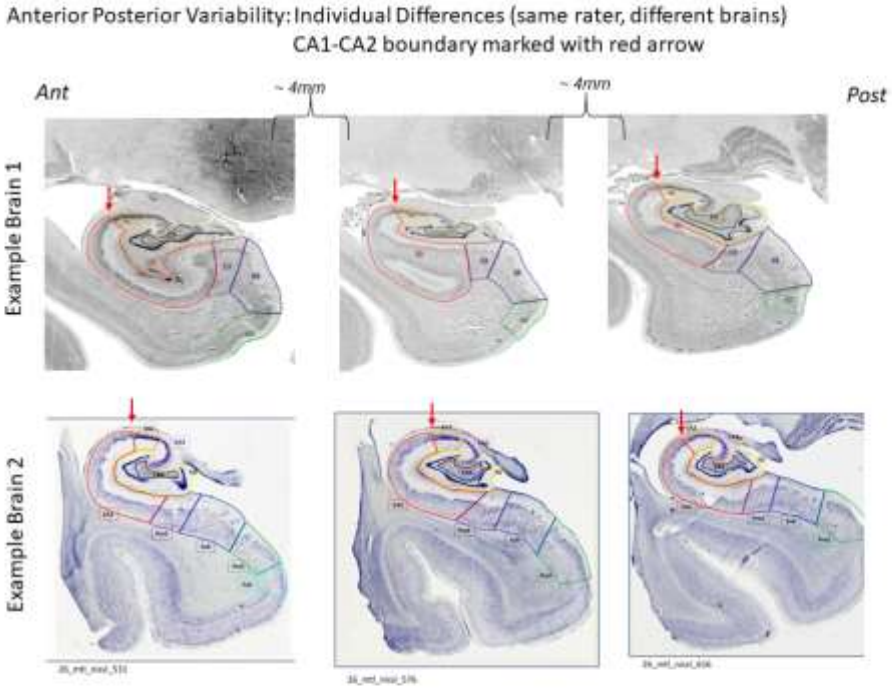

CA1-CA2 Boundary Definition Evaluation

*Rater Experience.* When asked, rater 2 indicated that this boundary was “very easy” to segment, and rater 1 reported it was “easy”.

*Validation.* The proposed boundary appears to fall within the range of variability in the CA1-2 boundary noted from multiple sources.

Fig 12. Visual comparison of the CA1-CA2 boundary between three neuroanatomists (blue, green, and yellow lines) and the geometric heuristic (red line). Two different brains (Boston and Philadelphia samples) are depicted in the panel.

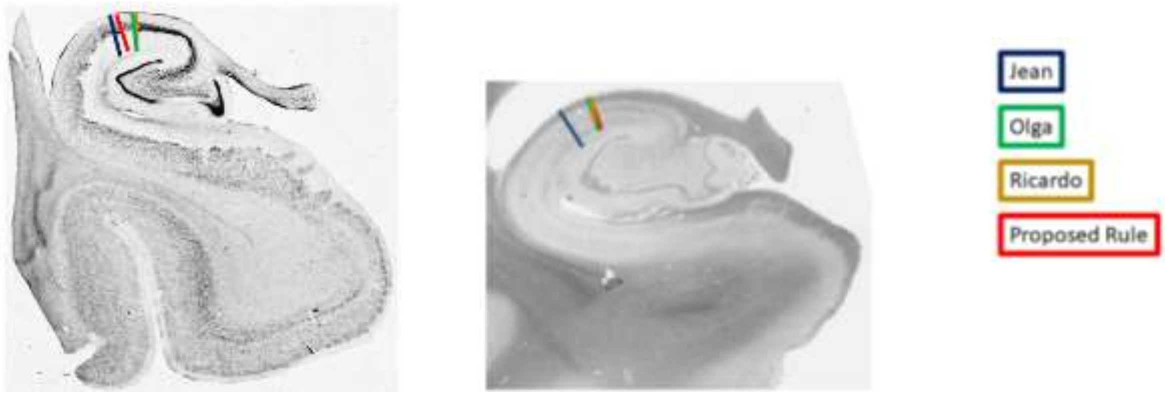

CA2-CA3 Histology Information and Reference Material

Variability was observed in the CA2-CA3 boundary definition between neuroanatomists, as well as within- and between-person differences.

Fig 13. Comparison of 3 HSG neuroanatomists labeling the same slice of the same brain (post-mortem image with Nissl stain). CA2-CA3 boundary is indicated by red arrow.

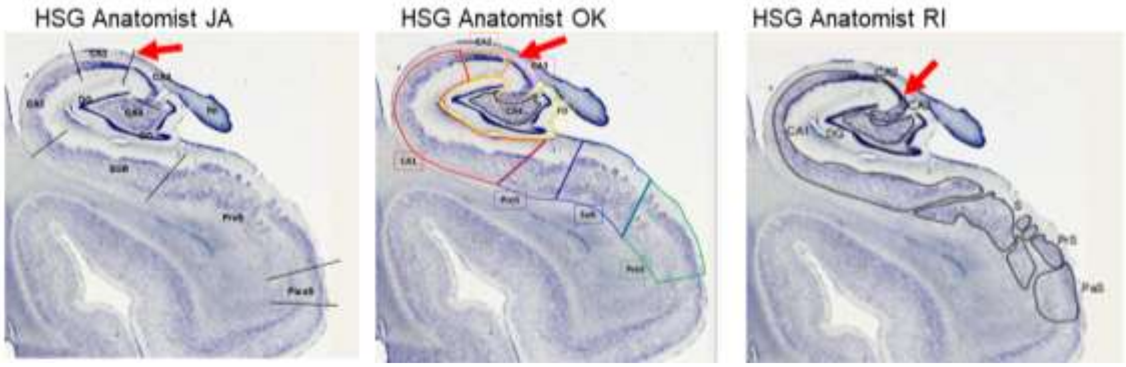

Fig 14. Comparison of the CA2-CA3 boundary along the length of the hippocampal body, and between individuals at approximately the same anatomical level along the anterior-posterior axis. The first (left) image of each series is in the anterior body (posterior to the uncus) and images are shown with approximately 4mm gaps. The CA2-CA3 boundary is indicated with a red arrow and all images are labeled by the same HSG neuroanatomist.

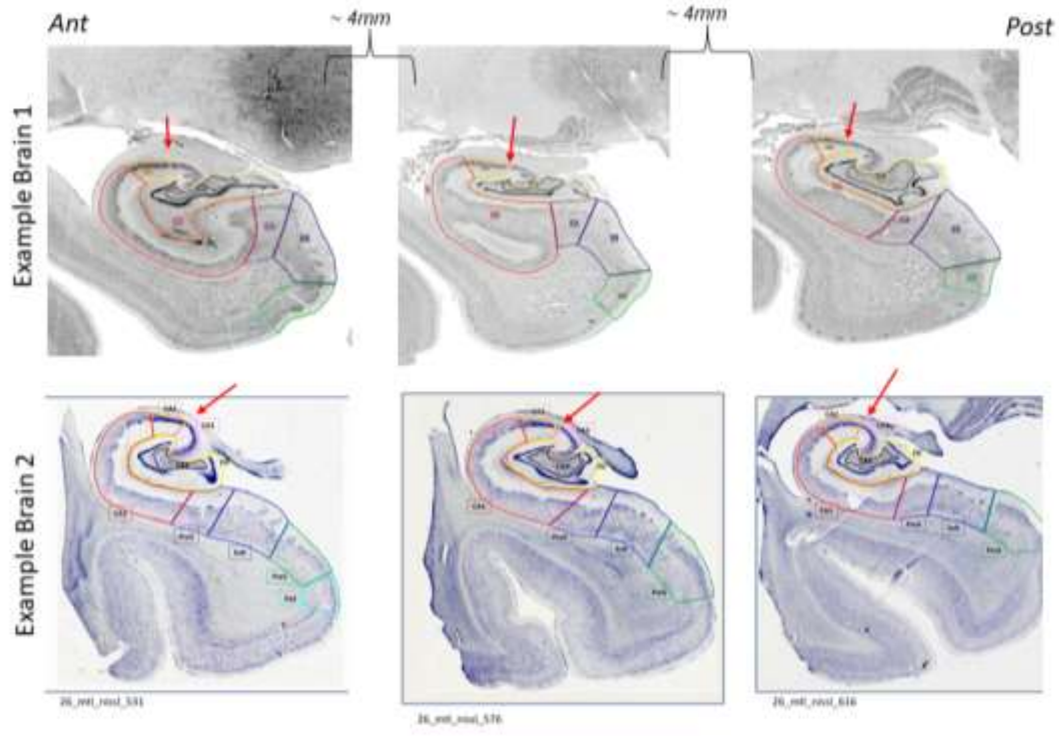

CA2-CA3 Boundary Definition Evaluation

*Rater Experience.* When asked, rater 2 indicated that this boundary was “very easy” to segment, and rater 1 reported it was “easy”.

*Validation.* The proposed boundary appears to fall within the range of variability noted from multiple sources. However, we note that CA2 is smallest subfield and therefore even small deviation in the boundary may have large consequences on the resulting measurement. The CA2 subfield volume estimate had good agreement between raters (ICC(2) = 0.83) but had the smallest amount of overlap between raters (average Dice = 0.61 $\pm$ 0.10).

Fig 15. Visual comparison of the CA2-CA3 boundary between three neuroanatomists (blue, green, and yellow lines) and the geometric heuristic (red line). Two different brains (Boston and Philadelphia samples) are depicted in the panel.

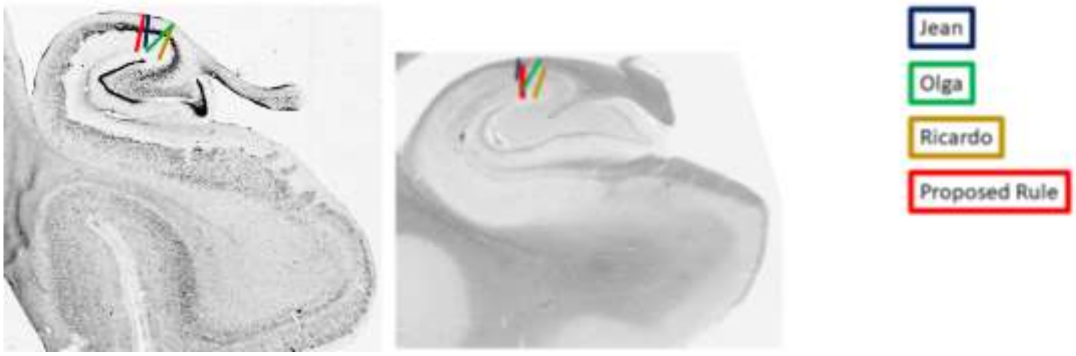

### CA3-Dentate gyrus Histology Information and Reference Material

Definitions of the CA3-dentate gyrus boundaries were overall more consistent between neuroanatomists, but they differed in the decision to label CA4 (or hilus). For the purpose of developing a protocol for MRI, area CA4 was included in the dentate gyrus label.

Fig 16. Comparison of three HSG neuroanatomists labeling the same slice of the same brain (post-mortem image with Nissl stain). CA3-dentate gyrus boundary is indicated by bolded, bright red line.

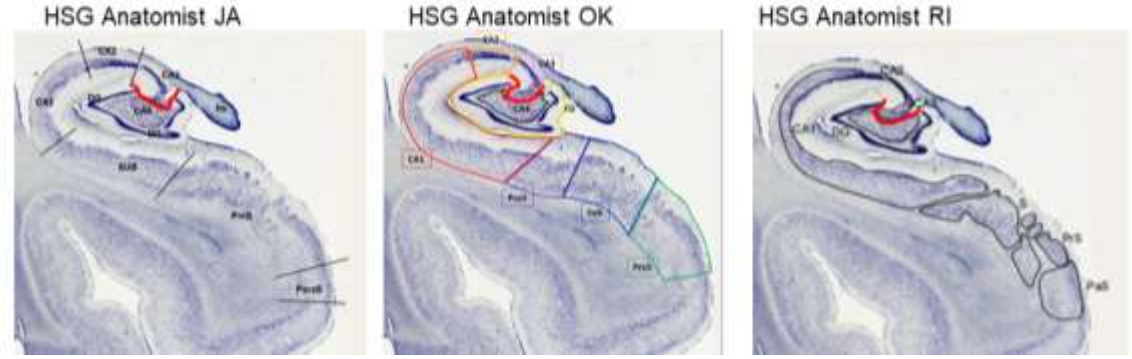

Fig 17. Comparison of the CA3-dentate gyrus boundary along the length of the hippocampal body, and between individuals at approximately the same anatomical level along the anterior-posterior axis. The first (left) image of each series is in the anterior body (posterior to the uncus) and images are shown with approximately 4mm gaps. The CA3-dentate gyrus boundary is indicated by the bold, bright red line and all images are labeled by the same HSG neuroanatomist.

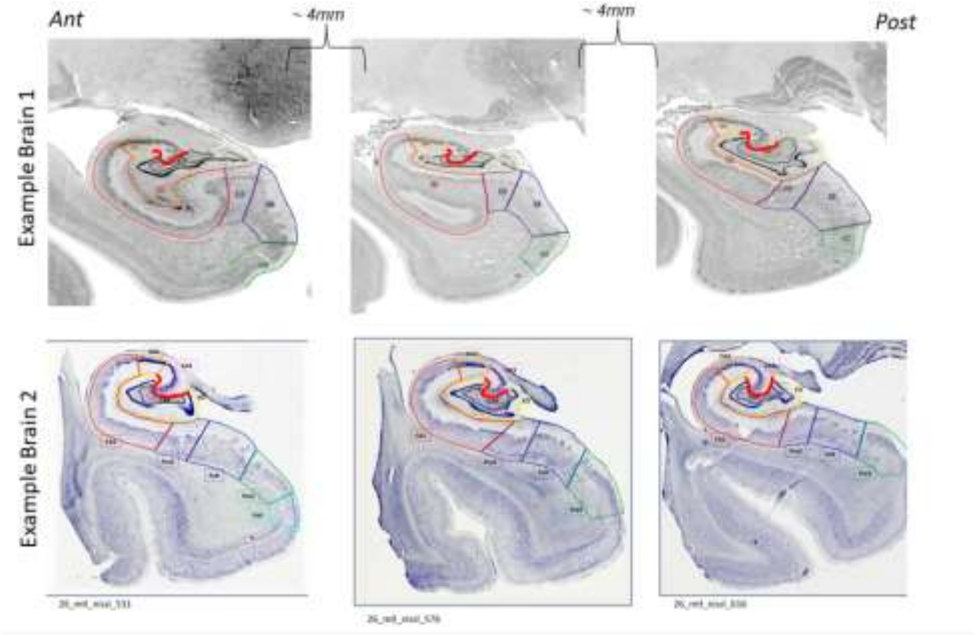

Geometric Heuristic, CA3-Dentate gyrus Boundary Definition Evaluation

*Rater Experience.* When asked, rater 2 indicated that the dentate gyrus boundaries were "very easy" to segment, and rater 1 reported it was "easy".

*Validation.* The proposed boundary appears to overlap with the CA3 region, but may include some of the dentate gyrus in the label. Conversely, the dentate gyrus label is considered to exclude the CA3.

Fig 18. Visual comparison of the CA3-dentate gyrus boundary between three neuroanatomists (blue, green, and yellow lines) and the geometric rule (red line). Two different brains (Boston and Philadelphia samples) are depicted in the panel.

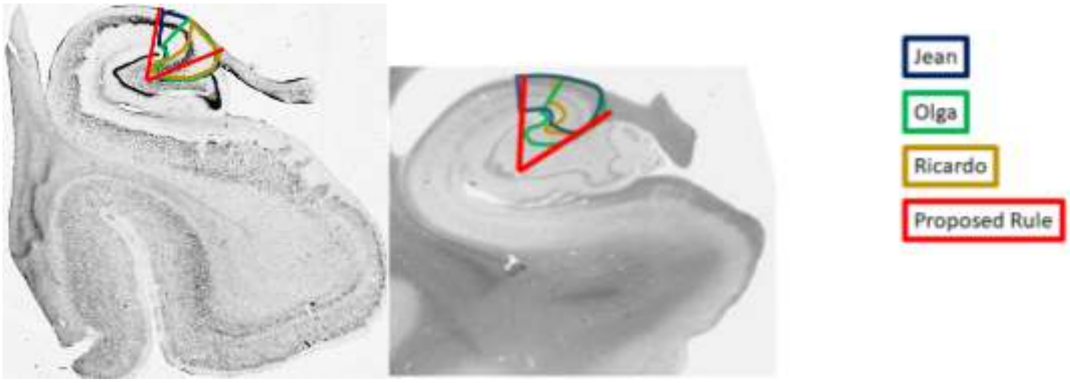

#### Endfolial Rule, CA3-Dentate Gyrus Boundary Evaluation

Observing that the CA3 label in the geometric heuristic overlapped with the dentate gyrus, an alternate rule was developed specifically for the CA3-dentate gyrus boundaries. The alternate rule approximates the location of the endfolial pathway landmark. Note that the subiculum-CA1, CA1-CA2 and CA2-CA3 boundaries are the same. This alternate rule employs a different geometric heuristic that would be applied after labeling those subfields.

<u>Endfolial Rule Description:</u>
The modified rule uses the same definitions for the subiculum-CA1, CA1-CA2 and CA2-CA3 boundaries (Line 1-3), and a different definition for demarcating the CA3-dentate gyrus boundaries (lines 4e-6e), summarized in Figure 19 below.

| | | |
|---|---|---|
| **1.** Line 1 should be anchored at the opening of the hippocampal fissure adjacent to the superior edge of the subiculum (**a,** the "arm pit") and extended to the most lateral, outside edge of the alveus (white matter structure) of the CA1 sector. | 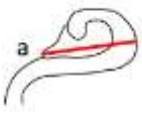  | 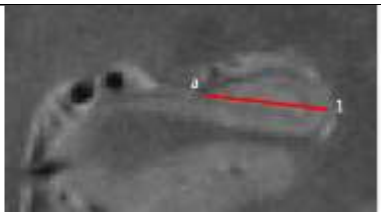 |
| **2.** Line 2 is then placed perpendicular to the middle of Line 1, extending from the most superior edge of the hippocampus to the parahippocampal white matter. | 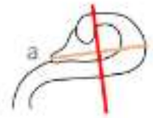  | 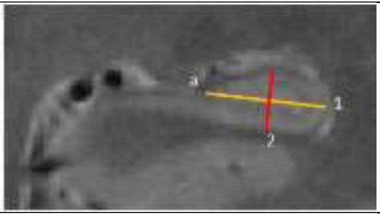 |
| **3.** An additional vector is extend from the point of bisection at 30° to the lateral side. | 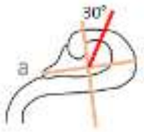  | 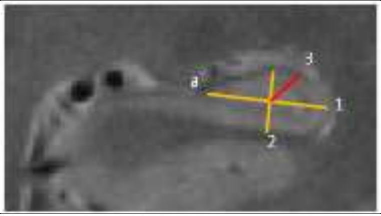 |
| **4e.** Extend Line 4e from the most medial point of the dentate gyrus (placed adjacent to the subiculum) to the most superior, middle point of the CA fields. Note that depending upon the morphometry of the hippocampus, the dentate gyrus may extend medial to the opening of the hippocampal fissure (i.e., not the same position as **a**, "the armpit"). | 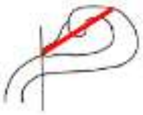 | 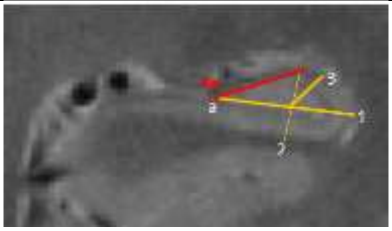 |

| | | |
|---|---|---|
| **5e.** Find the half point of Line 4e; extend Line 5e from the most lateral point of the SLRM (internal boundary between dentate gyrus and CA1) to the medial edge of the dentate gyrus, bisecting the half point of Line 4e. Ignore cysts for this line placement. | 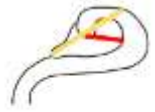 | 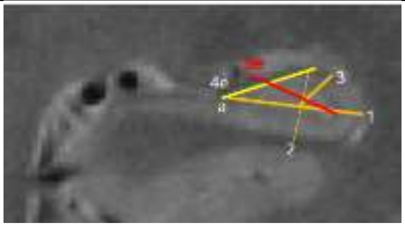  |
| **6e.** Find the half point of Line 5e, extend Line 6e from the half point to the superior edge of CA, aligned perpendicular to Line 5e. | 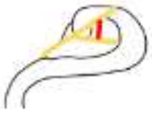 | 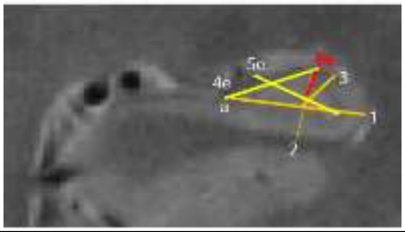  |
| Lines 5e and 6e create a wedge to the superior medial edge, and this is labeled CA3. Note that the CA2 label should not be changed and CA3 is labeled in the adjacent tissue. The remainder of the volume is dentate gyrus. | 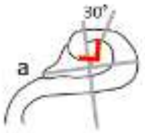  | 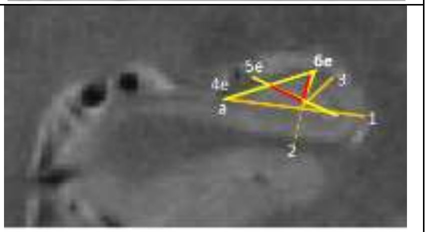  |

Fig 20. Example MR image (0.42 x 0.42 $mm^2$ in plane) with the endfolial rule illustrated with subfield labels.

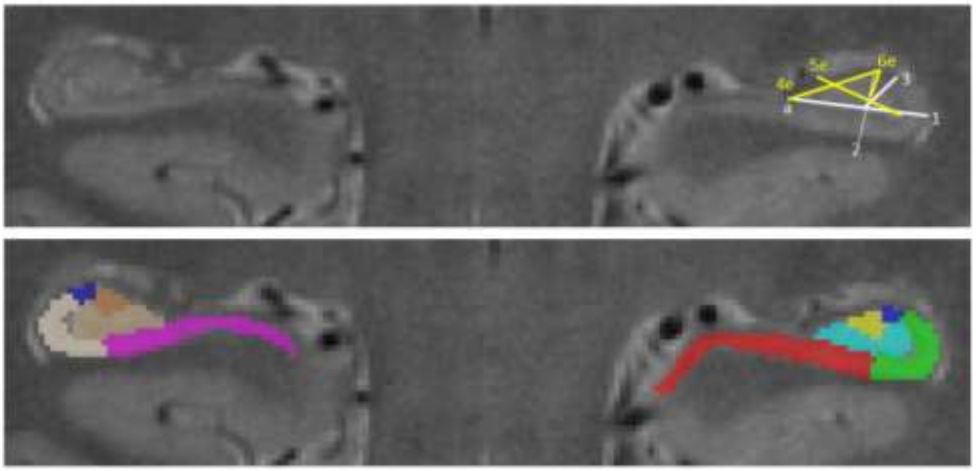

*Rater Reliability and Example Tracings.* The reliability of the CA3 subfield labeled with the endfolial rule was low, although based on feedback from the raters we believe that reliability could be improved with additional training and answering specific questions about the protocol.

| **Subfield** | **Rater 1, Bilateral Average Volume Across Cases (mm$^{3}$’ M $\pm$ SD)** | **Rater 2, Bilateral Average Volume Across Cases (mm$^{3}$; M $\pm$ SD)** | **ICC(2)** | **Average Dice (M $\pm$ SD)** |
|---|---|---|---|---|

| | | | | |
|---|---|---|---|---|
| CA3 | 53.36 ± 13.98 | 51.62 ± 19.41 | 0.56 | 0.62 ± 0.08 |
| Dentate gyrus | 222.54 ± 46.25 | 232.96 ± 48.57 | 0.97 | 0.80 ± 0.04 |

Reliability was assessed from segmentations made throughout the total hippocampal body. Example segmentations on three slices of the hippocampal body on one MRI scan are shown.

Fig 21. Example Segmentations on MRI by two raters from the feasibility assessment.

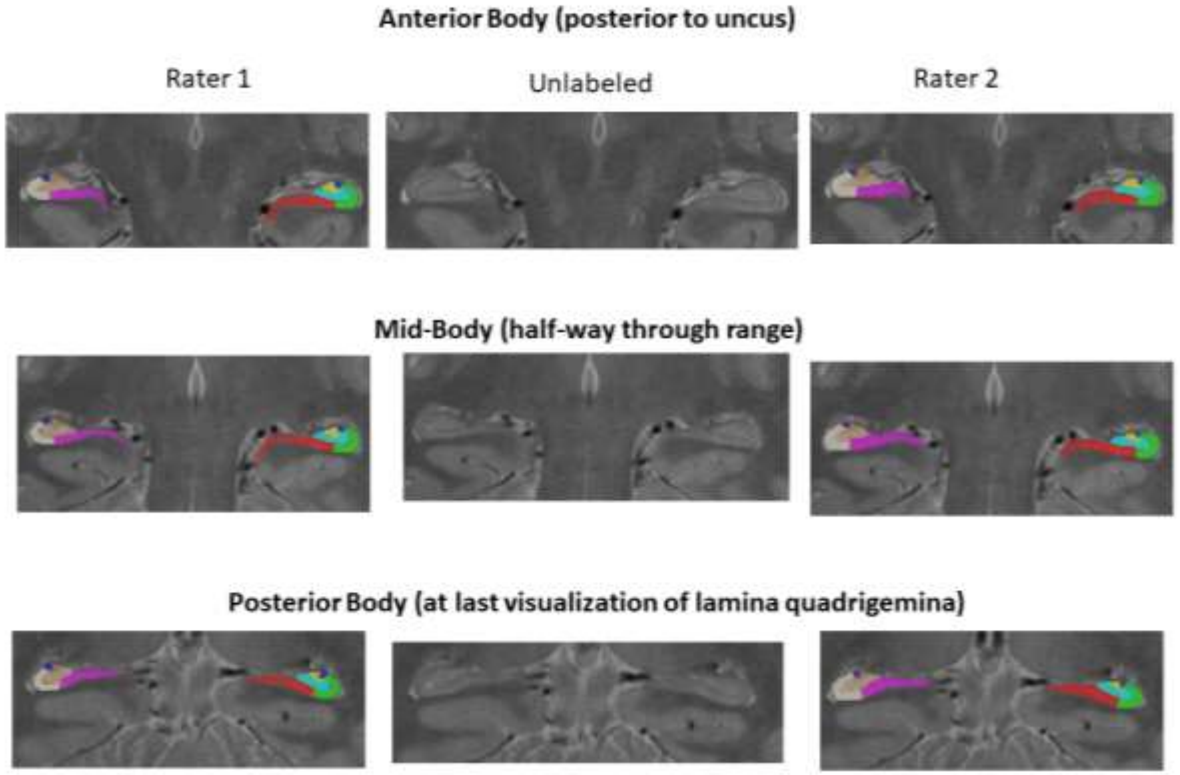

*Rater Experience*. When asked on a 7-point scale (1—Not at All, 7—Extremely Well), rater 2 indicated a poor understanding of the protocol (rating = 3) and rater 1 indicated understanding the protocol well (rating = 6); although both raters indicated that the rules were well defined in the training documents (both ratings = 6). When asked about the overall difficulty of performing the tracings on a 7-point scale (1—Very Easy, 7—Very Difficult), rater 2 indicated it was difficult (response = 6) and rater 1 indicated it was easy (response = 2).

When asked, both raters indicated that they had a similar experience applying the protocol on anterior and posterior slices. Both raters reported that it was more difficult to apply the protocol on a hippocampus with unusual shape, and one thought this introduced bias to their tracings, whereas the other did not. Both raters indicated that images with poor contrast that compromised visualization of the SRLM made applying the rules more difficult, and one rater believed this introduced bias when tracing whereas the other did not.

> One rater elaborated: “I felt that unusual hippocampal shapes were problematic for the CA3-DG boundary. My best recollection is that this came up when the hippocampus was flat (very stretched in the medial-lateral direction relative to its dorsal-ventral dimension). My instinct was that I wound up either misusing the "most superior, middle point of the hippocampus" (4) instructions in this case, or that a modified rule might need to be implemented for such scenarios (because the resulting CA3-DG boundary appeared unusual in some cases)…”

Rater 2 indicated that the endfolial rule for CA3-dentate gyrus boundaries was “difficult” and rater 1 indicated it was “somewhat easy”.

*Validation.* The proposed boundary for the endfolial rule appears to overlap with the CA3 region and excludes the dentate gyrus from the label.

Fig 22. Visual comparison of the DG boundary between three neuroanatomists (blue, green, and yellow lines) and the endfolial rule (red line). Two different brains (Boston and Philadelphia samples) are shown in the panel.

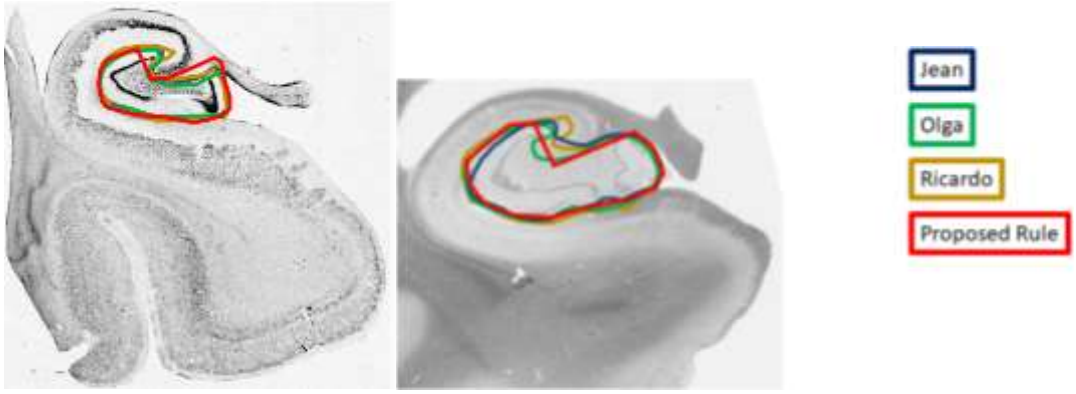

Comparison between Geometric and Endfolial Rules for CA3-Dentate Gyrus Boundary

In addition to considering the merits of each boundary rule for CA3-Dentate gyrus, we ask that you compare the two and vote for which protocol to adopt and move forward for testing.

A summary of the comparisons for reliability and rater experience, as well as comparisons to evaluate construct validity are provided below.

*Protocol Reliability.* Reliability estimates of the dentate gyrus measurement were comparable between protocols; but reliability was notably worse for CA3 with use of the endfolial rule.

| **Subfield** | **Rater 1, Bilateral Average Volume Across Cases (mm$^{3}$’ M ± SD)** | **Rater 2, Bilateral Average Volume Across Cases (mm$^{3}$; M ± SD)** | **ICC(2)** | **Average Dice (M ± SD)** |
|---|---|---|---|---|
| Geometric: CA3 | 50.85 ± 13.41 | 42.56 ± 11.03 | 0.84 | 0.69 ± 0.07 |
| Endfolial: CA3 | 53.36 ± 13.98 | 51.62 ± 19.41 | 0.56 | 0.62 ± 0.08 |
| Geometric: Dentate gyrus | 236.73 ± 50.87 | 242.03 ± 54.52 | 0.99 | 0.79 ± 0.06 |
| Endfolial: Dentate gyrus | 222.54 ± 46.25 | 232.96 ± 48.57 | 0.97 | 0.80 ± 0.04 |

*Rater Experience.* When asked to choose, both raters indicated that the geometric rule was easier to understand and easier to use when tracing; they felt more confident in their use of the geometric rule; and both preferred the geometric rule.

| | Geometric | | | | Endfolial | | |
|---|---|---|---|---|---|---|---|
| Rater | Understanding the Protocol | Overall Difficulty | CA3-Dentate Gyrus Difficulty | | Understanding the Protocol | Overall Difficulty | CA3-Dentate Gyrus |
| Rater 1 | 6 (Very Well) | 3 (Somewhat Easy) | 3 (Somewhat Easy) | | 6 (Very Well) | 2 (Easy) | 3 Somewhat Easy |
| Rater 2 | 7 (Extremely Well) | 6 (Difficult) | 1 (Very Easy) | | 3 (Poor) | 6 (Difficult) | 6 (Difficult) |

*Validation*. The endfolial rule appears to enjoy better face validity and is a closer approximation of the dentate gyrus, excluding CA3.

Fig 23. Visual comparison of the dentate gyrus boundary between three neuroanatomists and the geometric and enfolial rules. The neuroanatomists' notations were segmented and colored (blue—JA; green—OK; yellow—RI) to aid visual comparison. Only the DG labels are shown. The bold red lines indicates the placement of the boundaries according to the draft protocols. Two different brains are depicted in the panel, left and right. **Top Panel: Geometric; Bottom Panel: Endfolial.**

Geometric:

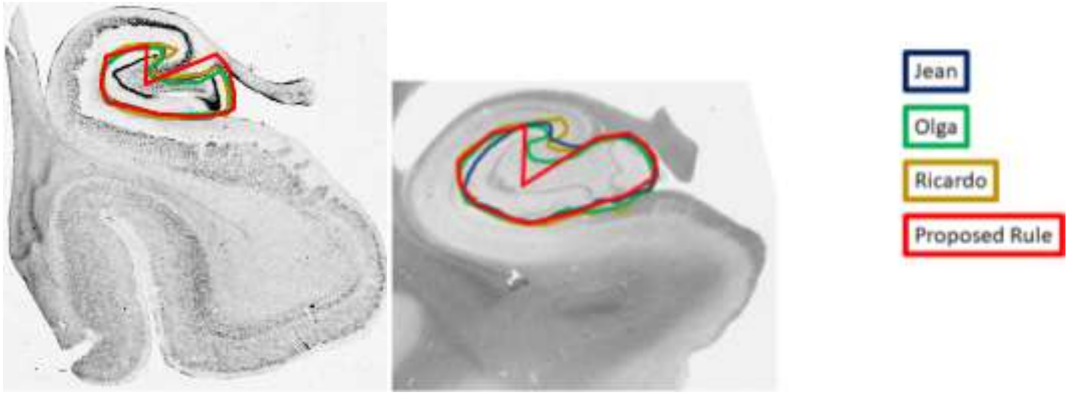

Endfolial:

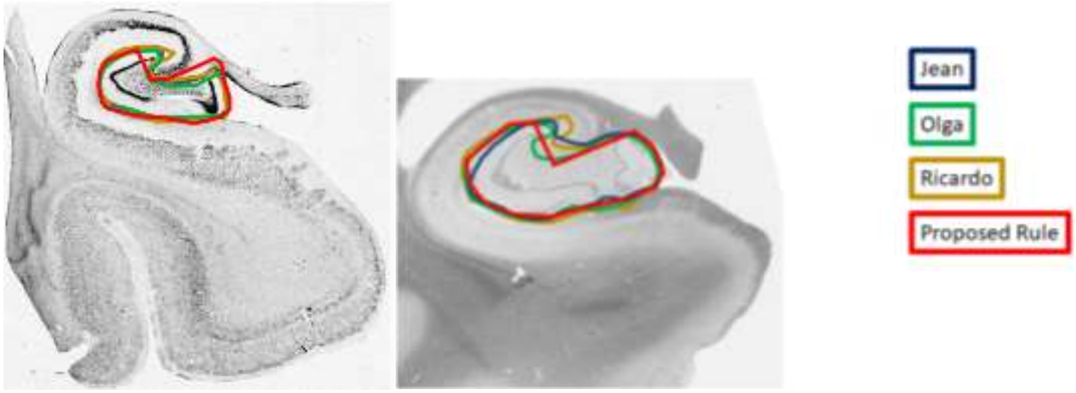

Evaluation of SRLM Boundary

The SRLM lines the internal edge of the hippocampal fissure and separates the dentate gyrus from the CA fields. The proposed protocol includes the SRLM within the subiculum and CA subfield labels, and excludes it from the dentate gyrus.

This protocol decision was informed by review of published ex vivo MRI and histology data (Adler et al., 2018; de Flores et al., 2019).

A sample of 9 subjects were collected with high-resolution (~200 × 200 × 200 $\mu m^3$) *ex vivo* MRI scans and detailed histology images. The histology images were manually segmented to delineate CA-SRLM (turquoise) and DG-SM (light blue) based upon cytoarchitecture. The histology-based segmentations were co-registered to the ex vivo MRI, and the result identified that  the dark band on T2-weighted MRI falls within the CA-SRLM region. Therefore, the proposed protocol excludes SRLM from the dentate gyrus.

Fig 24. Depiction of hypointense band on T2-weighted image that is interpreted as SRLM falls within the CA-SRLM histological region (Fig 6 from de Flores et al., 2019). The dark band (red arrows in a) thickness was measured on MRI following the dashed line starting at the medial anchor point (red circle) to the most lateral point of the hippocampus (green circle) (b; dark band measurement is depicted in red). The corresponding histology section (c) and its segmentation (d) were registered to the MRI, where measurements along the same dashed line were performed. (f) shows magnification of the slice showed in € 7 regions were labeled: CA1 (red), CA2 (green), CA3 (yellow), DG-SM (light blue), DG-G/H (dark blue), SRLM (cyan) and SUB (pink).

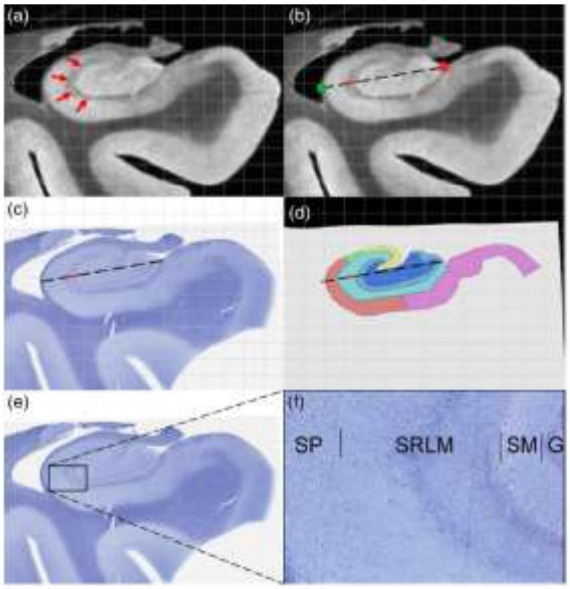

Complete Comparison of Proposed Protocol Boundaries to Neuroanatomists' Annotations on Histological Reference Set

#### Subiculum-Cornu Ammonis 1

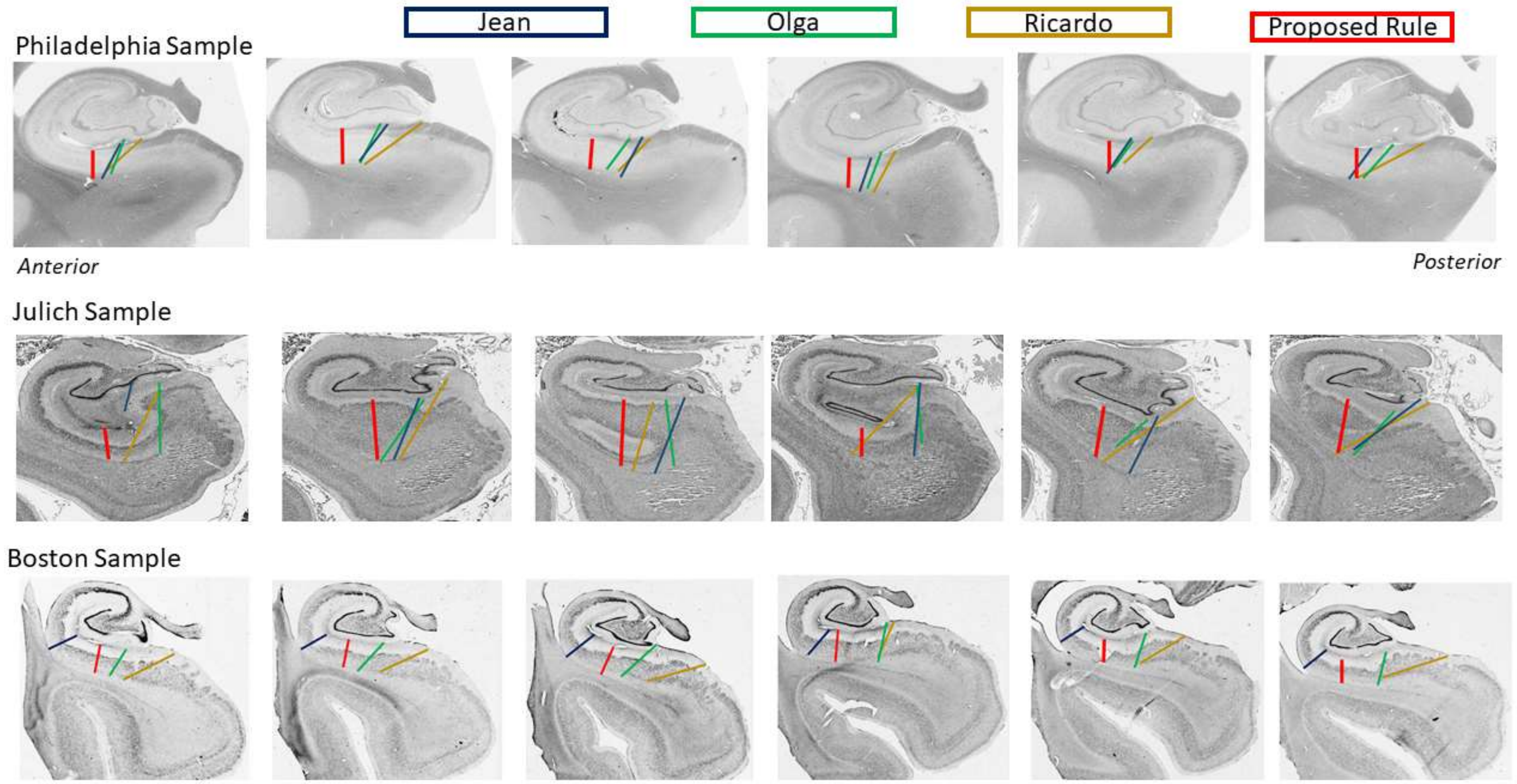

#### Cornu Ammonis 1-2

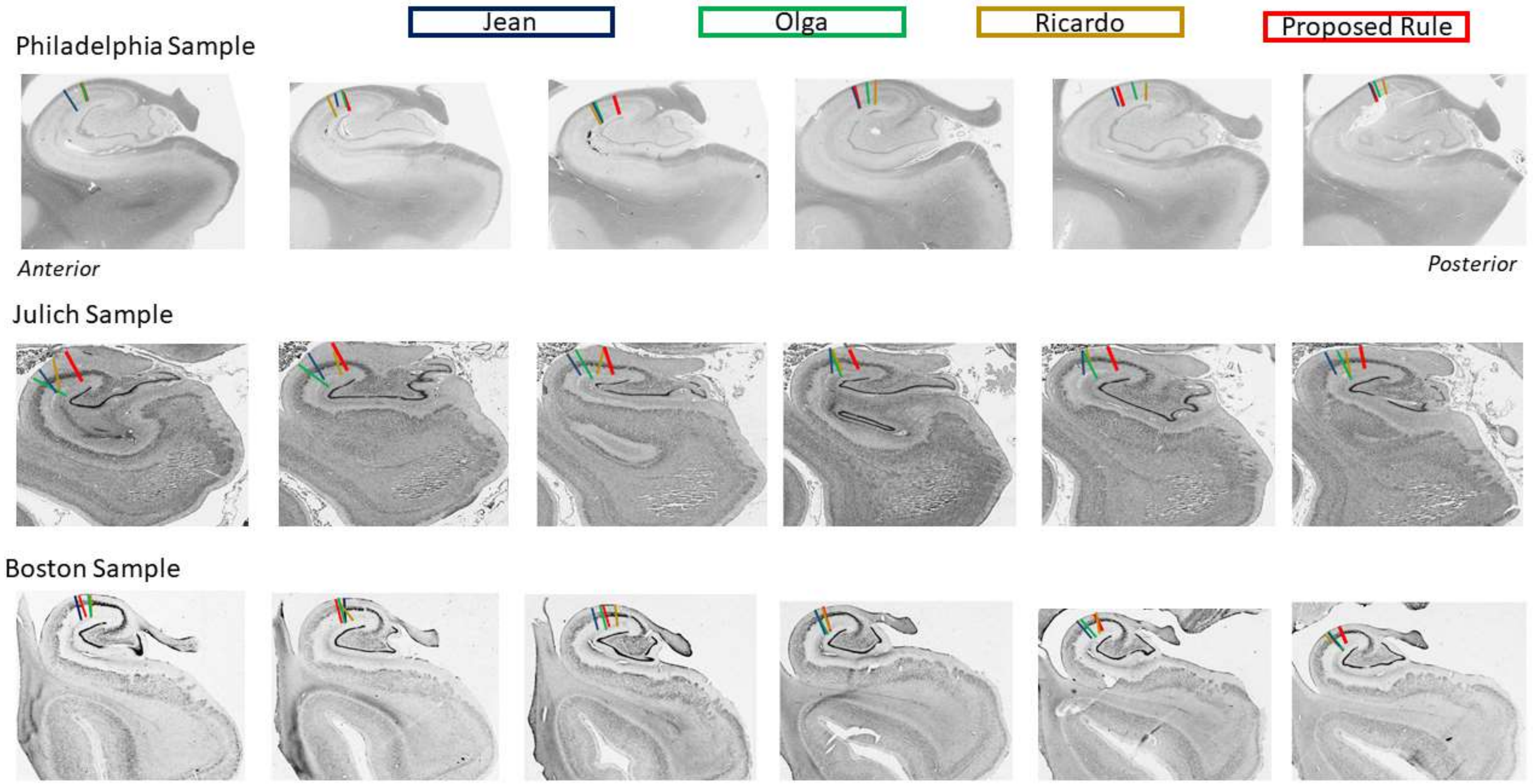

#### Cornu Ammonis 2-3

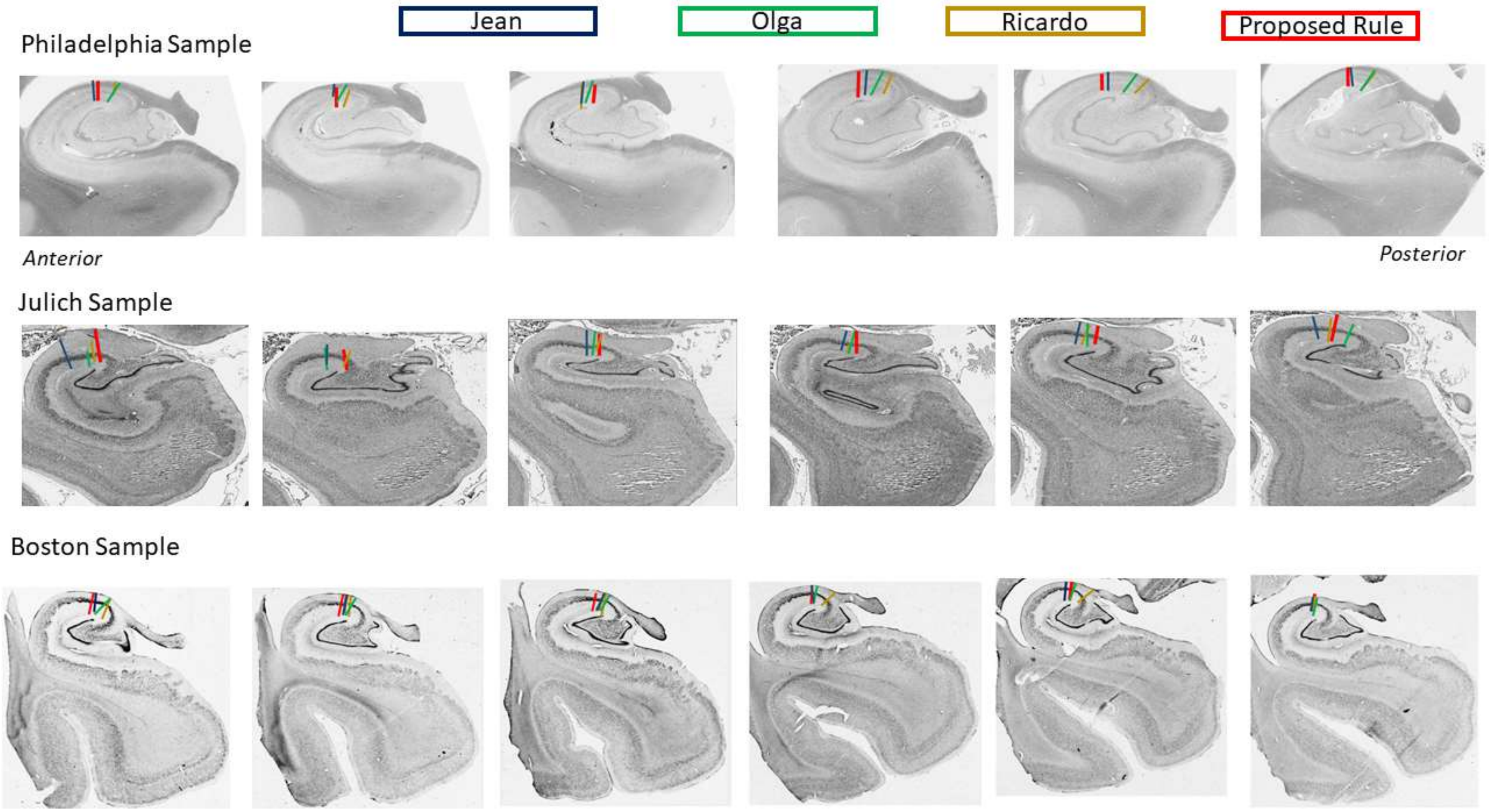

#### Cornu Ammonis 3-Dentate Gyrus (Geometric Rule)

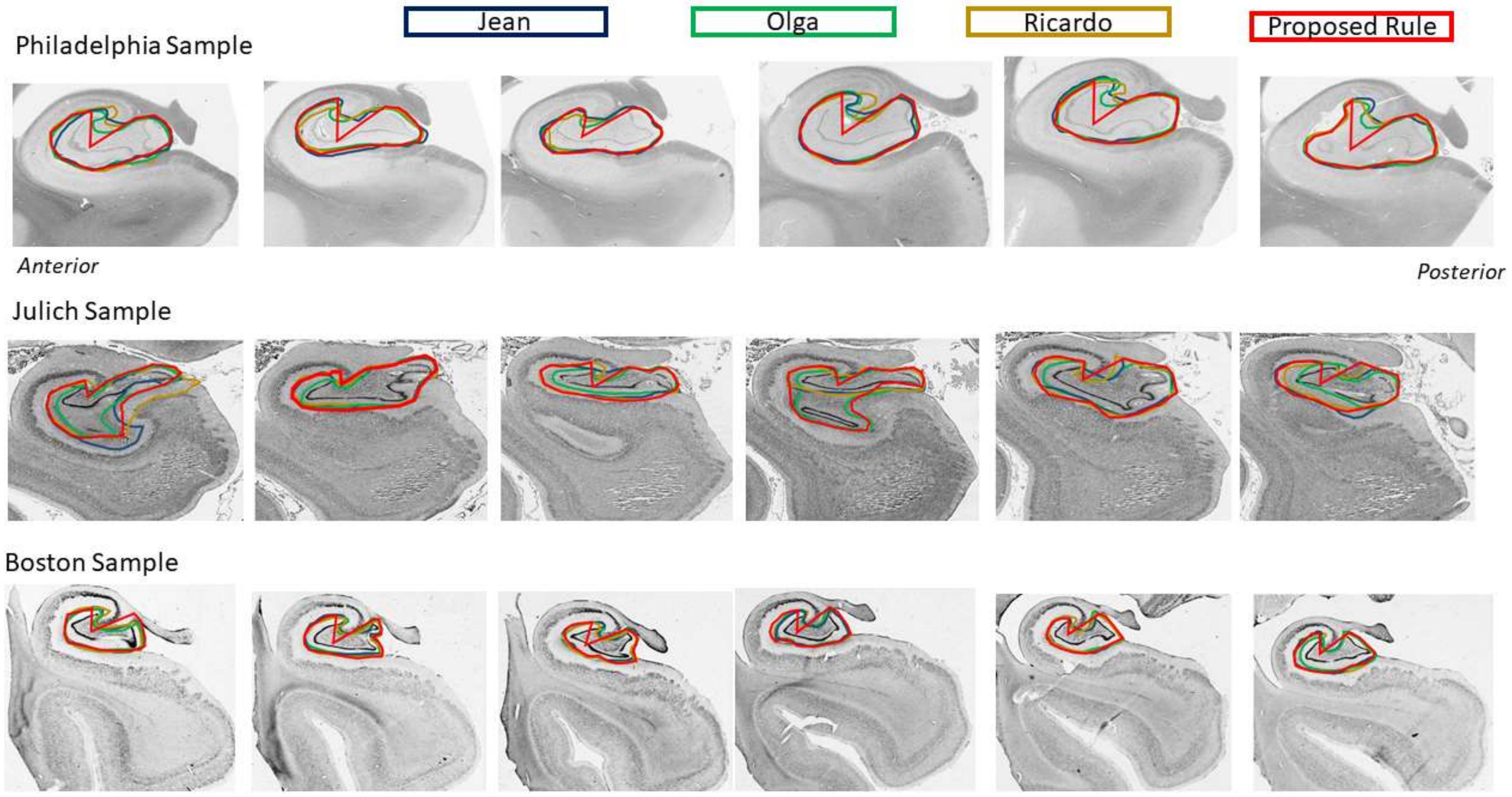

#### Cornu Ammonis 3-Dentate Gyrus (Protocol 2, Endfolial Rule)

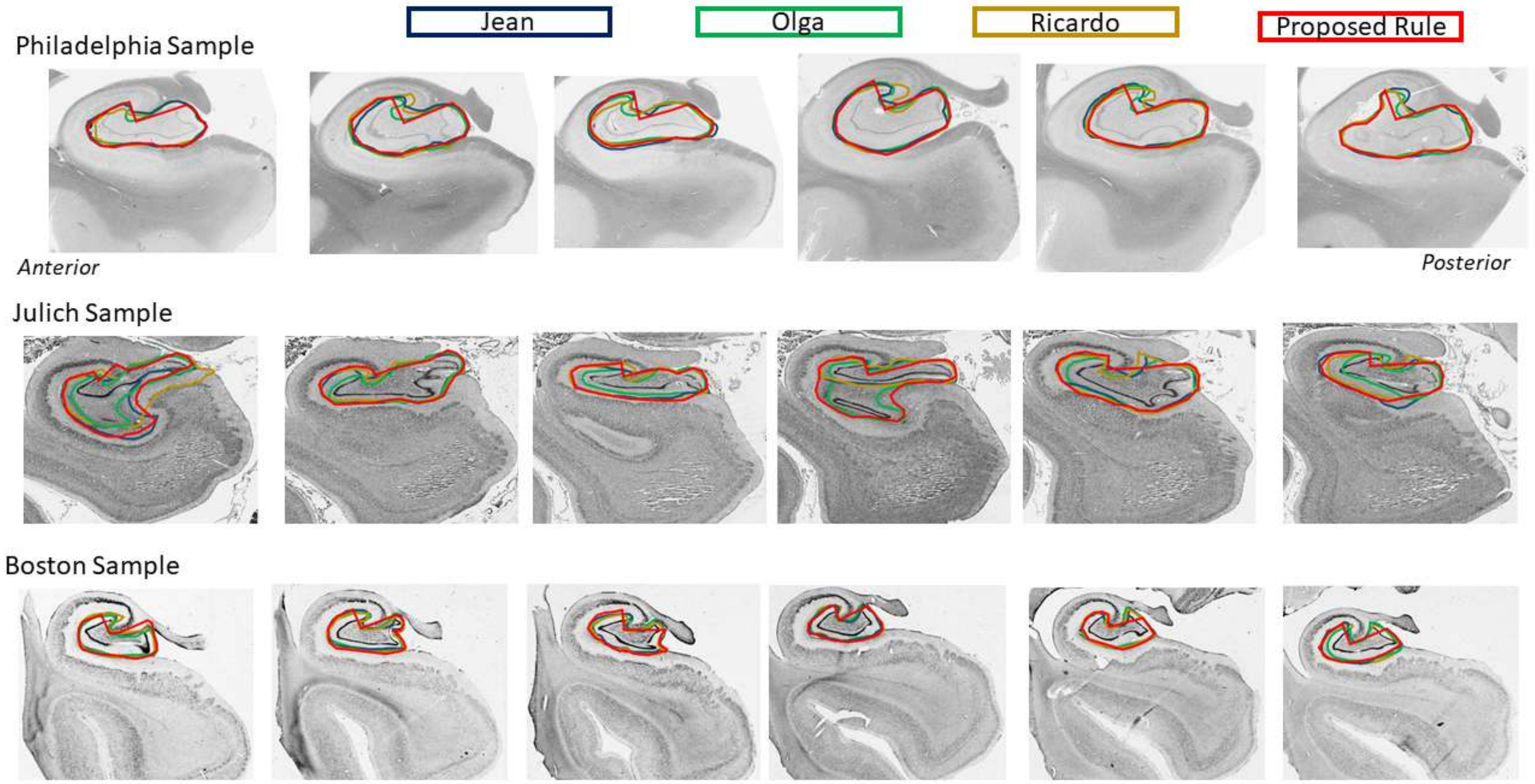

Complete Histological Reference Set with Anatomical Annotations and Inferred Segmentation

Three different post-mortem cases were collected and imaged with standard histological procedures by the neuroanatomists. These samples are labeled by the site of their collection: Boston, Julich and Philadelphia. Histological slices were selected to approximate the 2-mm slice thickness that is common of MRI protocols. The neuroanatomists (JA, OK, RI) each labeled every slice of the hippocampal body, approximating a 2-mm slice thickness, within the length of the hippocampal body.

The segmentations for each slice are presented below with a comparison across neuroanatomists. The labels were interpreted for volume masks and labeled as Subiculum (blue), CA1 (grey), CA2 (green), CA3 (orange), and dentate gyrus (yellow).

#### Boston Sample

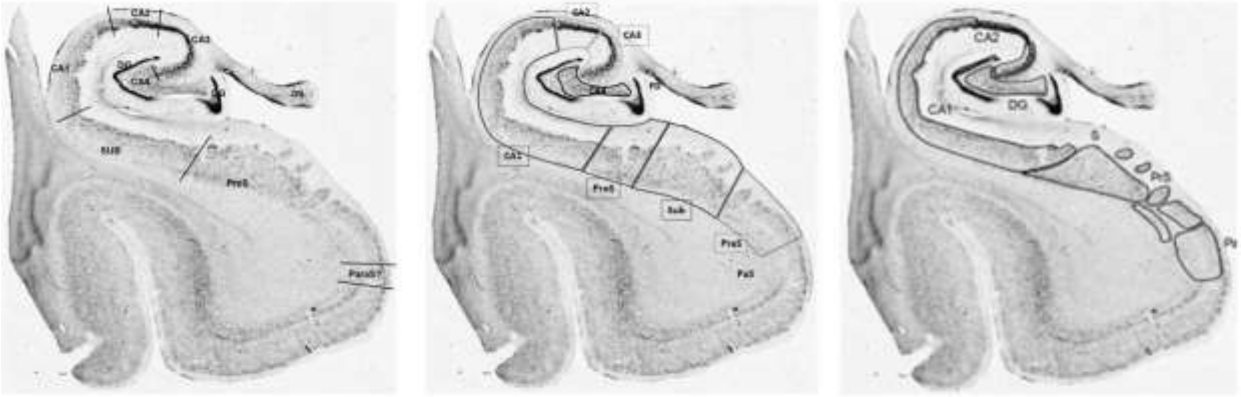

Jean Olga Ricardo

B-531

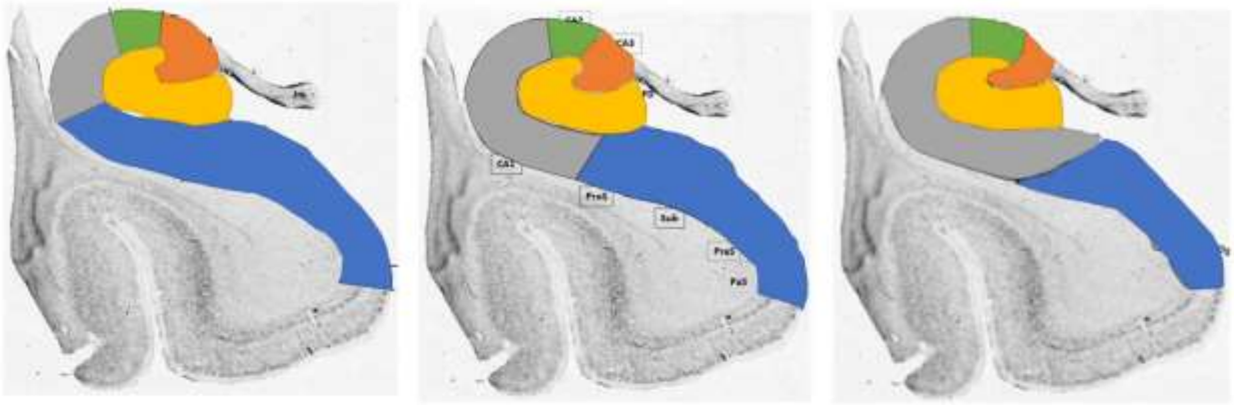

Jean Olga Ricardo

B-531

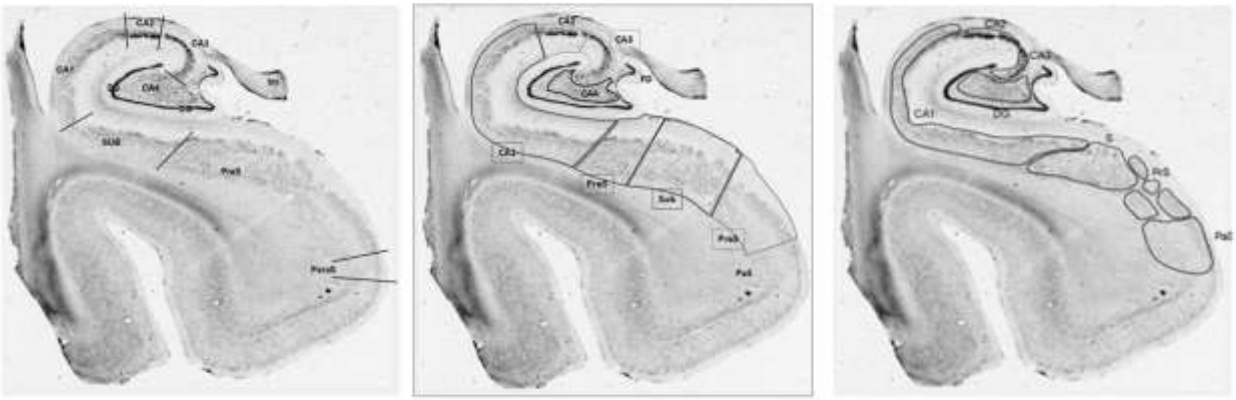

Jean Olga Ricardo

B-555

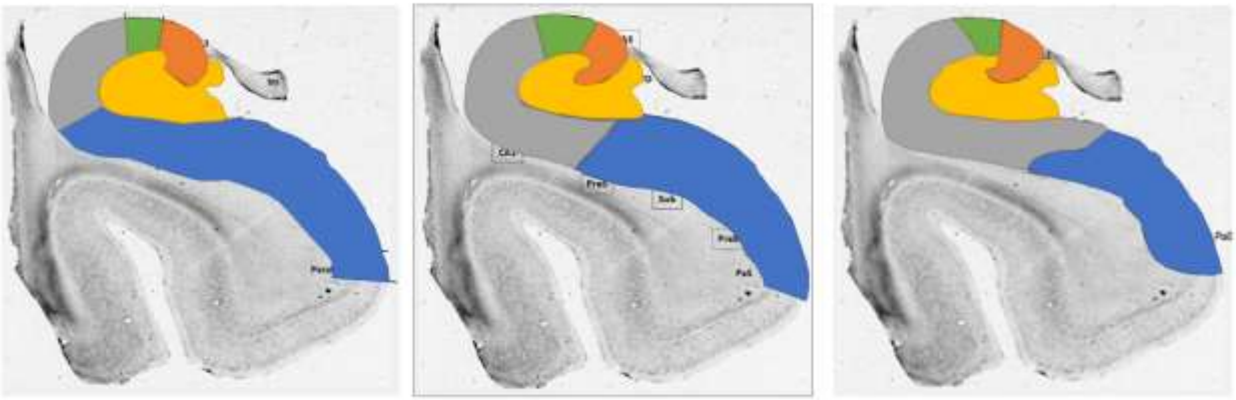

Jean Olga Ricardo

B-555

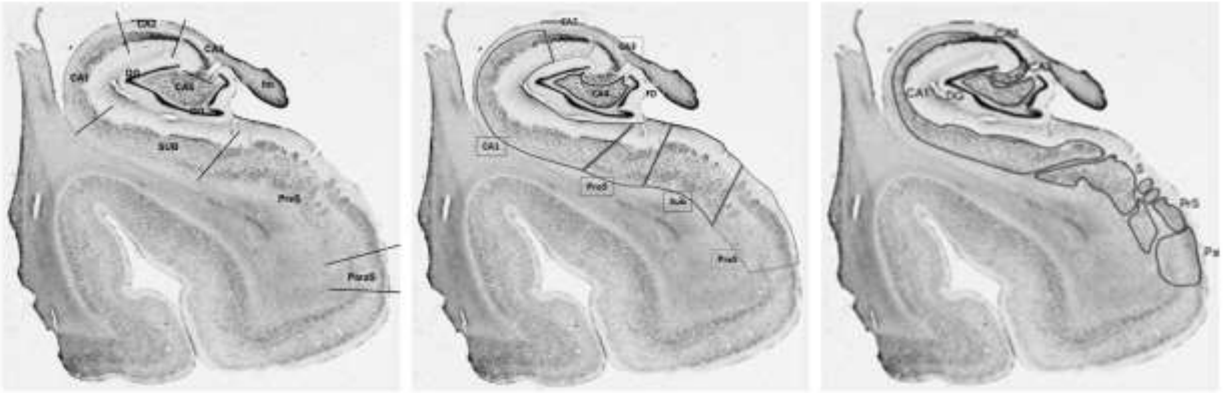

Jean Olga Ricardo

B-576

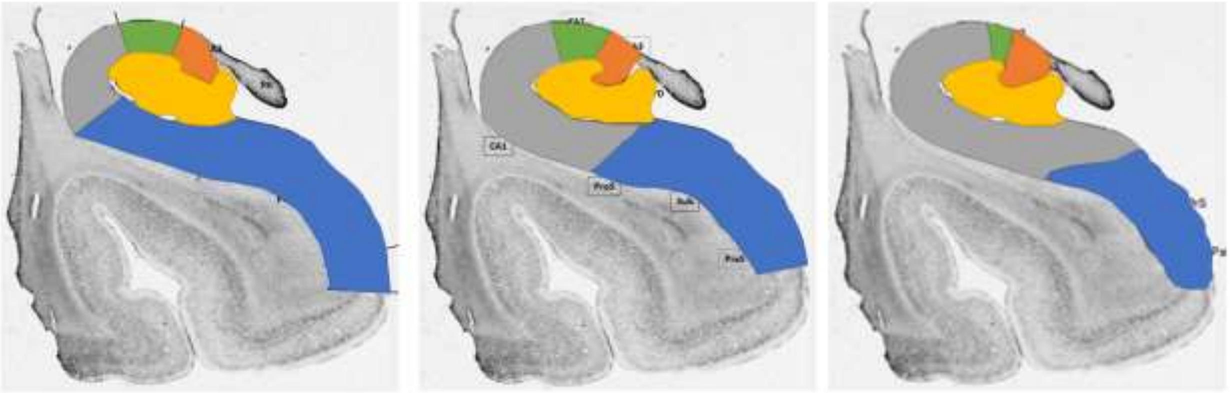

Jean Olga Ricardo

B-576

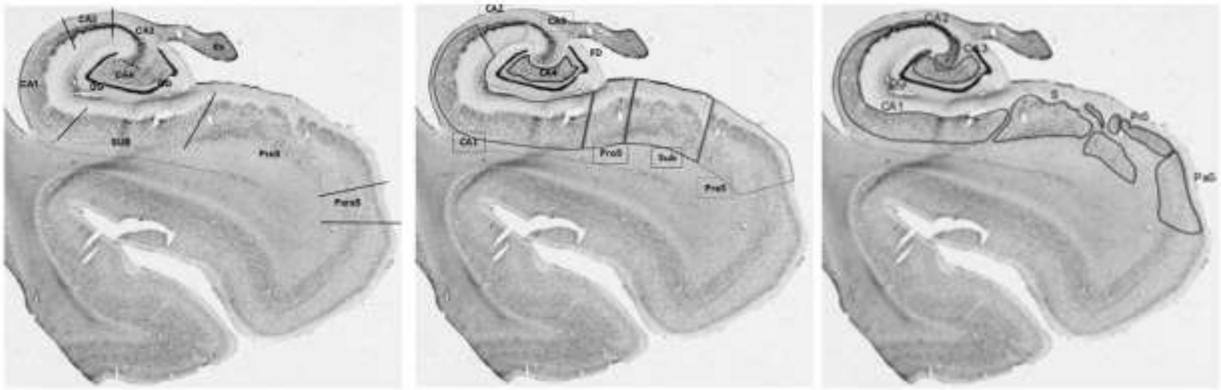

Jean Olga Ricardo

B-596

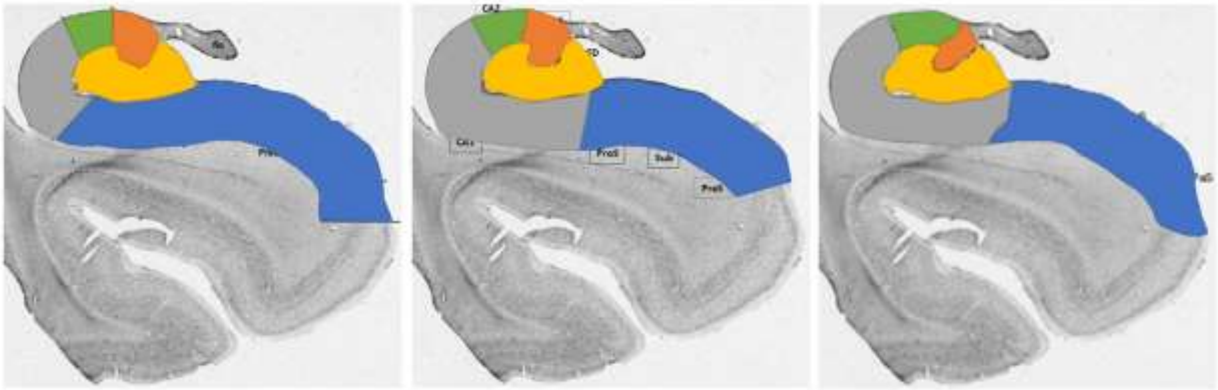

Jean Olga Ricardo

B-596

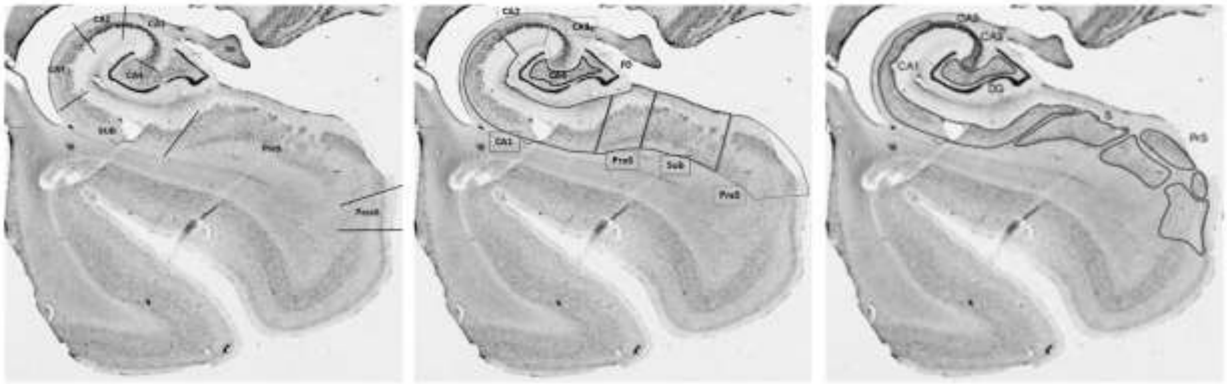

Jean Olga Ricardo

B-616

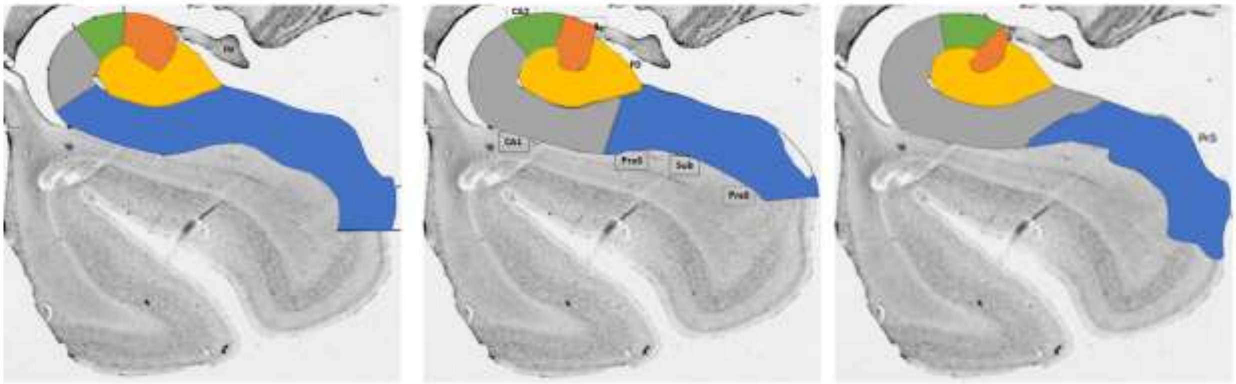

Jean Olga Ricardo

B-616

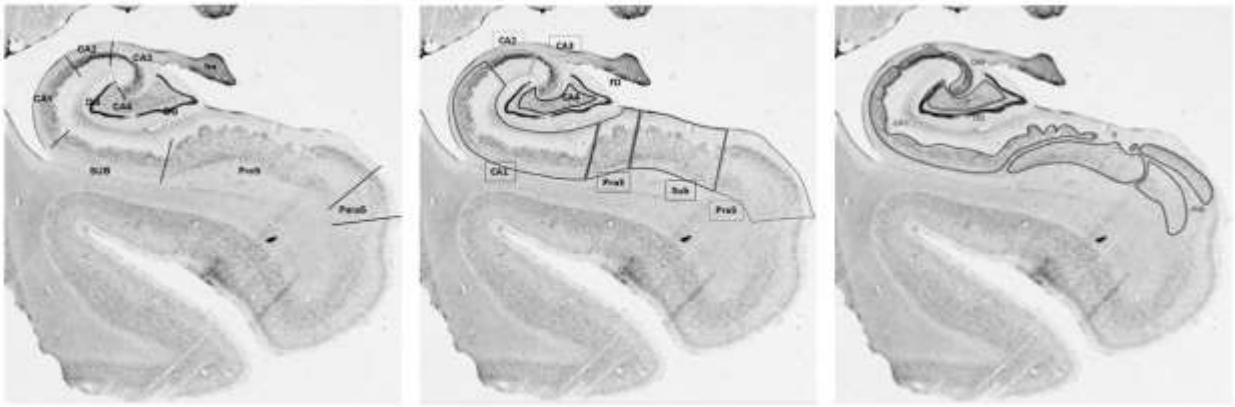

Jean Olga Ricardo

B-621

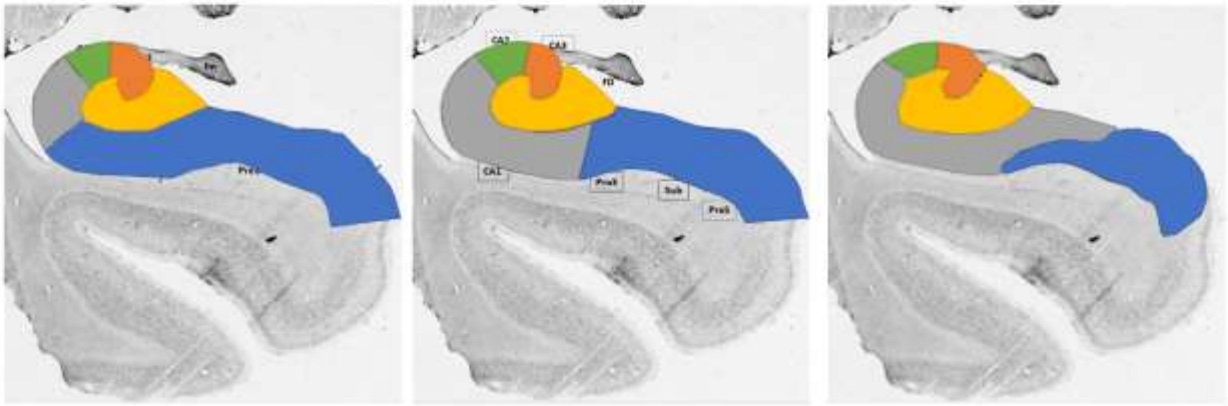

Jean Olga Ricardo

B-621

#### Julich Sample

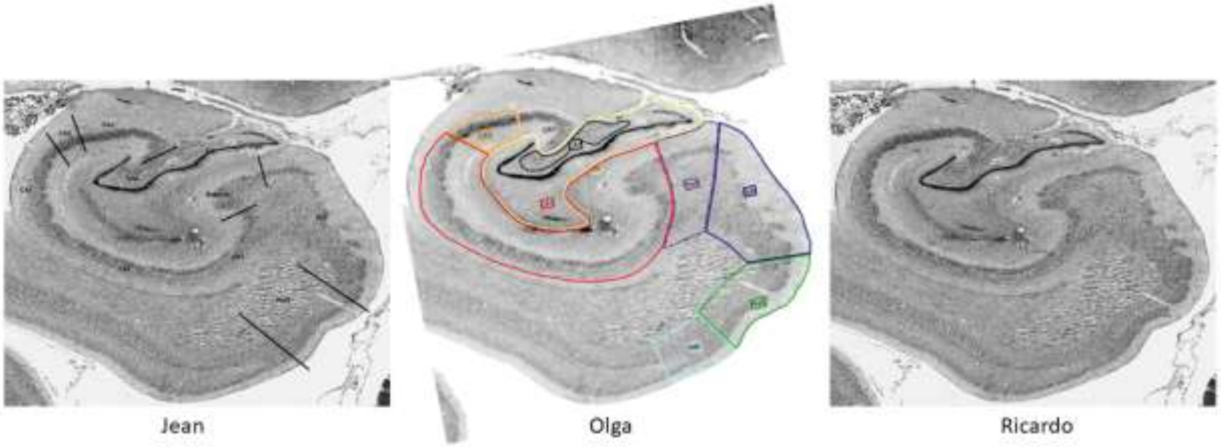

J-306

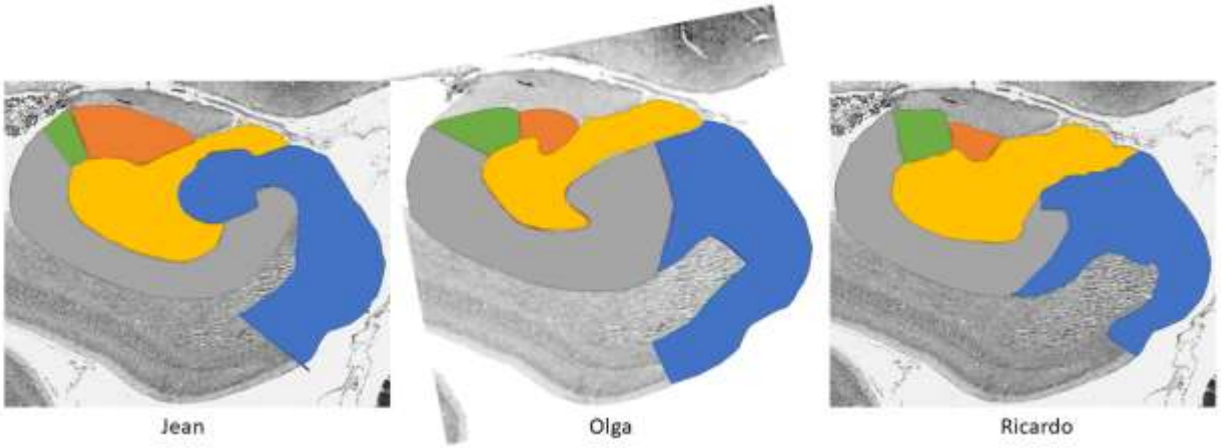

J-306

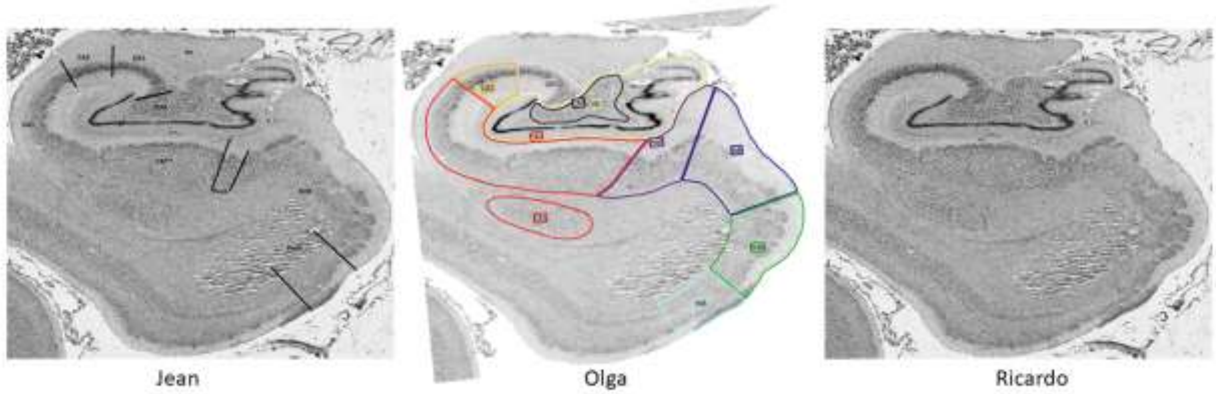

J-312

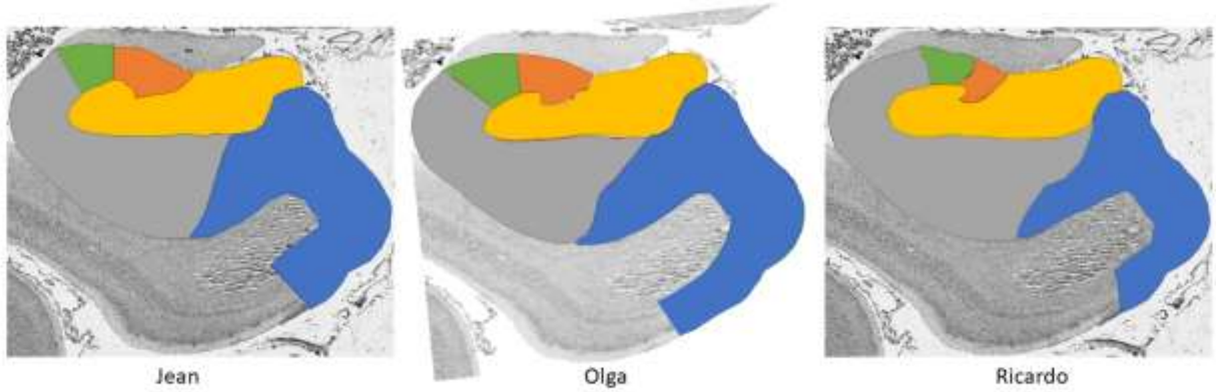

J-312

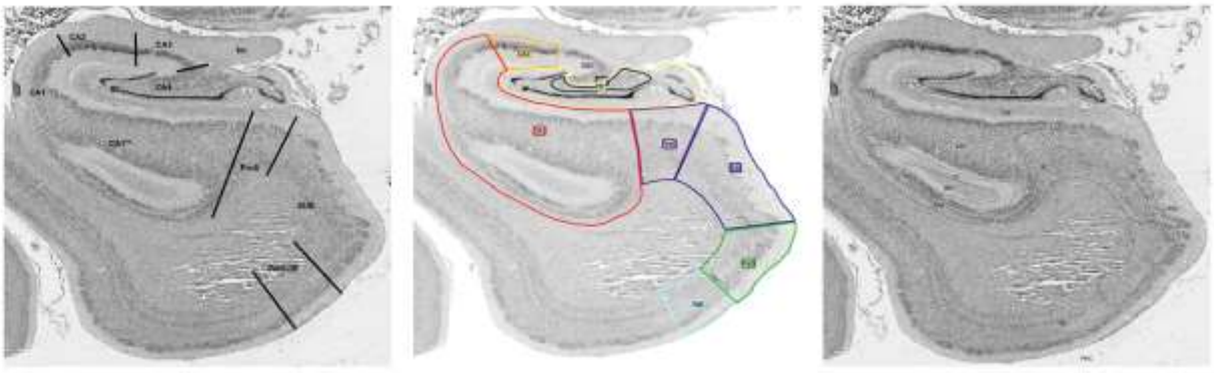

Jean Olga Ricardo

J-318

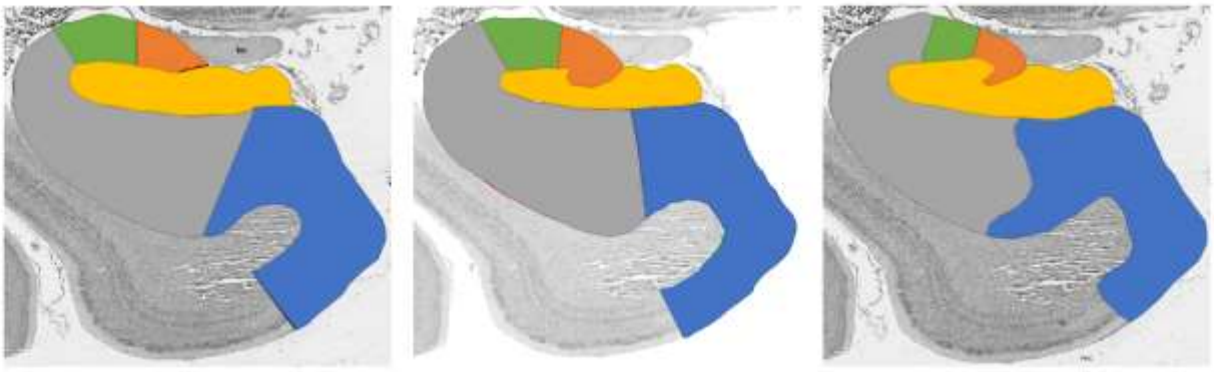

Jean Olga Ricardo

J-318

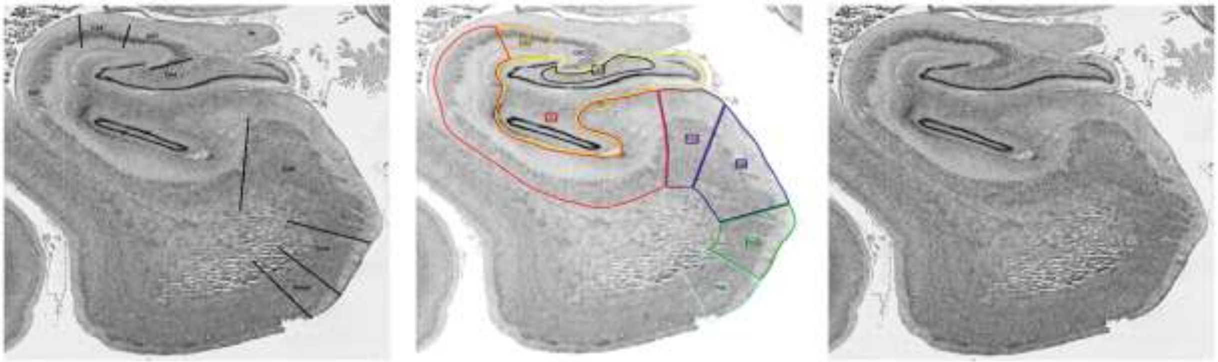

Jean Olga Ricardo

J-324

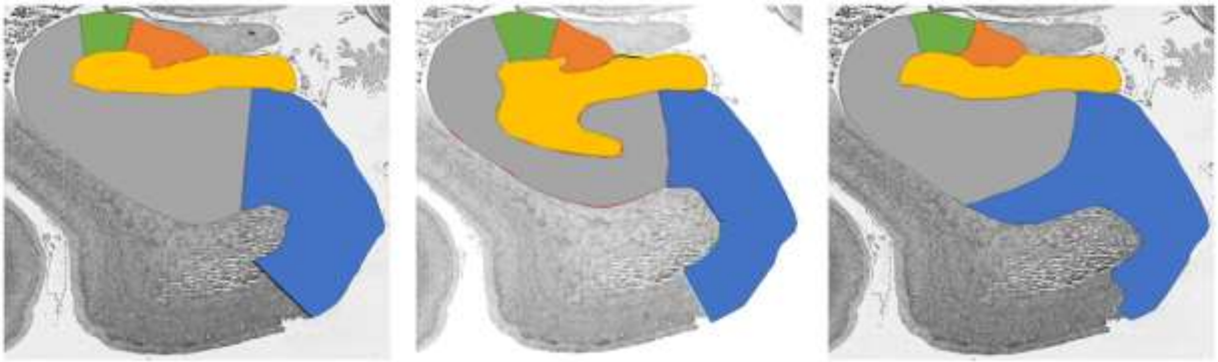

Jean Olga Ricardo

J-324

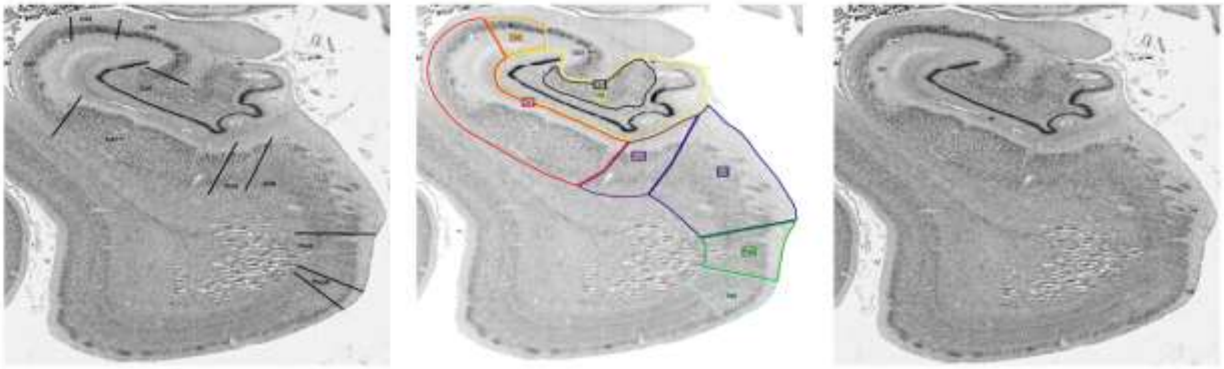

Jean Olga Ricardo

J-330

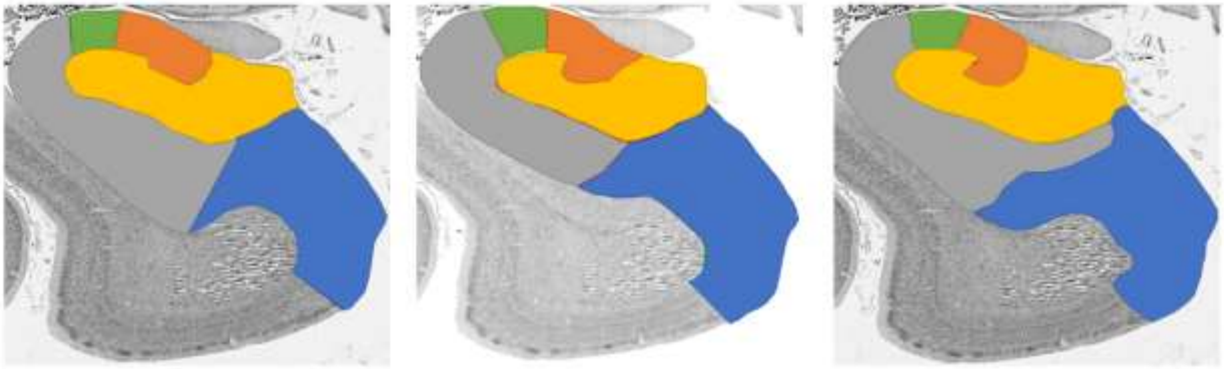

Jean Olga Ricardo

J-330

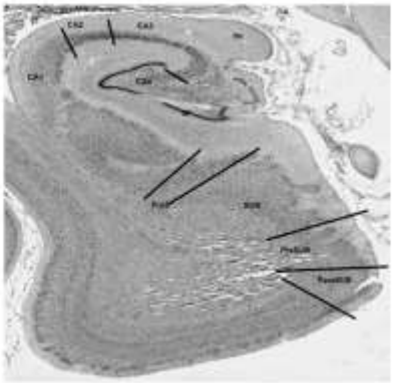

Jean

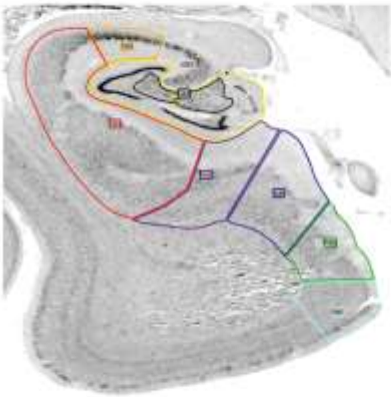

Olga

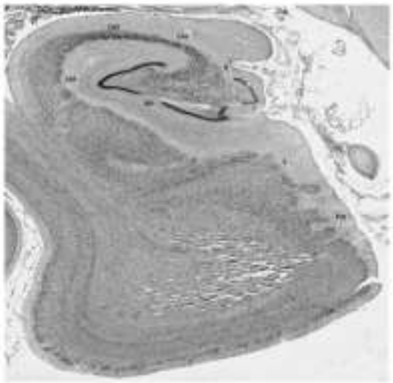

Ricardo

J-336

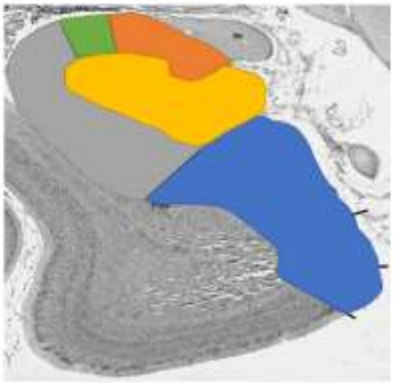

Jean

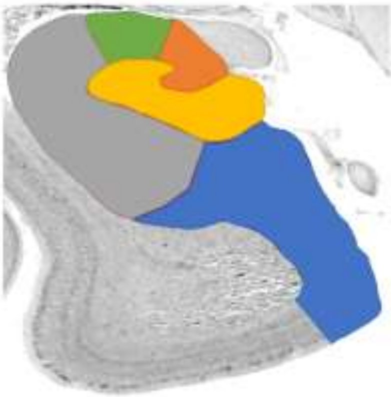

Olga

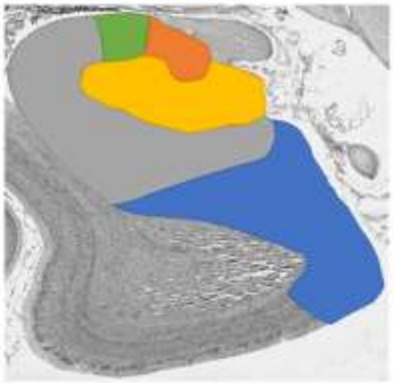

Ricardo

J-336

#### Philadelphia Sample

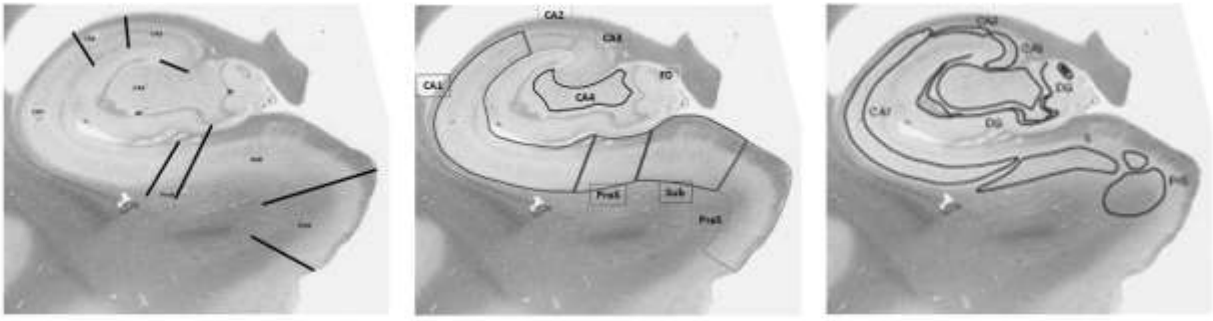

Jean Olga Ricardo

P-513

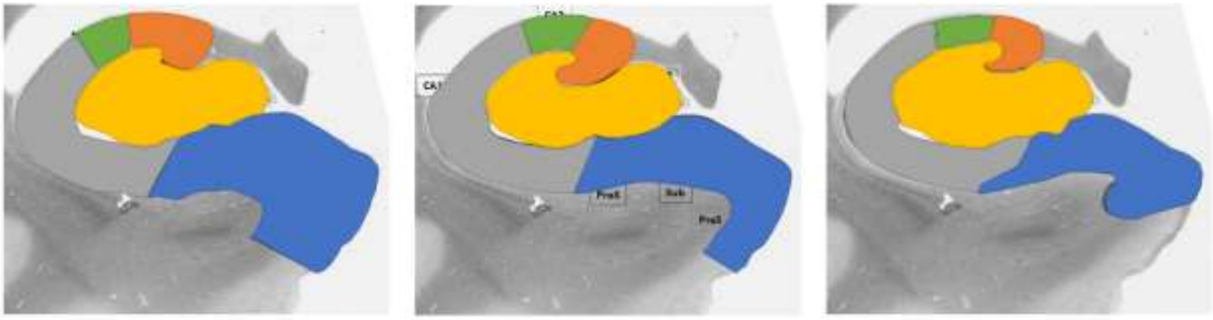

Jean Olga Ricardo

P-513

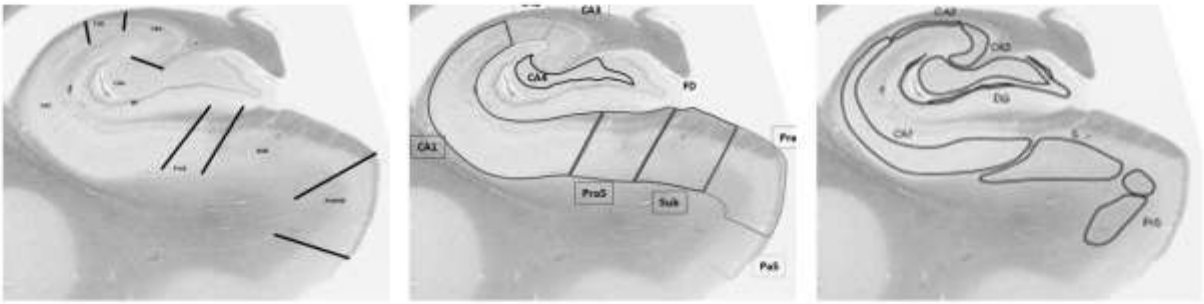

Jean Olga Ricardo

P-526

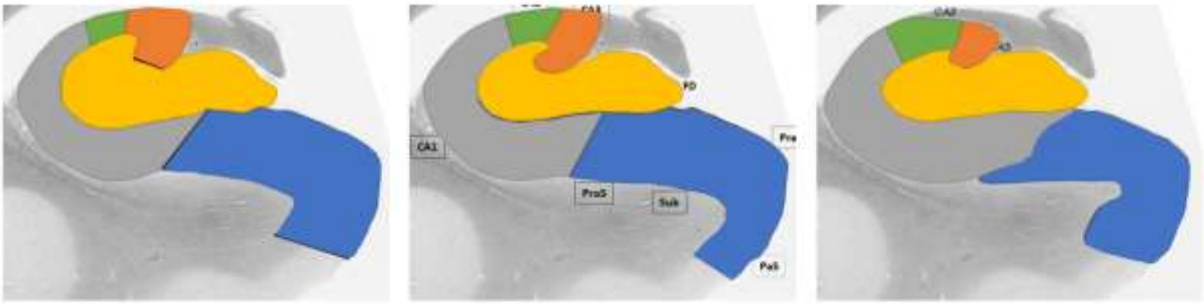

Jean Olga Ricardo

P-526

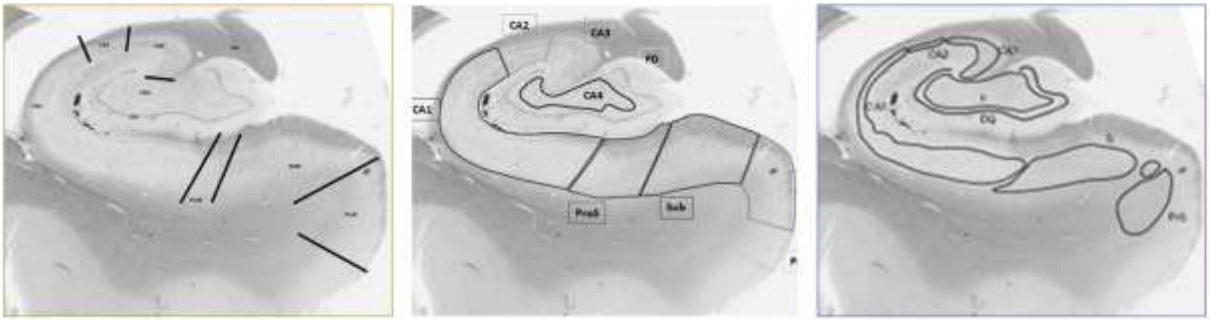

Jean Olga Ricardo

P-543

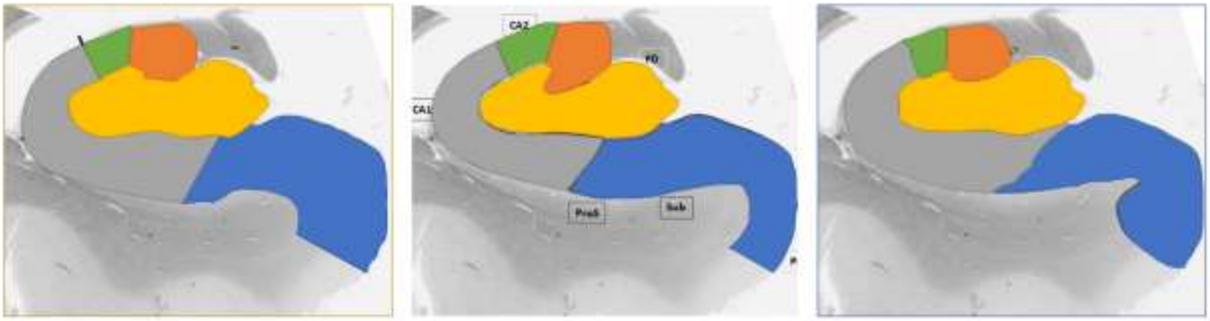

Jean Olga Ricardo

P-543

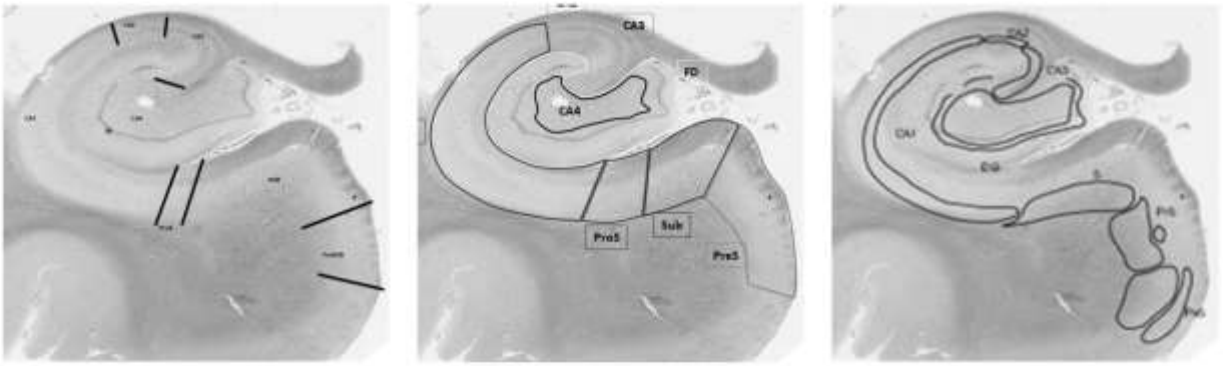

Jean Olga Ricardo

P-626

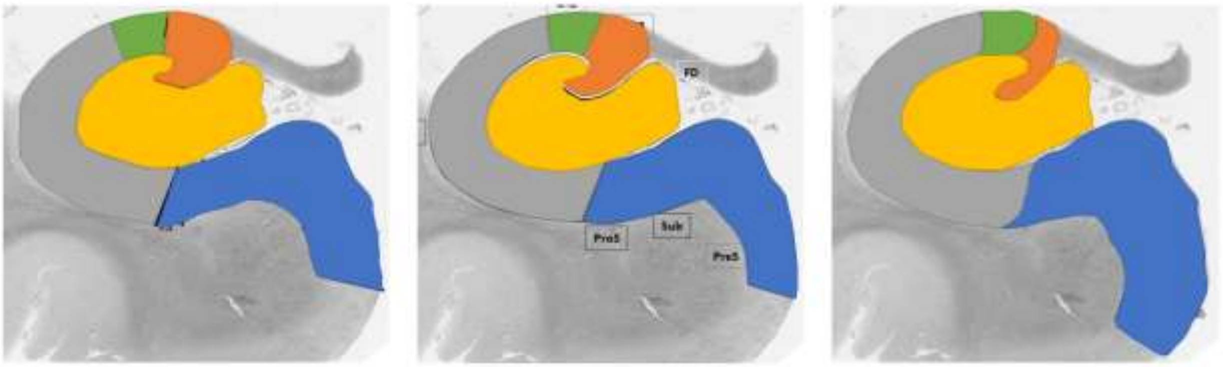

Jean Olga Ricardo

P-626

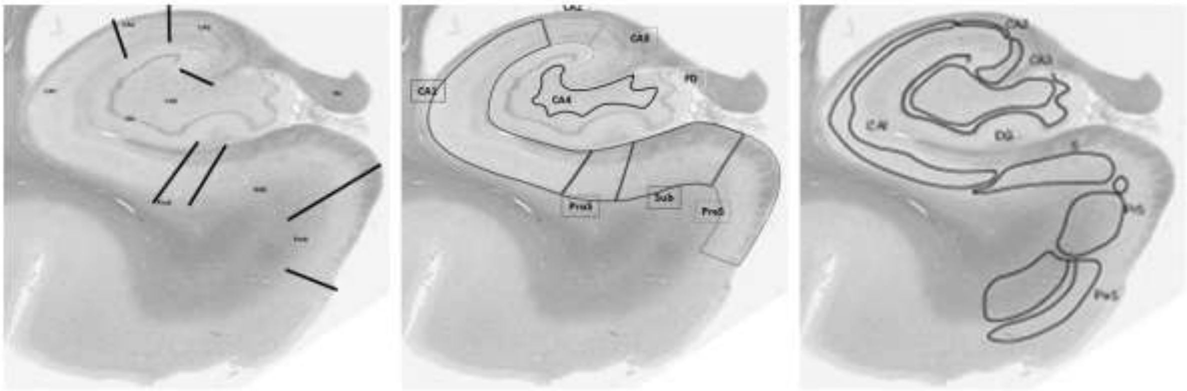

Jean Olga Ricardo

P-635

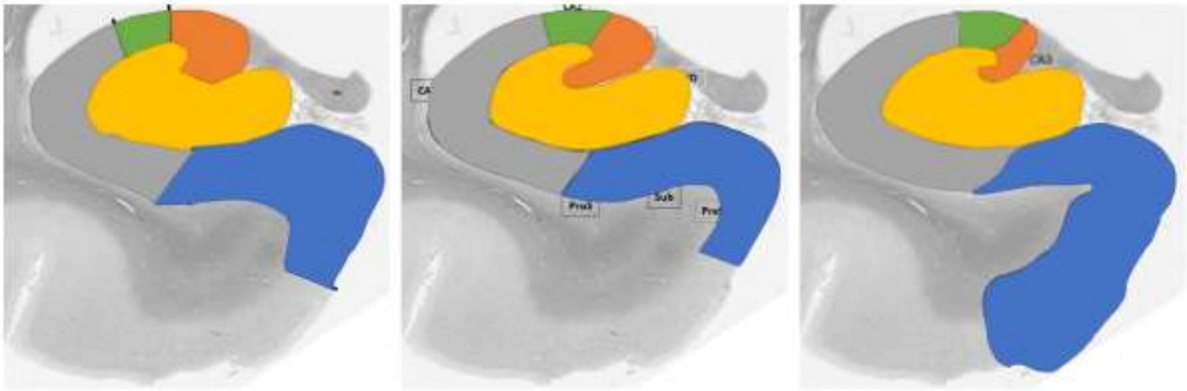

Jean Olga Ricardo

P-635

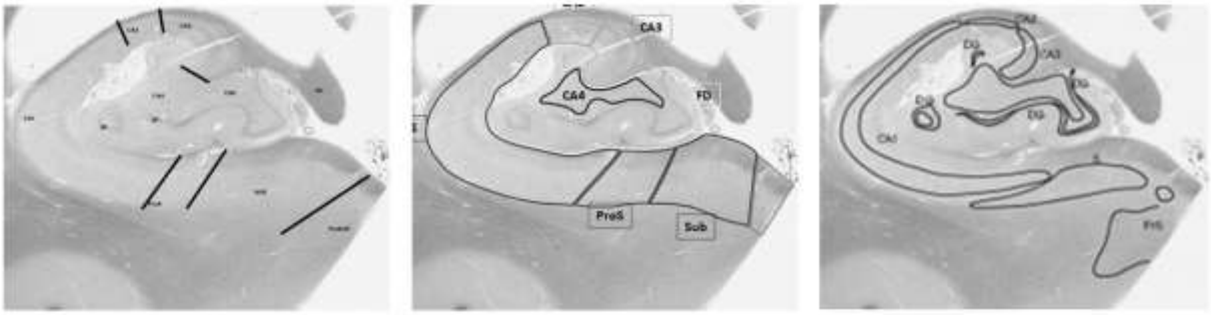

Jean Olga Ricardo

P-671

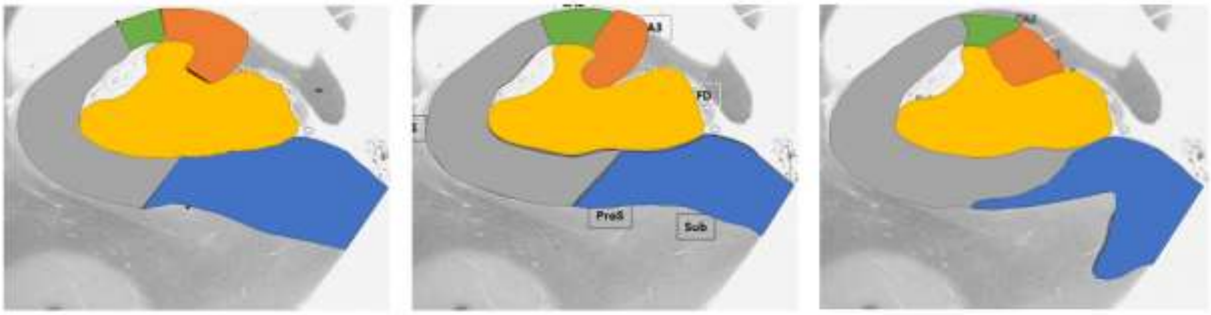

Jean Olga Ricardo

P-671

Appendix A: Protocol Training Documentation

#### DRAFT Inner Boundary Definitions in the Hippocampal Body

There are two proposed rule sets: (1) the Geometric Heuristic Rule and (2) a modification with the Endfolial Pathway Rule.

##### Geometric Heuristic

**<u>Protocol Description:</u>** Boundaries within the hippocampal body are determined by a geometric heuristic that approximates the location of anatomical landmarks observable in histological samples. (**1**) Line 1 should be anchored at the opening of the hippocampal fissure (**a,** the "arm pit") and extended to the most lateral, outside edge of the alveus (white matter structure) of the CA1 sector. (**2**) Line 2 is then placed perpendicular to the middle of Line 1, extending from the most superior edge of the hippocampus to the parahippocampal white matter. (**3**) An additional vector is extend from the point of bisection at 30° to the lateral side and (**4**) another vector at 45° to the medial side.

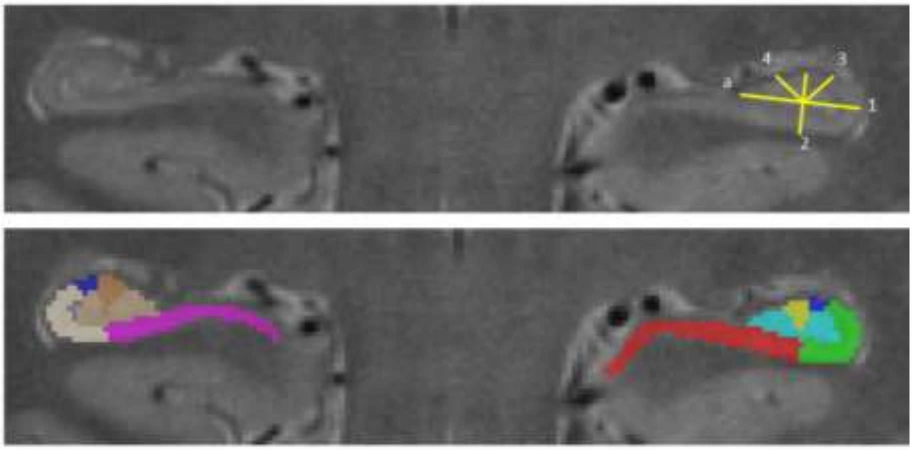

| | | |
|---|---|---|
| 1. Line 1 should be anchored at the opening of the hippocampal fissure adjacent to the superior edge of the subiculum (**a,** the "arm pit") and extended to the most lateral, outside edge of the alveus (white matter structure) of the CA1 sector. | 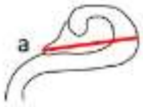 | 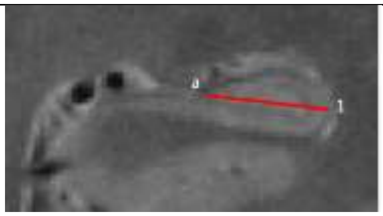 |

| | | |
|---|---|---|
| 2. Line 2 is then placed perpendicular to the middle of Line 1, extending from the most superior edge of the hippocampus to the parahippocampal white matter. | 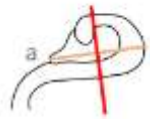  | 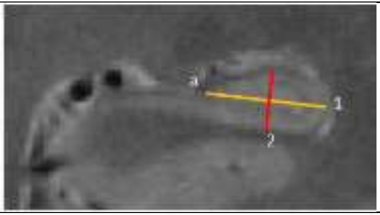  |
| 3. An additional vector is extend from the point of bisection at 30° to the lateral side. | 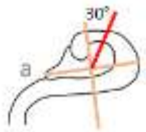  | 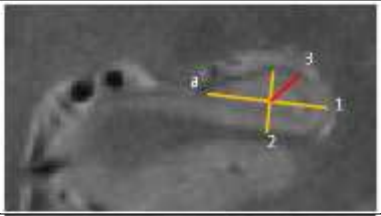  |
| 4. Extend another vector at 45° to the medial side. | 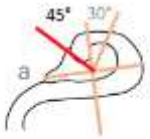  | 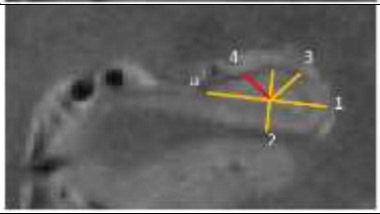  |

Cysts or hippocampal sulcal cavities should not be labeled and are excluded from the subfield labels.

**<u>Manual Tacing Example in ITKSnap:</u>**

The following is an example of the steps to implement the protocol in ITKSnap. We will use the annotation tool to map out the geometric heuristic and then manually segment the subfields.

1. Load the Image and Segmentation Labels
2. Click the magnifying glass, and **zoom 4x**
3. Click the **C** button to only view the coronal plane
4. With the Annotation Tool 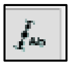
    a. Draw Line 1 from the **a** anchor point—opening of the hippocampal fissure, adjacent to the superior edge of the subiculum—to the most lateral edge
        i. If the lateral edge is flat, choose the middle most lateral point
    b. Draw Line 2 at the mid-point of Line 1, perpendicular to it, extending from the white matter to the superior edge
    c. Draw a bisector at 30° from Line 2 at the bisection to the lateral, superior edge

d. Draw a bisector at 45° from Line 2 at the bisection to the medial, superior edge

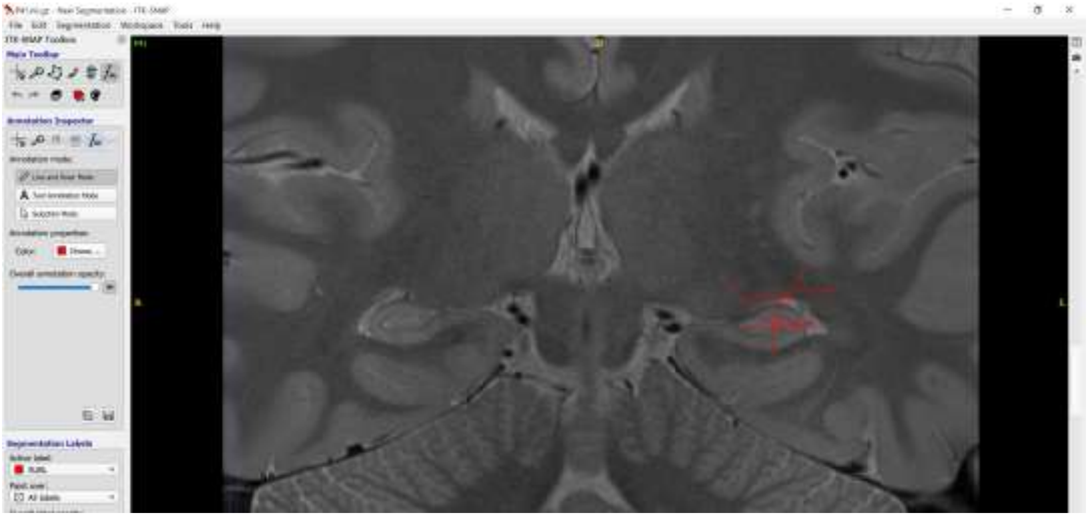

5. Using the segmentation tools, outline each subfield
   a. The polygon tool is useful for outlining
      i. With the oblique tool, close the shape and click accept at the bottom right of the screen to complete the label
   b. The paintbrush is useful for smaller edits
      i. Under the paint over drop down menu, choose Clear Label in order to not change adjacent subfield labels
      ii. With the paint tool, hold down the right mouse button to erase/remove pixels
   c. All subfields should be contiguous—no gaps between boundaries
      i. If there is a gap in the hippocampal fissure, a cyst or sulcal cavity, do not label it

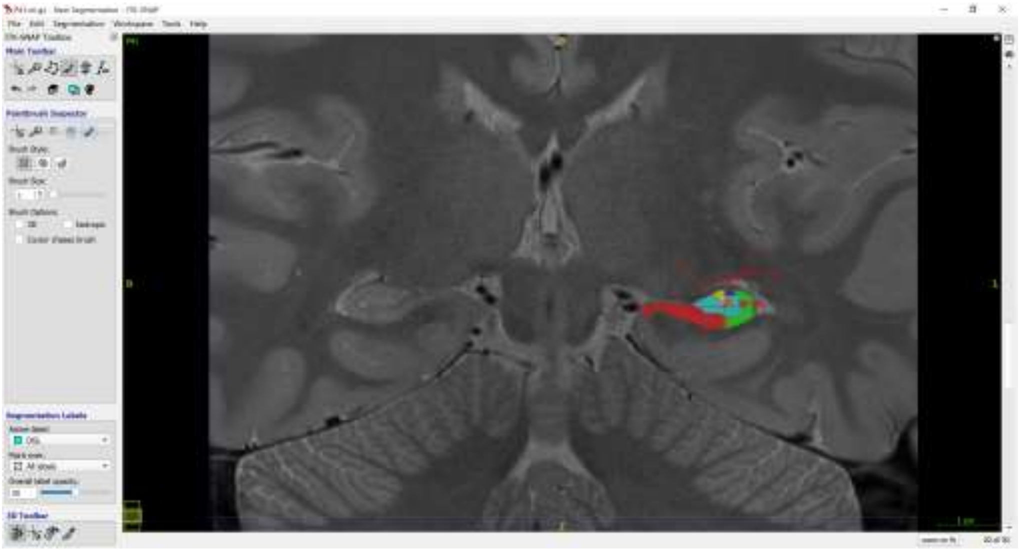

6. Remove the lines—Click select in the Annotation pane, click select all in bottom right, click clear
7. Edit segmentation to ensure labeling all voxels—the paintbrush is useful for this

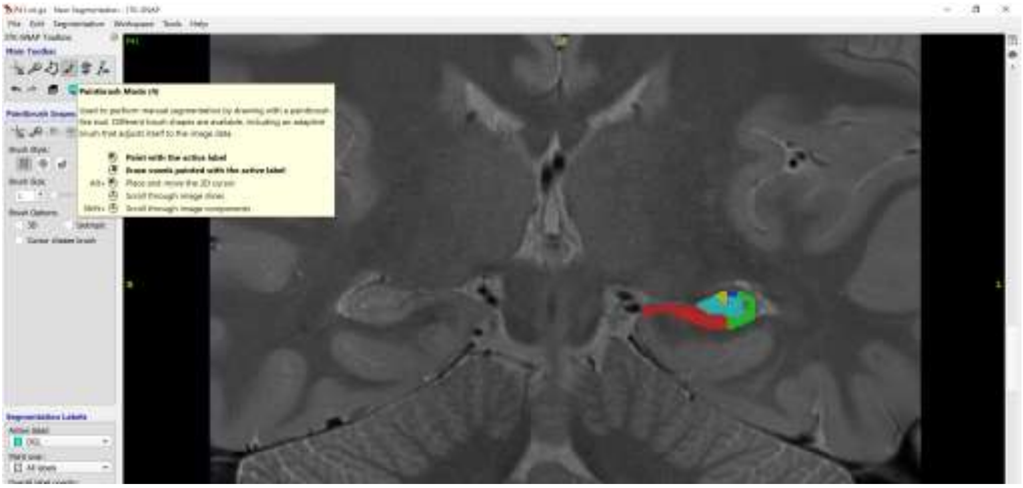

#### Endfolial Pathway Rule

The modified rule uses the same definitions for the subiculum-CA1, CA1-CA2 and CA2-CA3 boundaries (Line 1-3). The additional Endfolial Pathway Rule (Line 4e-6e) defines a different boundary for CA3-dentate gyrus.

**<u>Protocol Description:</u>** The first 3 reference lines are the same as explained in the heuristic rule. (**1**) Line 1 should be anchored at the opening of the hippocampal fissure (**a,** the "arm pit") and extended to the most lateral, outside edge of the alveus (white matter structure) of the CA1 sector. (**2**) Line 2 is then placed perpendicular to the middle of Line 1, extending from the most superior edge of the hippocampus to the parahippocampal white matter. (**3**) An additional vector is extend from the point of bisection at 30° to the lateral side. The CA3-dentate gyrus boundary is determined by a different heuristic that approximates the location of the endoflial pathway, labeled **lines 4e-6e**.

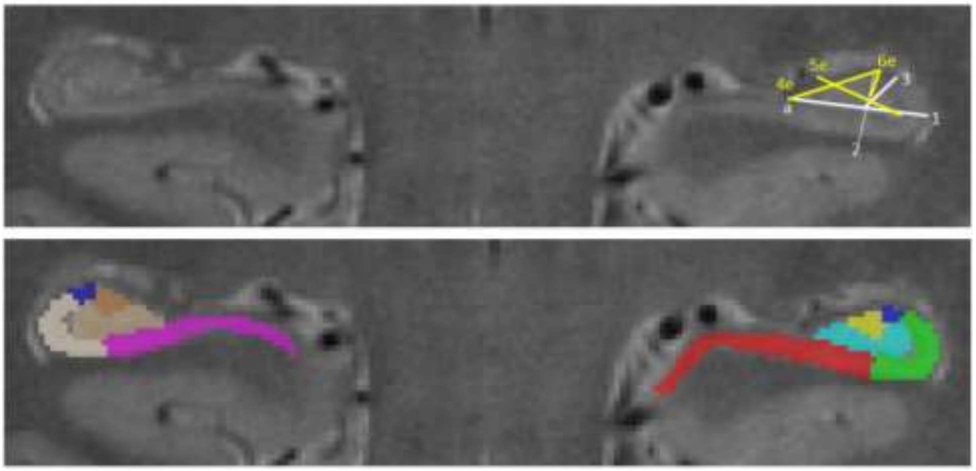

| | | |
|---|---|---|
| **1.** Line 1 should be anchored at the opening of the hippocampal fissure adjacent to the superior edge of the subiculum (**a,** the "arm pit") and extended to the most lateral, outside edge of the alveus (white matter structure) of the CA1 sector. | 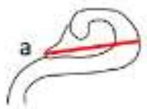  | 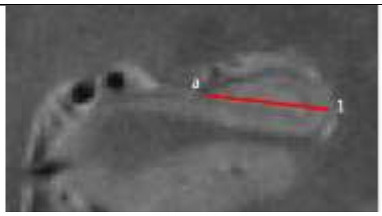  |
| **2.** Line 2 is then placed perpendicular to the middle of Line 1, extending from the most superior edge of the hippocampus to the parahippocampal white matter. | 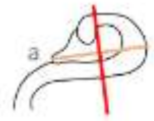  | 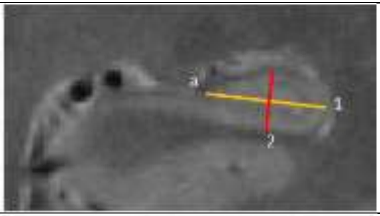  |
| **3.** An additional vector is extend from the point of bisection at 30° to the lateral side. | 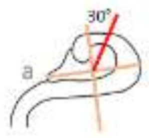  | 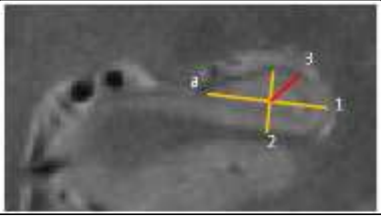  |

| | | |
|---|---|---|
| **4e.** Extend Line 4e from the most medial point of the dentate gyrus (placed adjacent to the subiculum) to the most superior, middle point of the CA fields. Note that depending upon the morphometry of the hippocampus, the dentate gyrus may extend medial to the opening of the hippocampal fissure (i.e., not the same position as **a**, "the armpit"). | 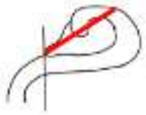 | 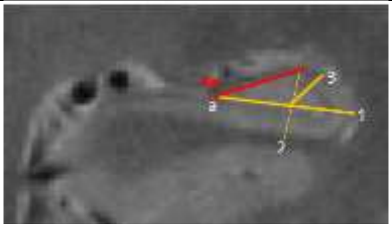  |
| **5e.** Find the half point of Line 4e; extend Line 5e from the most lateral point of the SLRM (internal boundary between dentate gyrus and CA1) to the medial edge of the dentate gyrus, bisecting the half point of Line 4e. Ignore cysts for this line placement. | 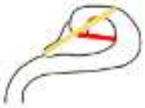 | 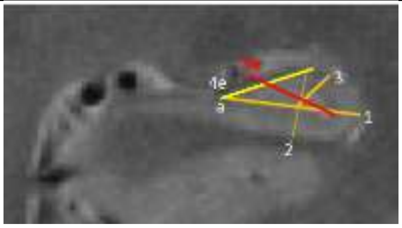  |
| **6e.** Find the half point of Line 5e, extend Line 6e from the half point to the superior edge of CA, aligned perpendicular to Line 5e. | 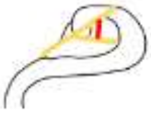 | 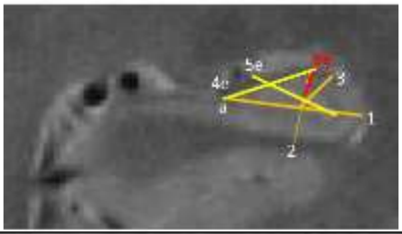  |
| Lines 5e and 6e create a wedge to the superior medial edge, and this is labeled CA3. Note that the CA2 label should not be changed and CA3 is labeled in the adjacent tissue. The remainder of the volume is dentate gyrus. | 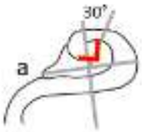  | 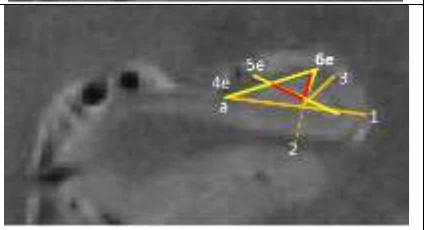  |

**<u>Manual Tacing Example in ITKSnap:</u>**

1. Follow the steps for the Geometric Heuristic rule for Lines 1-3
2. Label the subiculum, CA1 and CA2
3. Remove all lines—Select all and clear

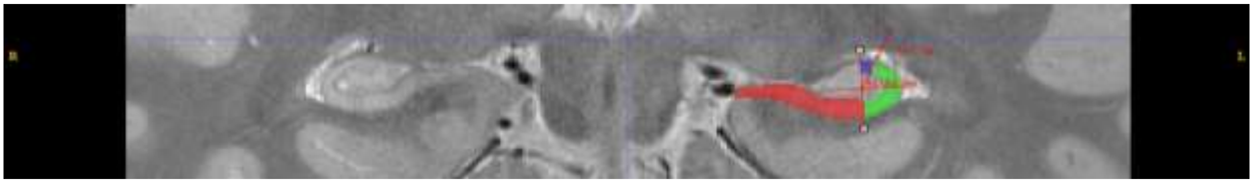

4. With the Annotation tool
   a. Draw Line 4e from the most medial edge of the dentate gyrus (adjacent to subiculum) to the most superior edge
   b. Draw Line 5e from the half point of 4e to the most lateral point of the SRLM
   c. Draw Line 6e from half point of 5e to the superior SRLM
5. Label CA3 and dentate gyrus

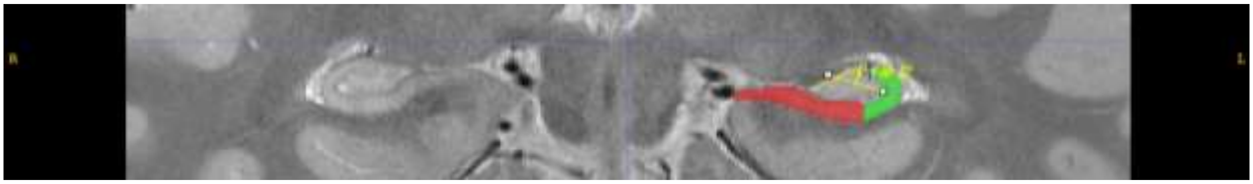

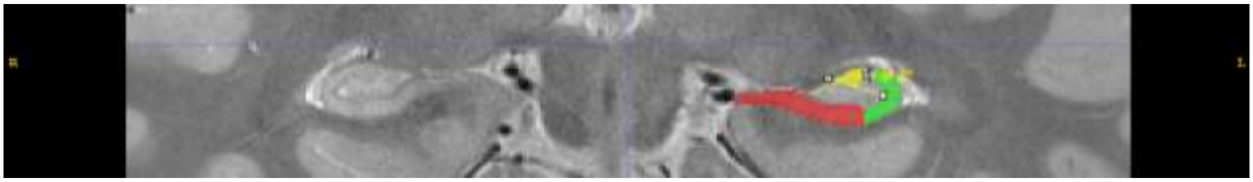

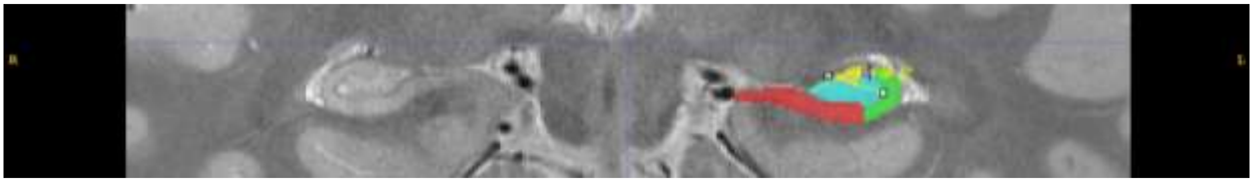

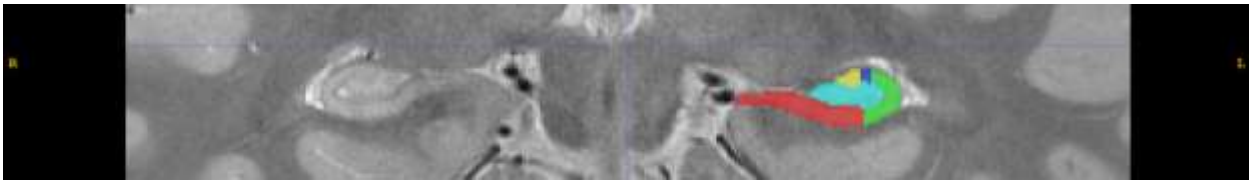

Appendix B: Description of Outer Boundaries

**Outer boundaries protocol**

Ana Daugherty and Renaud La Joie

On behalf of the Boundary Working Group

Last updated: July 2019

---

**Table of contents**

I. Anterior
II. Posterior
III. Dorsal
IV. Ventral
V. Medial
VI. Lateral
VII. Blood vessels
VIII. CSF and cysts

---

**Notes:**

All protocol rules relate to viewing the hippocampal body in the coronal view, and assume that boundaries will be drawn independently for the left and right hemispheres (such that asymmetries in boundary placement are possible).

All figures are T2 coronal images acquired perpendicular to the longitudinal axis of the hippocampus and are centered on the hippocampal body, unless otherwise specified.

#### I. ANTERIOR

The disappearance of the uncus is used to determine the transition from hippocampal head to body. The uncus lies on the medial edge of the hippocampal head, and on posterior head slices, it is often connected to the rest of the hippocampus via the fimbria only (See FIG 1). The anterior boundary of the hippocampal body (HB), i.e., the anterior-most slice included in the HB, is the **first slice posterior to the last visualization of the uncus**.

Note that head misalignment and/or anatomical differences between hemispheres may lead to differences in defining the anterior boundary of the HB in each hemisphere (usually differing by one slice in images with 2mm slice thickness). See FIG 1 and FIG 2 for examples.

Importantly, conservative judgement should be used and slices showing partial voluming of the uncus should be categorized as a "head slice" and not part of the HB. Partial voluming refers to situations in which a voxel represents an average of two or more tissue types (e.g., both grey matter and cerebrospinal fluid). When determining partial voluming, consult the previous contiguous slice to evaluate if the portion of tissue falls within the uncus region. See FIG 3 for an example.

**FIG 1**. Top left: Hippocampal head displaying the uncus in both hemispheres. Top right: Slice showing the disappearance of the uncus in the left hemishphere, and as such, the anterior-most slice of the hippocampal body in the left hemisphere. T2-weighted image, resolution 0.39 x 0.39 x 2mm. Bottom: Sketch adapted from Duvernoy et al., 2013 with portions of the uncus labeled as 4 and 6

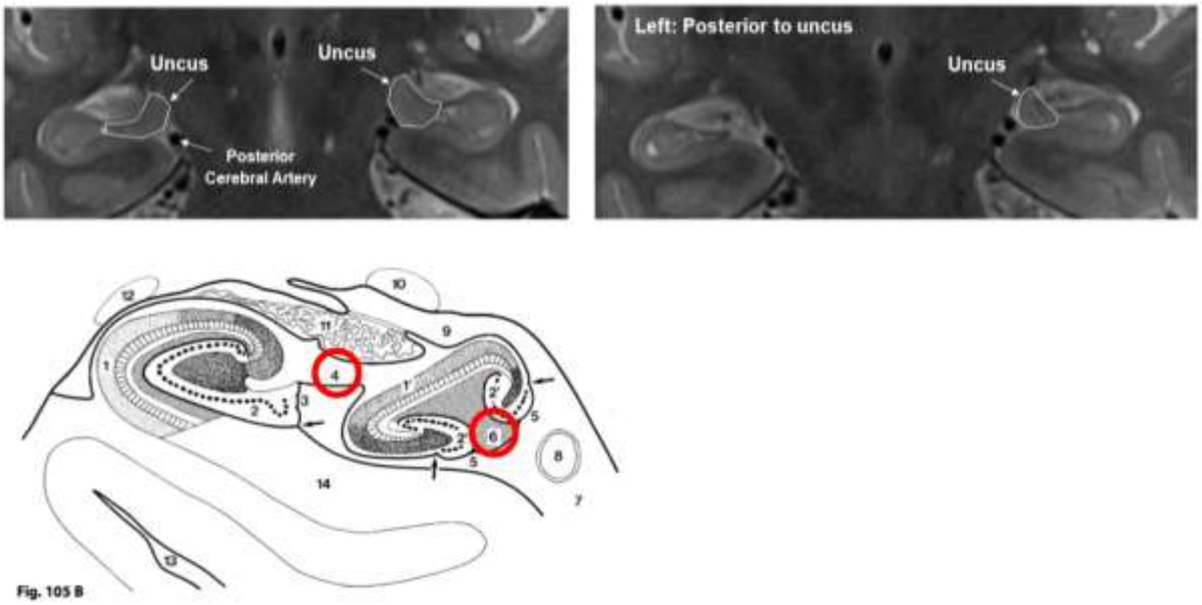

**FIG 2**. Second example demonstrating hemispheric asymmetry in the definition of the anterior boundary of the hippocampal body. Slices proceed from anterior to posterior. T2-weighted image, resolution 0.39 x 0.39 x 2mm.

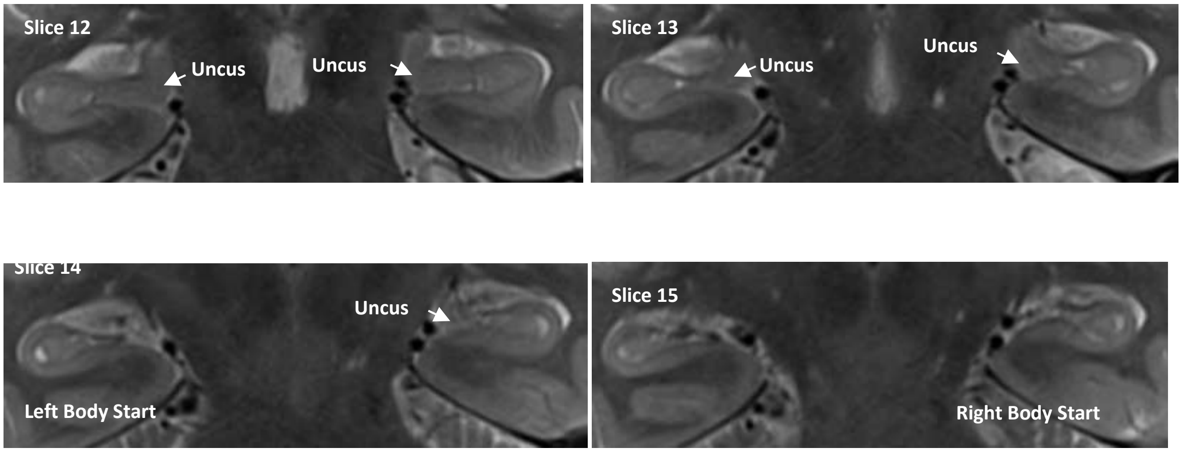

**FIG 3**. Example demonstrating partial voluming of the uncus. Slices proceed from anterior to posterior. T2-weighted image, resolution 0.39 x 0.39 x 2mm.

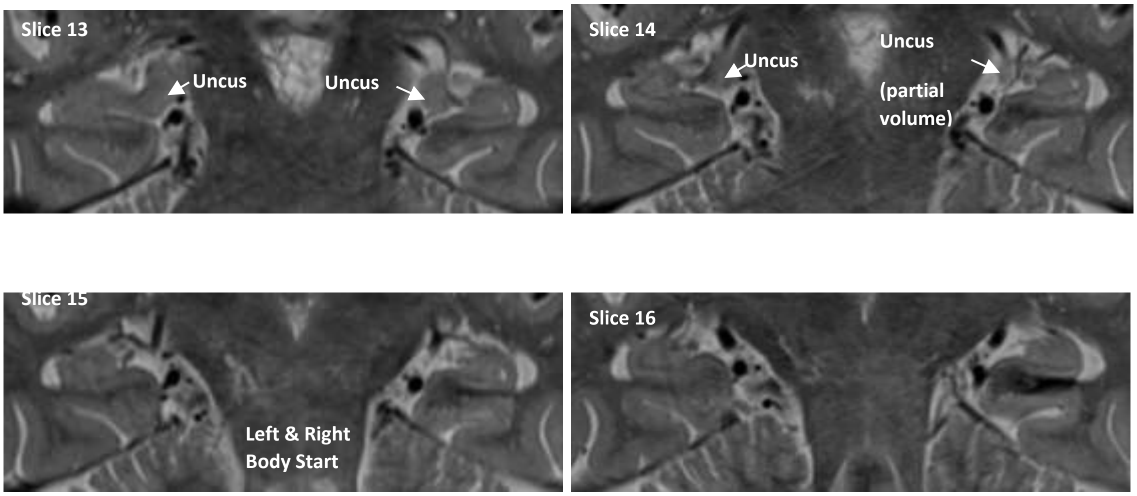

**FIG 4**. Example demonstrating partial voluming of the uncus (slice 10) on an image with motion artifacts. Slices proceed from anterior to posterior. T2-weighted image, resolution 0.39 x 0.39 x 3mm.

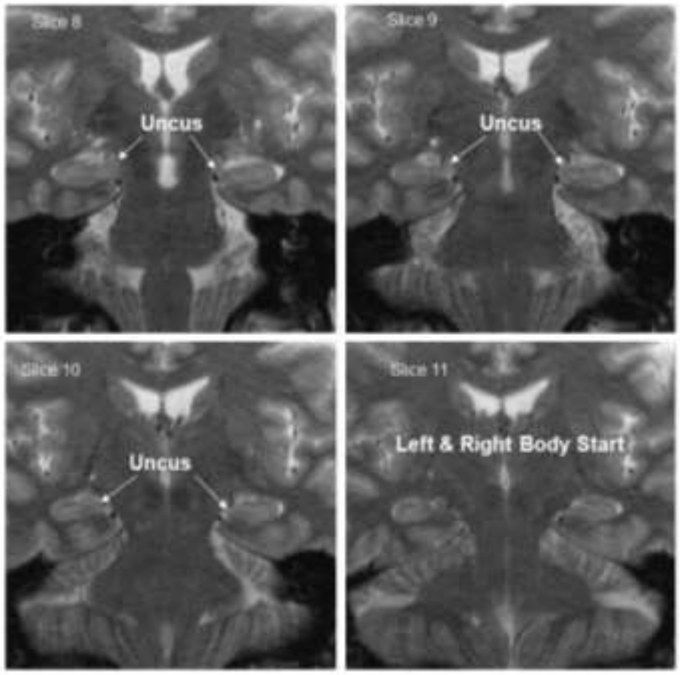

⇒ Back to table of contents

#### II. POSTERIOR

The posterior boundary of the HB, i.e., the posterior-most slice included in the HB, is determined by a single landmark: **the posterior-most slice on which the colliculi are visible**. The colliculi, also called the lamina quadrigemina, refer to the superior and inferior colliculi of the midbrain. On a T2-weighted coronal image, they appear as four hypointense (i.e., dark) round structures along the midline of the brain, posterior to the cerebral peduncles. When all four colliculi are visualized in the coronal plane, their arrangement may appear like a “butterfly”. On 2-mm coronal slices, the colliculi will be visualized on 2-3 slices (see FIG 5 and FIG 6 for examples).

Segmentation of the subfields within the hippocampal body only stops when the colliculi have entirely disappeared. Note that partial voluming of the colliculi may occur, as shown in FIG 7-9, and that any visualization -- even partial -- should be considered a body slice when defining the posterior boundary of the HB.

The image acquisition of the hemispheres may be misaligned, and similar to the anterior-most slice, the posterior-most slice of the hippocampal body may differ between hemispheres. Only one colliculi in a hemisphere need be visualized to be considered the most-posterior slice of body in that hemisphere. It is worth noting that because the colliculi are structures that fall along the midline, this landmark will be less sensitive to differences between hemispheres as the anterior ranging rule. The use of this landmark is a conservative posterior definition of the HB, ensuring exclusion of the tail. In some cases, this may result in a portion of the HB mislabeled as hippocampal tail.

**FIG 5**. Left: Final posterior slice of the hippocampal body displaying the colliculi. This landmark coincides with others that are commonly reported in the literature: the crus fornix and the "tear drop" shape of the hippocampal body. Right: Colliculi are no longer visible, and as such this slice is considered the first slice of the hippocampal tail. T2-weighted image, resolution 0.39 x 0.39 x 2mm.

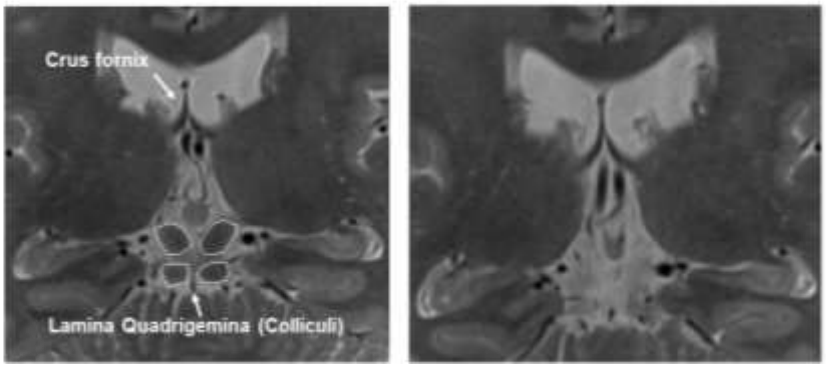

**FIG 6**: Series of coronal slices proceeding from anterior to posterior, demonstrating the definition of the posterior boundary of the hippocampal body. T2-weighted image, resolution 0.39 x 0.39 x 2mm.

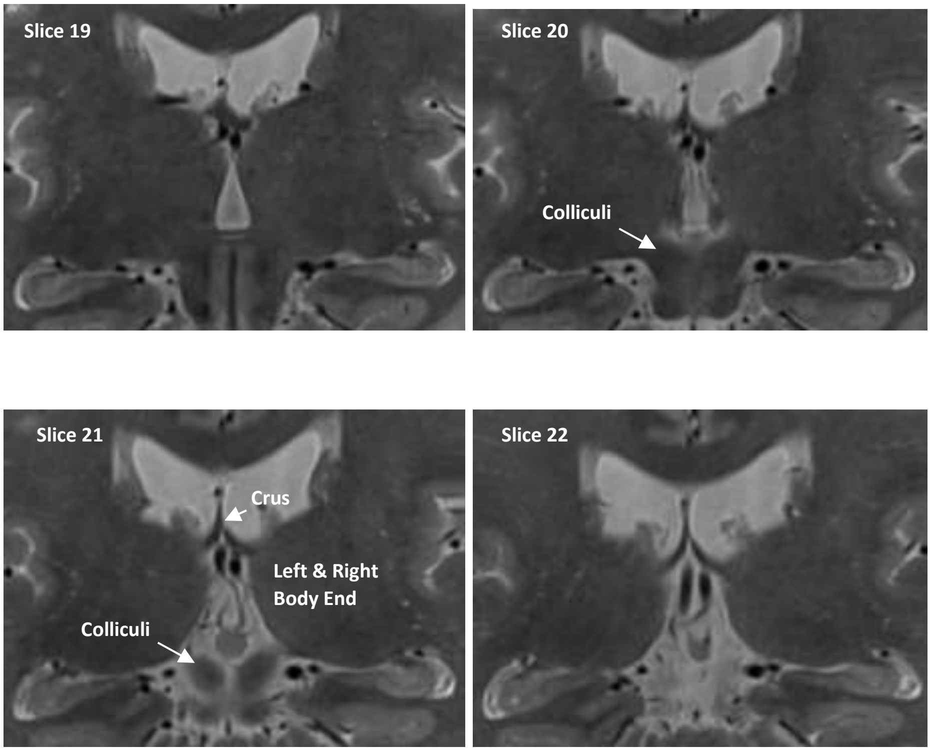

**FIG 7**: Series of coronal slices proceeding from anterior to posterior, demonstrating partial voluming of the colliculi and how even a partial visualization of the colliculi should be interpreted as a body slice. Note that the superior colliculi in each hemisphere are visualized with a portion of dark tissue that is consistent with the prior slice and this is interpreted as partial voluming. As such, the most-posterior body is on slice 22. T2-weighted image, resolution 0.39 x 0.39 x 2mm.

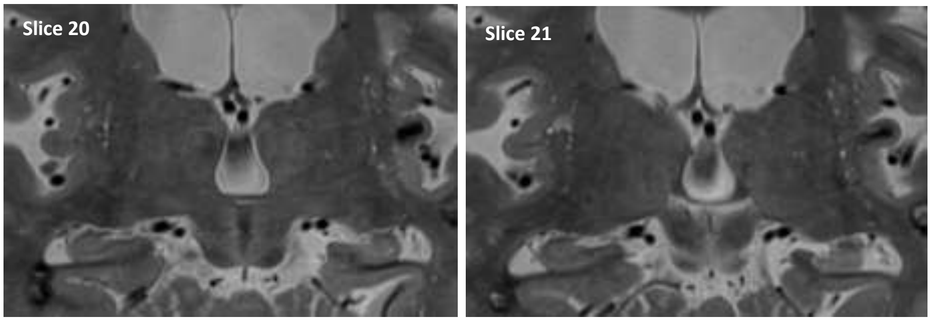

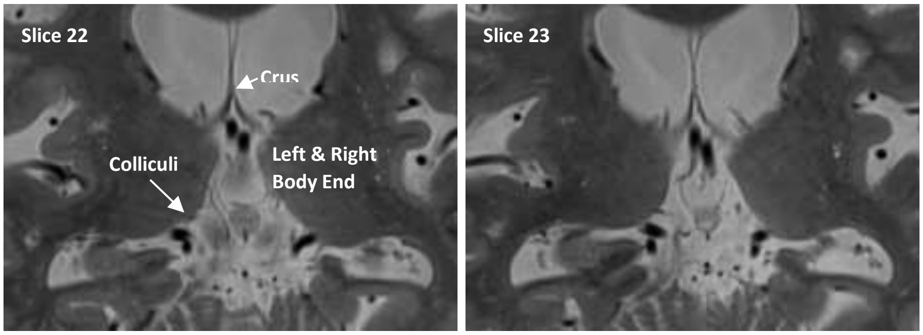

**FIG 8**: Series of coronal slices proceeding from anterior to posterior, demonstrating partial voluming of the colliculi with motion arifact. Note that the inferior colliculi in each hemisphere are visualized with a portion of dark tissue that is consistent with the prior slice and this is interpreted as partial voluming. As such, the most-posterior body is on slice 15. T2-weighted image, resolution 0.39 x 0.39 x 3mm.

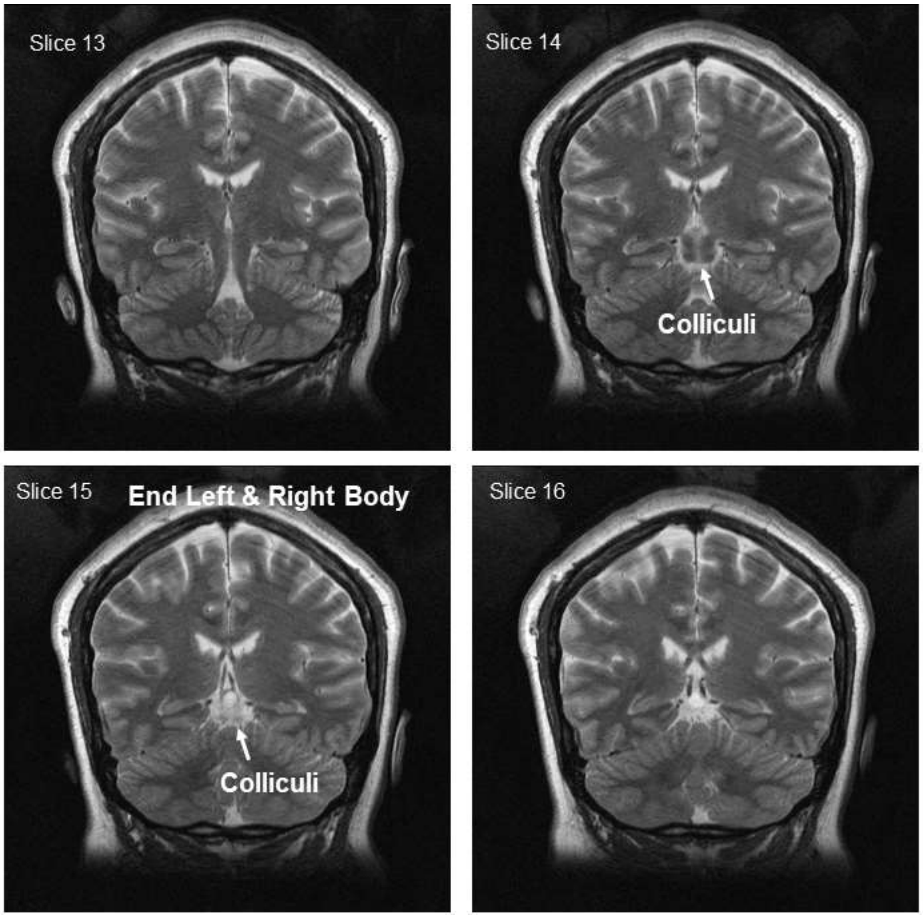

**FIG 9**: Series of coronal slices proceeding from anterior to posterior, demonstrating partial voluming of the colliculi with head misalignment at acquisition. Note that the inferior colliculi in each hemisphere are visualized with a portion of dark tissue that is consistent with the prior slice and this is interpreted as partial voluming. As such, the most-posterior body is on slice 23. T2-weighted image, resolution 0.39 x 0.39 x 2mm.

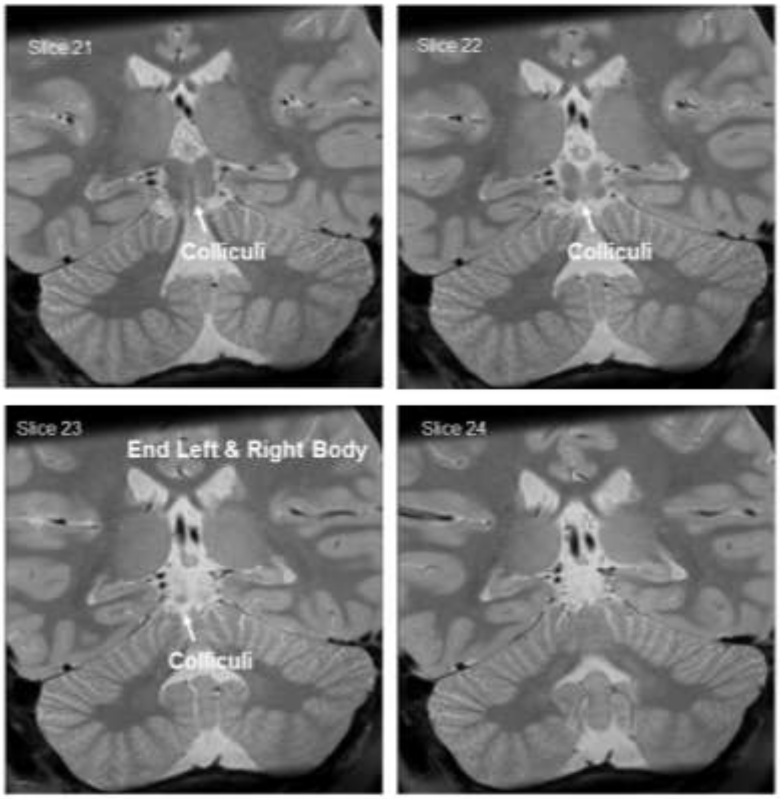

The boundary definition should be made foremost by visibility of the colliculi, even in instances of hemispheric or subtle head pitch misalignment. As can be seen in FIG 5-7, the posterior-most slice on which the colliculi are visualized often (but not always) shows the crus fornix and the HB often has the typical "tear drop" shape on this slice. If HB-like slices--on which the dentate gyrus is still visualized and the hippocampus has a tear drop shape--appear posterior to the last visualization of the colliculi, the researcher may suspect scan misalignment. On a scan aligned to be perpendicular to the hippocampal long axis, the inferior and superior colliculi will be visualized at the same time within the slice sequence, whereas on a misaligned scan, the inferior or superior colliculi (depending upon the direction of pitch) will be visualized out of sync. If the image acquisition is misaligned, and the image cannot be resliced to correct the alignment, then additional landmarks may be considered when defining the posterior boundary of the HB. The appearance of the crus of the fornix concurrent with a “tear drop” like shape of the hippocampus (including visualization of the dentate gyrus) should be considered the posterior-most slice of the hippocampal body.

⇒ Back to table of contents

**III. DORSAL**

The dorsal boundary is defined as the interface between the gray matter tissue of the HB and the white matter of the alveus and fimbria. In contrast to the low-intensity band of white matter sitting on top of the HB, the gray matter appears as high-intensity voxels on T2 MRI.

**FIG 10**: The alveus and fimbria in a coronal view in the anterior hippocampal body. Left: Sketch adapted from Duvernoy, 2005. Right: T2 MRI, image resolution: 0.42 x 0.42 x 2mm.

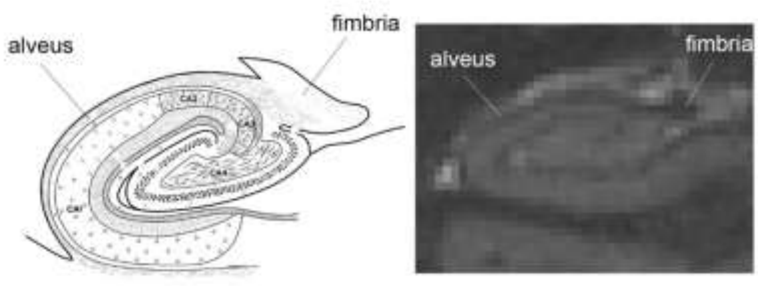

With respect to subfield segmentation, **both the alveus and fimbria must be excluded from any regions of interest**. They receive inputs from different hippocampal subregions and cannot be clearly assigned to one specific subregion. In addition, their inclusion would pose additional segmentation issues at the level of the crux of the fornix that would also contribute to reliability issues.

Therefore, the **hypointense (dark) voxels belonging to the fimbria** as well as the **thin hypointense band covering the hippocampal body on the dorso-lateral boundary** corresponding to the alveus **have to be excluded.**

**FIG 11**: Coronal view of the hippocampal body demonstrating the dorsal boundary along the anterior-posterior axis. The red line indicates the dorsal-most voxels to be included in the HB. Note: In the posterior part of the hippocampal body, the dentate gyrus extends medially. This extension has to be included in the hippocampal segmentation (panels D and E). Image resolution: 0.42 x 0.42 x 2mm.

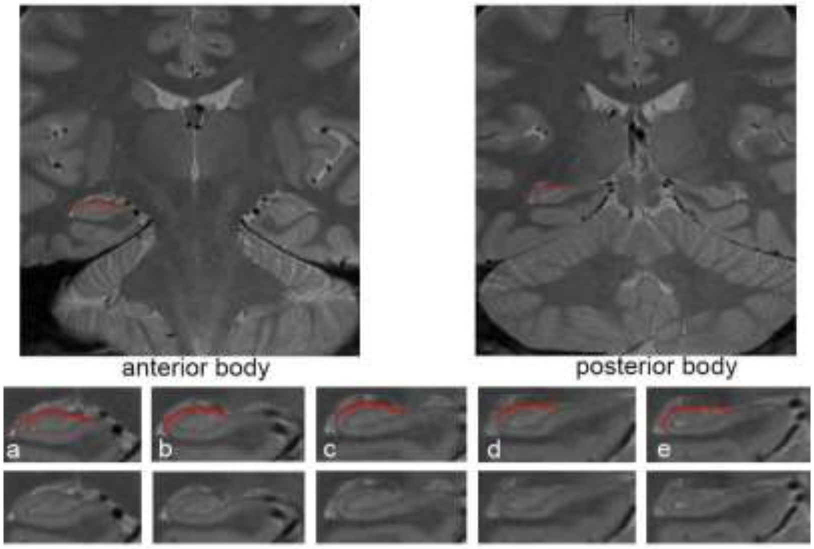

In contrast, the **hypointense voxels bordering the subiculum superiorly, and continuous with the stratum-lacunosum moleculare (SRLM) - are considered part of the hippocampus and included in the segmentation.**

**FIG 12**: Coronal view of the hippocampal body depicting incorrect and correct labeling of the hypointense voxels bordering the subiculum. Image resolution: 0.42 x 0.42 x 2mm.

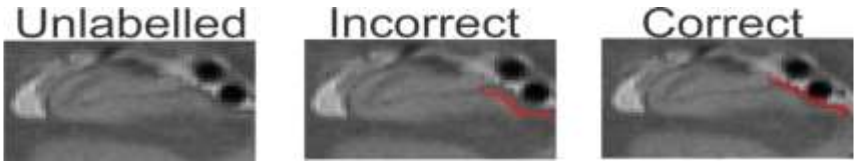

⇒ Back to table of contents

#### IV. VENTRAL

At the level of the HB, the ventral hippocampus is bordered by the white matter sitting above the parahippocampal gyrus. Following this logic, the ventral boundary is simply taken to be the **border between the grey matter of the hippocampus and the white matter that occurs inferior to it**, such that white matter is excluded from the definition of the HB.

**FIG 13**: Ventral boundary depicted in a coronal view of the hippocampal body. The red line indicates the ventral-most voxels to be included in the definition of the HB. Image resolution: 0.42 x 0.42 x 2mm.

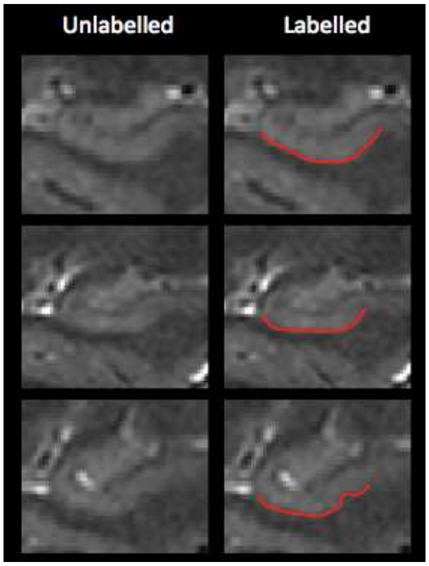

**FIG 14**: Coronal view of the hippocampal body demonstrating ventral boundary, adapted from Duvernoy et al., 2013.

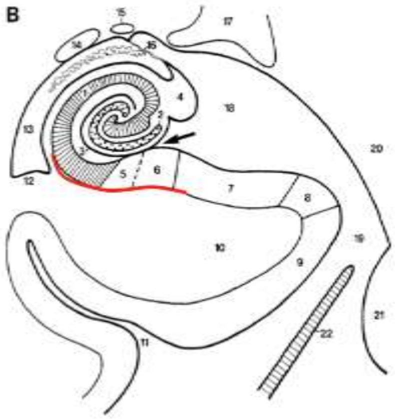

⇒ Back to table of contents

**V. MEDIAL**

At the level of the HB, the medial portion of the hippocampus corresponds to the subiculum and must be separated from the entorhinal, perirhinal and parahippocampal cortices. This separation is inherently difficult since few morphological or contrast differences exist to aid in differentiating the subiculum from the entorhinal/perirhinal/parahippocampal cortices.

The EADC-ADNI Harmonized Protocol (HaRP, http://www.hippocampal-protocol.net) defines the medial boundary by '*tracing an irregular line continuing from the visible interface with the WM of the parahippocampal gyrus, to the ventro-medial aspect based on the continuity of the boundary as detected from morphological details and GM intensity*'.

**FIG 15**: Top: **HaRP (<u>not HSG!</u>) medial boundary** depicted on a coronal MRI slice at the level of the hippocampal body. Bottom: HaRP medial boundary depicted on a coronal sketch of the hippocampal body adapted from Duvernoy et al, 2013. Image resolution: 0.42 x 0.42 x 2mm.

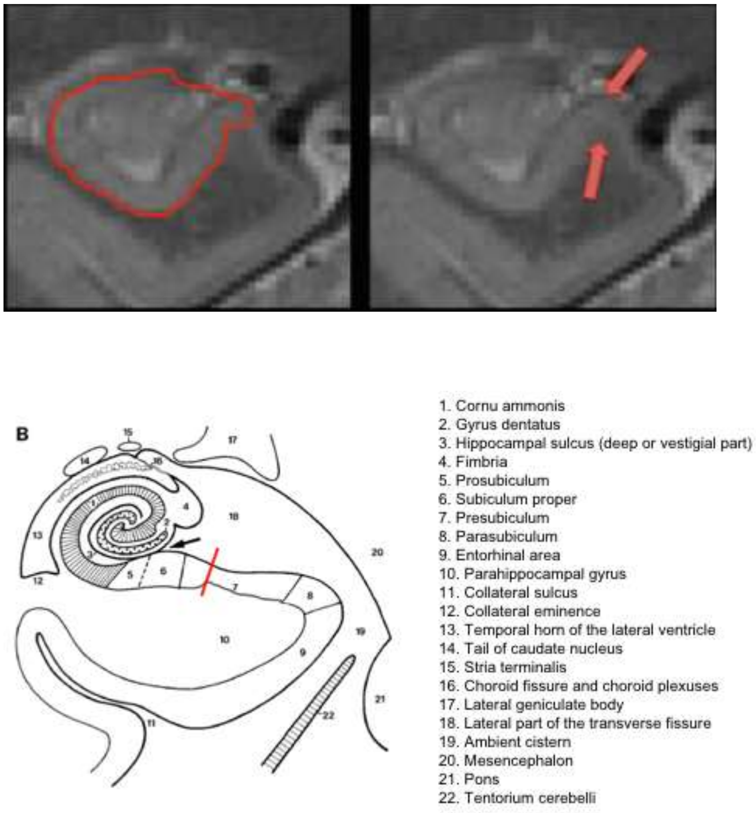

The HSG protocol defines the medial boundary of the hippocampus differently than HaRP. Namely, at the level of the HB, **the medial boundary is defined as the most <u>medial</u> corner of the parahippocampal gyrus** and typically coincides with the termination of the subicular complex. See Figs 16-19 for examples. The boundary is placed at the **level of maximum curvature in the cortical ribbon**, just before it runs parallel to the tentorium cerebelli (i.e. the meninges indicated by blue arrows in Fig 16). Note that the boundary is traced perpendicular to the cortical ribbon (see Fig 19). It is important to note that this boundary **would more likely allow for the inclusion of presubicular and parasubicular cortices** than the HaRP protocol (see lower panel of Fig 16).

Depending on the orientation of the image acquisition and/or the shape of the hippocampus, the location of this boundary may be ambiguous. If, for example, the hippocampus has a rounder shape as in the top portion of Fig 16, and shows multiple areas of curvature, the medial boundary of the hippocampus is set at the medial point, where the cortical ribbon curves downwards and sometimes laterally, just before running parallel to the tentorium cerebelli.

**FIG 16**: Top: **Current protocol's definition of the medial boundary** at the level of the hippocampal body on a coronal MRI slice. The medial boundary (red arrow) is defined as the most medial point of the cortical ribbon before the cortex runs parallel to the tentorium cerebelli (meninges, indicated by the blue arrows). Bottom: Current protocol's definition of the medial boundary displayed on a coronal sketch of the hippocampal body from Duvernoy et al, 2013. Image resolution: 0.42 x 0.42 x 2mm.

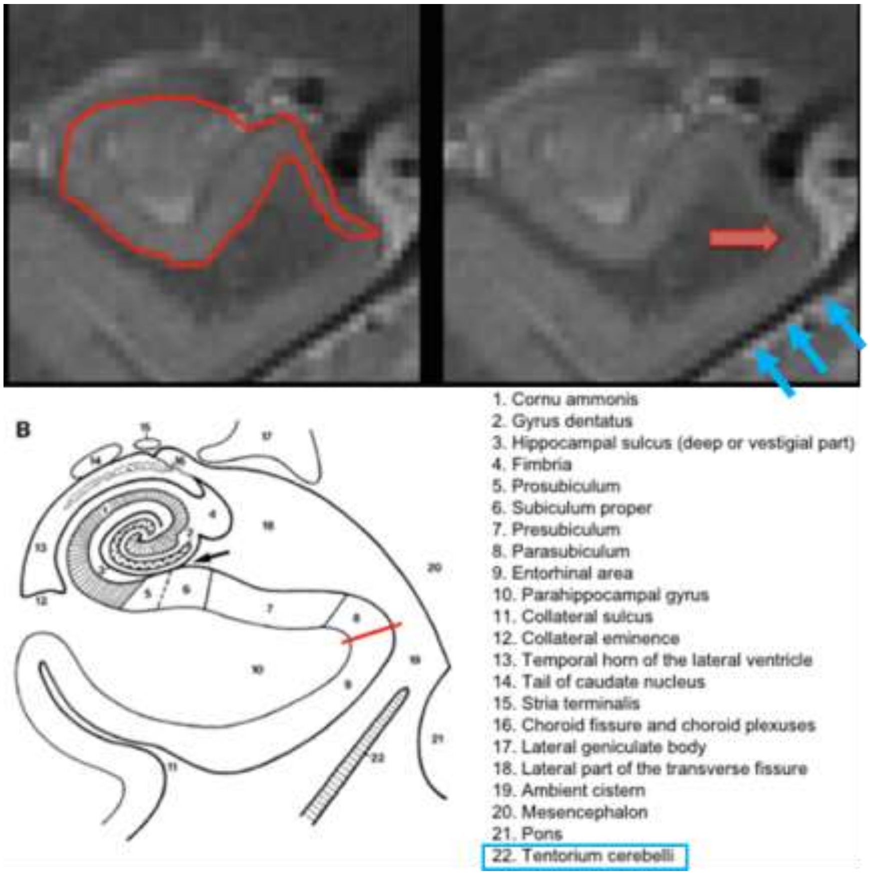

At times, the observed thinning of the subicular cortex and lower MRI contrast may limit tracing ability, yet most high resolution images offer the spatial resolution to complete tracing to this <u>most medial</u> point.

Depending on the subject anatomy and the image orientation, the most medial aspect of the medial temporal lobe can be flat (~vertical, see Subjects 2 and 3 from figure 17l) and the level of maximal curvature might correspond to the most supero-medial point of the parahippocampal gyrus.

**FIG 17**: Examples of defining the medial boundary at the level of the hippocampal body. The medial boundary is defined as the area of maixma curvature of the parahippocampal gyrus (most medial point for Subject 1; or superomedial corner in case the most medial aspect of the medial temporal lobe is flat/vertical, see Subjects 2-3), where the cortex curves downwards before running parallel to the tentorium cerebelli. Coronal slices progress from anterior (top row) to posterior (bottom row). Image resolution: 0.42 x 0.42 x 2mm.

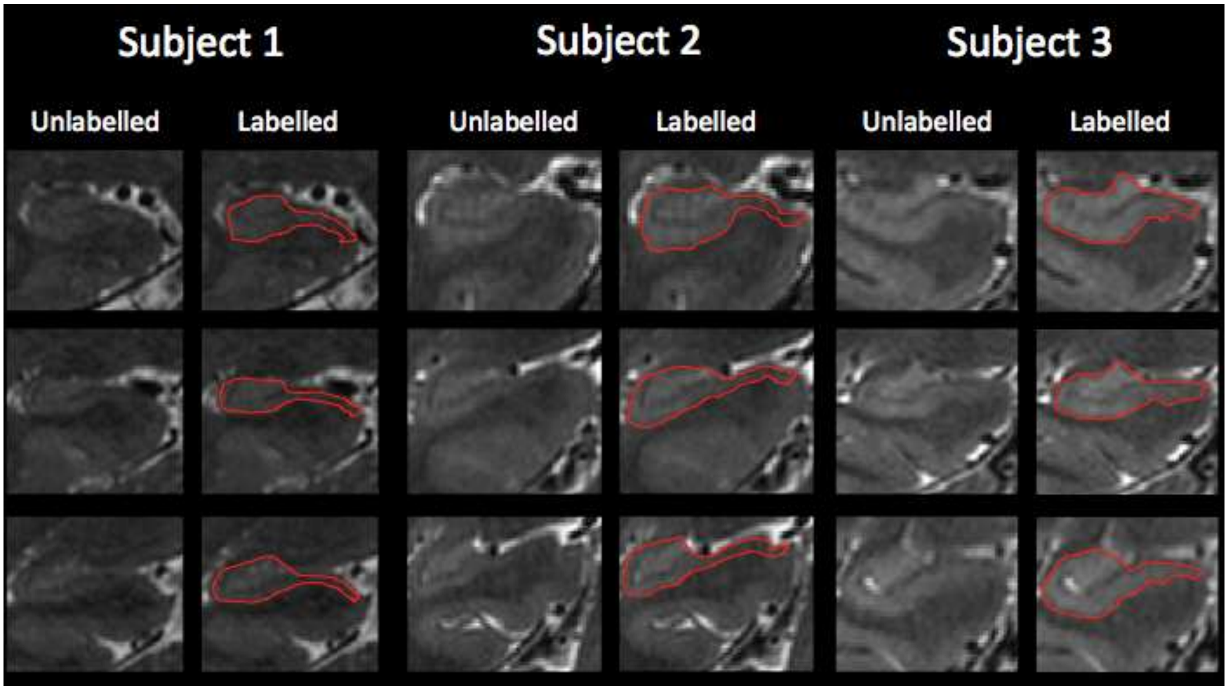

**FIG 18**: Examples of correct and incorrect labeling of the medial boundary depicted at the level of the hippocampal body. Correct labeling demonstrates that the boundary is placed at the most (supero)medial point of the parahippocampal gyrus, at its maximum curvature before turning downwards to run parallel to the tentorium cerebelli. Image resolution: 0.42 x 0.42 x 2mm.

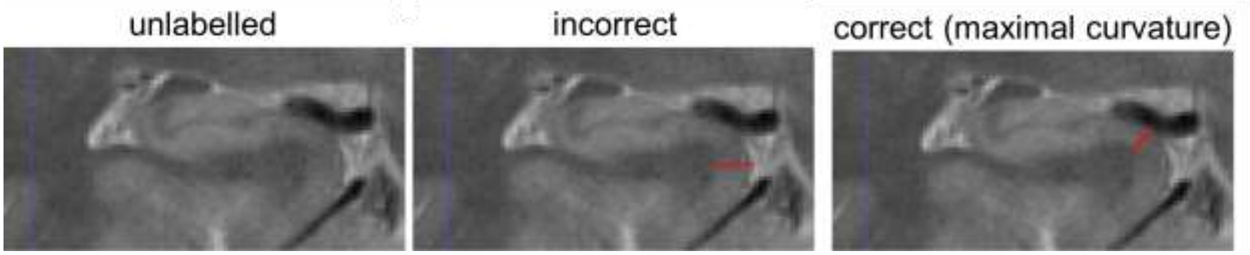

Furthermore, care should be taken to draw this border **orthogonal (perpendicular) to the cortical ribbon/pial surface**.

**FIG 19**: Segmentation of the medial boundary with cortical ribbon

highlighted to demonstrate correct orthogonal placement (the medial boundary of the hippocampus is set perpendicular to the dotted line that represents the pial surface at the level of maximal curvature)

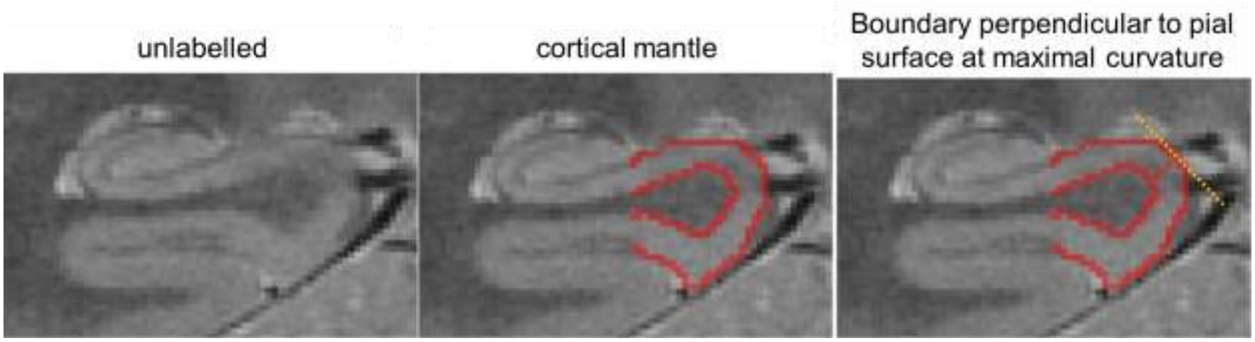

⇒ Back to table of contents

#### VI. LATERAL

The lateral boundary is the same as that presented in the "Dorsal Boundary" section. That is, the lateral boundary is taken to be the border between the gray matter tissue of the HB and the white matter of the alveus/fimbria, such that the **alveus and fimbria should not be included in the segmentation**.

**FIG 20**: Top: Examples of the lateral boundary drawn on several coronal slices at the level of the hippocampal body on MRI. Similar to the dorsal boundary, the lateral boundary is defined as the interface between the hippocampal grey matter and extra-hippocampal white matter. Image resolution: 0.42 x 0.42 x 2mm. Bottom: Lateral boundary displayed on a coronal sketch of the hippocampal body adapted from Duvernoy et al, 2013.

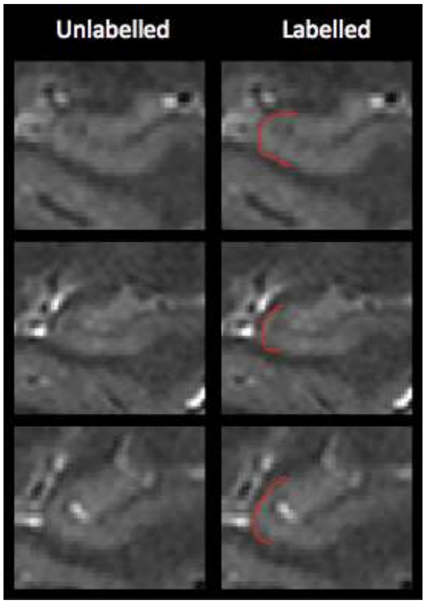

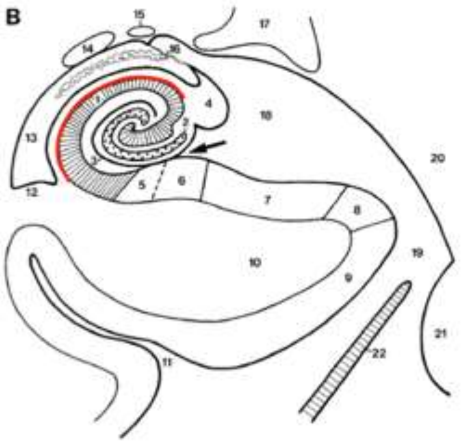

1. Cornu ammonis
2. Gyrus dentatus
3. Hippocampal sulcus (deep or vestigial part)
4. Fimbria
5. Prosubiculum
6. Subiculum proper
7. Presubiculum
8. Parasubiculum
9. Entorhinal area
10. Parahippocampal gyrus
11. Collateral sulcus
12. Collateral eminence
13. Temporal horn of the lateral ventricle
14. Tail of caudate nucleus
15. Stria terminalis
16. Choroid fissure and choroid plexuses
17. Lateral geniculate body
18. Lateral part of the transverse fissure
19. Ambient cistern
20. Mesencephalon
21. Pons
22. Tentorium cerebelli

⇒ Back to table of contents

#### VII. BLOOD VESSELS

There are several blood vessels within and close to the hippocampal formation. Of interest are especially the posterior cerebral artery and the basal vein.

**FIG 21**: Coronal sketch at the level of the hippocampal body depicting blood vessels neighboring the hippocampus. Adapted from Duvernoy, 2005.

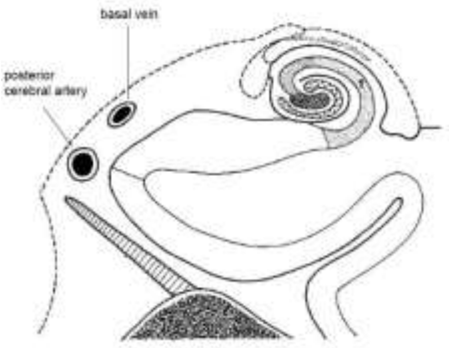

There are also smaller intrahippocampal branches from these vessels connecting the inner hippocampus. Blood vessels appear hypointense (dark) in T2 weighted images (note, however, that they are bright in T1 images). The big vessels close to the subiculum can be especially problematic during segmentation as they can cause signal drop out or may cause hippocampal anatomy to appear slightly different. **Vessels and/or signal drop out due to vessels has to be excluded from the segmentation.**

**FIG 22**: Two examples of arteries close or within the hippocampal body. The left column is unlabeled and the right column indicates proper segmentation according to the current protocol. In the first example (A and B), the artery is touching the grey matter, but in the second example (C and D) the artery appears within the grey matter and causes signal dropout. Image resolution: 0.42 x 0.42 x 2mm.

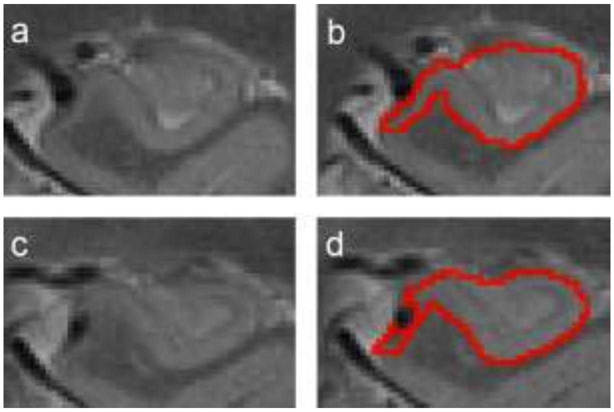

⇒ Back to table of contents

### VIII. CSF and CYSTS

In T2 weighted images, CSF cysts appear as hyperintense (bright) regions (though note that they appear dark in T1 images). Cysts are observed regularly on higher resolution scans. They are often located in the vestigial sulcus along the longitudinal axis of the hippocampus, and a majority of them appear in the ventrolateral flexion points of CA1 (see below; also see Veluw et al., 2013).

In general **we recommend segmenting the cysts separately** either before or following the delineation of the other subfields. Cysts will be segmented using a separate label.

They **can often be followed on consecutive slices** [rule 1]. However, especially on images with anisotropic voxel size and thicker slices this does not have to be the case.

Cysts occurring in the HB **should be removed if the tracer is sure that they represent CSF**. Only clusters that consist of **at least two or more contiguous voxels (in any direction) that are considerably brighter than their surroundings will be labeled** [rule 2]. Often the voxels in the center of the cyst are brightest and fade out towards the edges (see FIG 13 from prior section).

Therefore, **cysts will be segmented based on hyperintensity, presence on adjacent slices, and number of voxels**. However, not every cyst might have all three properties.
**FIG 23**: Unlabeled (left column), incorrectly labeled (middle column), and correctly labeled (right column) CSF cysts. Slices are from the same subject and move from anterior (top row) to posterior (bottom row) portions of the hippocampal body. Note that for correct labeling, only the center, brightest voxels of the cysts are labeled as CSF. Image resolution: 0.42 x 0.42 x 2mm.

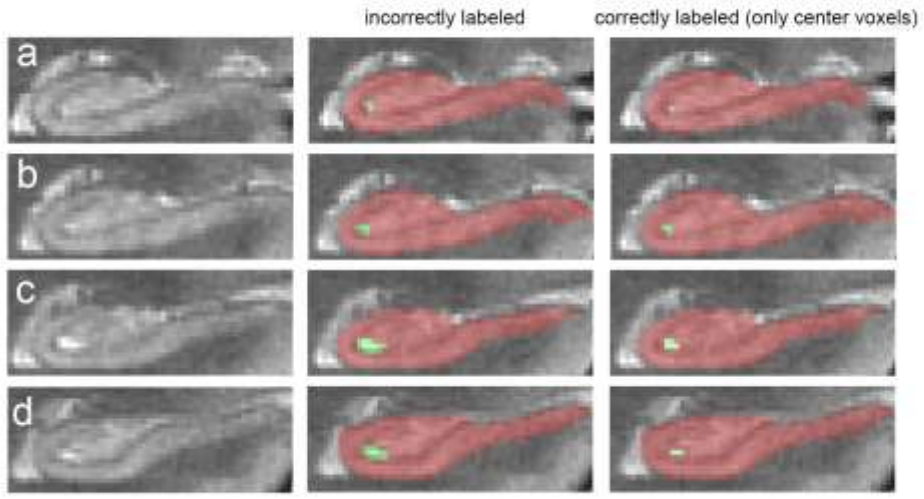

⇒ Back to table of contents

### Appendix C: Example Anterior-Posterior Outer Boundaries with Partial Voluming of Landmark Structures

The protocol definitions of hippocampal subfield boundaries differ in the head, body and tail of the hippocampus. The body anterior and posterior outer boundaries define the range of slices to apply the body inner body protocol. For a detail of the ranging rules, see Appendix B-ix to B-xvi; this supplemental document provides additional examples when the uncus and colliquli landmarks are visualized with partial voluming.

#### Anterior Range

The anterior outer boundary of the hippocampal body is defined as the first slice, posterior to the uncus. When the uncus is visualized with partial voluming, the judgement should be made if the uncus region can be labeled for volume based on the intensity of the region. If determined the region should not be labeled, then the slice is judged to be located posterior to the uncus.

Example: Case 2026

| | | |
|---|---|---|
| 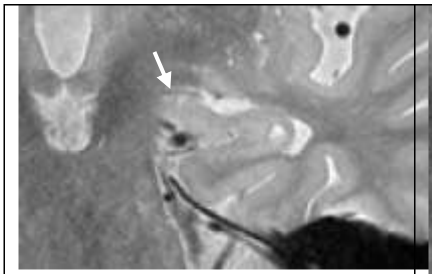 | 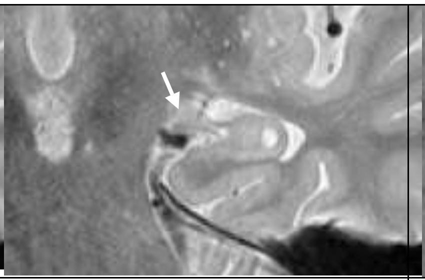 | 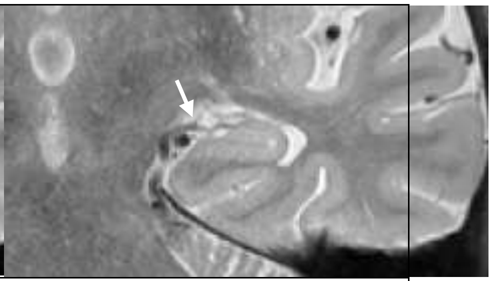 |
| Left, Slice 13: Uncus present | Left, Slice 14: Uncus present (partial voluming would be labeled for subfields using the head protocol) | Left, Slice 15: posterior to uncus (partial voluming is judged insufficient to label) |

### Appendix D: Example Tracing Information and Anterior-Posterior Ranges

To access and download the example images, example tracings in ITK Snap and volumetric data, go to the HSG website: http://www.hippocampalsubfields.com/

#### Anterior-Posterior Ranges

| Scan ID | Left Begin | Left End | Right Begin | Right End |
|---|---|---|---|---|
| 1640 | 17 | 24 | 17 | 24 |
| P41 | 15 | 22 | 14 | 22 |
| 1643 | 13 | 21 | 13 | 21 |
| P47 | 16 | 24 | 16 | 24 |
| ADNI020 | 15 | 21 | 14 | 21 |

Appendix D: Brief Introduction to ITK Snap for Applying Manual Segmentation Protocol

The protocol was designed to be implemented in any software package that supports manual segmentation. For the purpose of the training and initial reliability assessment, we will be using ITKSnap.

To download ITKSnap: http://www.itksnap.org/pmwiki/pmwiki.php (last accessed 10/09/18)

**ITKSnap: Review of Relevant Tools**

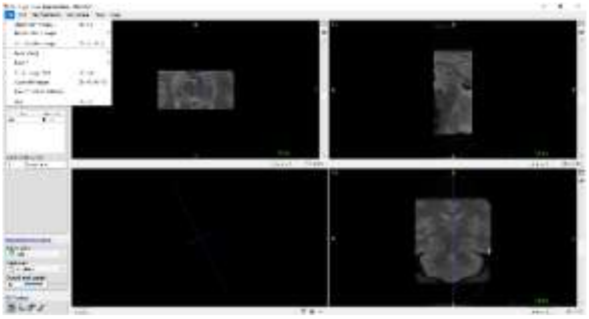

1. To load an image set: go to "Open New Image" under the **File** drop down menu
   a. Load either the DICOM series or NIFTI
   b. Click Finish
2. Create Segmentation Labels by clicking the pallet icon
   a. Load the Hippocampal Subfield Label.txt template by choosing "**Import Label Description**" under **Actions**
3. To Save Segmentation and Export Statistics go to **Segmentation** drop down menu

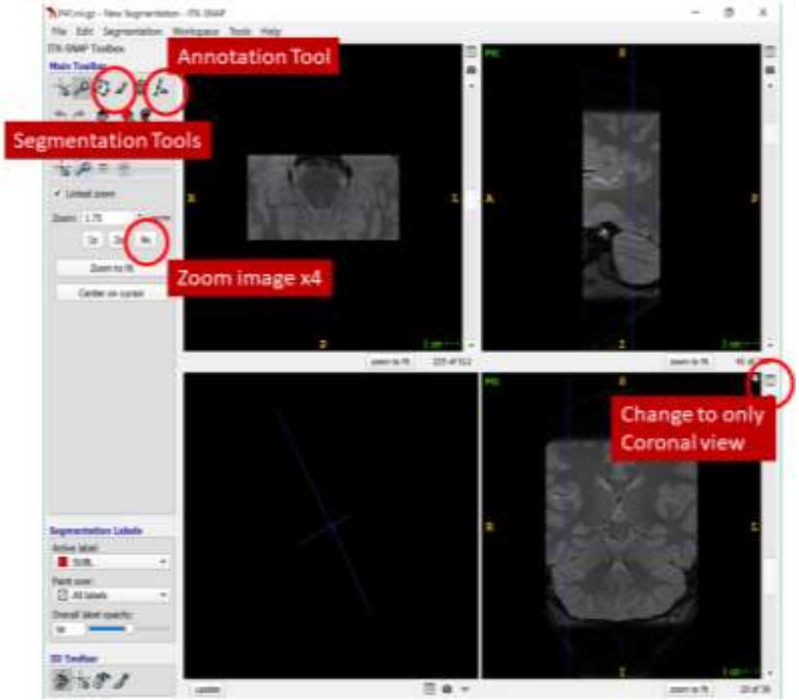

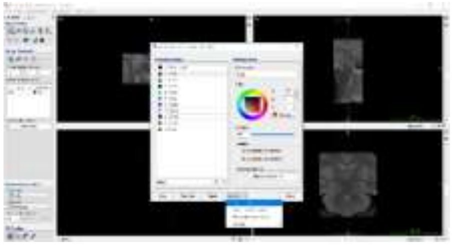

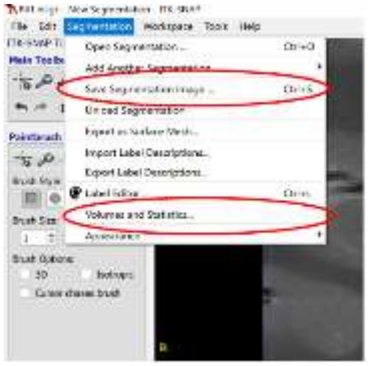
